# On the *independent irritability* of goldfish eggs and embryos – a living communication on the rhythmic yolk contractions in goldfish

**DOI:** 10.1101/2023.11.02.564871

**Authors:** Paul Gerald Layague Sanchez, Chen-Yi Wang, Ing-Jia Li, Kinya G. Ota

## Abstract

Rhythms play an important role in the precise spatiotemporal regulation of biological processes during development and patterning of embryos. We here investigate the rhythmic contractions of the yolk during early development of the goldfish *Carassius auratus*. We quantify these contractions and record robust and persistent rhythmic yolk movements that are not seen in closely-related species (common carp *Cyprinus carpio* and zebrafish *Danio rerio*). We report that yolk contractions are an intrinsic emergent property of the egg, i.e. goldfish eggs are independently irritable / excitable. These contractions do not require sperm entry / fertilization nor cell division, and they notably emerge at a precise time — suggesting that goldfish eggs are able to measure elapsed time from what we infer to be egg activation. We further show that these rhythmic contractions persist even in yolk in isolation. As the yolk itself is known to confer critical cues for early dorsoventral (DV) patterning of teleost embryos, we hypothesize that its contractions in goldfish may influence the patterning process of this species. Indeed, we find that embryos of the naturally more ventralized twin-tail goldfish strain *Oranda* display altered yolk contraction dynamics (i.e. faster contractions). We also present that the period of yolk contractions is independent of *ChdS*, a key gene involved in DV pattening and linked to the twin-tail phenotype, but is instead a trait that is maternal in origin. We aim to uncover whether the yolk contractions happening during early development of domesticated goldfish are the licensing process which permit the emergence of novel patterning phenotypes naturally-observed in this species (e.g. twin-tail and dorsal-finless strains) and which instead have not been found among closely-related species (e.g. common carp) whose yolks do not contract.

This manuscript is here published as a living communication (as described in Gnaiger (2021)). The authors intend to share findings when they are available, encourage feedback and discussion, and invite knowledge exchange and collaboration.

## Introduction

Proper embryonic development and patterning require precise spatiotemporal regulation so that cellular and biochemical events occur at the right place and at the right time. This coordination is highly dynamic and is often mediated by processes that themselves exhibit dynamic behaviors, such as oscillations and waves (Cartwright et al., 2009; Deneke and Di Talia, 2018; Di Talia and Vergassola, 2022; Goodwin and Cohen, 1969; Turing, 1952; Uriu, 2016). The establishment of the animal body plan is a concrete example of the importance of such spatiotemporal regulation (Bénazéraf and Pourquié, 2013; Cooke, 1988; Grimes and Burdine, 2017; Hibi et al., 2018; Meinhardt, 2006): variation in this process and deviations from archetypal development are often lethal or pathological, as is the case of congenital scoliosis (Pourquié, 2011). At the same time, variations in archetypal processes of body plan establishment are e.g. at the origin of the variety of unique phenotypes of the many present-day strains of domesticated goldfish *Carassius auratus* (Abe et al., 2014; Ota and Abe, 2016). One central question in both developmental biology and animal evolution thus remains that of how changes in the spatiotemporal regulation of common developmental programmes can give rise to such a diversity of body forms from a single cell, the egg.

A huge body of research pioneered and inspired by Ernest Everett Just has revealed over the years the dynamic processes involved in- and resulting from-the activation, fertilization, and patterning of the egg (Byrnes and Newman, 2014). E. E. Just advocated for melding of physics and biology in the study of embryonic development (Just, 1939) at a time when this was not common practice, and described the egg as “*self-acting, self-regulating and self-realizing – an independently irritable system*”, i.e. an excitable soft matter (Byrnes and Newman, 2014; Newman, 2009). For instance, the egg exhibits dynamic changes of its cytoskeleton (Just, 1919, 1939; Santella and Chun, 2022), and waves of dynamic calcium signaling during activation and fertilization (Sardet et al., 1998; Stricker, 1999). These dynamics are now known to have the capacity to carry spatiotemporal information that pre-determines later developmental events, e.g. the site of gastrulation in ascidians (Roegiers et al., 1995; Sardet et al., 2007), and are linked to emergent contractile behavior of the egg (Brownlee and Dale, 1990; Ishii and Tani, 2021; Kyozuka et al., 2008; Limatola et al., 2022a) in both ascidian eggs (Brownlee and Dale, 1990; Ishii and Tani, 2021) and in mouse oocytes (Deguchi et al., 2000).

Unlike the egg contractions mentioned above, which are short-lived and occur just after egg activation and fertilization (Brownlee and Dale, 1990; Deguchi et al., 2000; Ishii and Tani, 2021), some eggs and embryos also display rhythmic contractions that last until much later stages of develop-ment. For example, cell-autonomous periodic cortical waves of contraction (PeCoWaCo), linked to the softening of the actomyosin cortex, are evident during the cleavage stages preceding mouse embryo compaction (Maître et al., 2015; Özgüç et al., 2022). Perhaps more strikingly, in goldfish embryos rhythmic contractions of the yolk surface emerge from the 4-cell stage and persist for hours until the epiboly stage (Yamamoto, 1934). These yolk contractions are faster at higher temperatures (Yamamoto, 1934), with temperature constants that cluster with those of processes that are mainly oxidative (Crozier, 1924) and those of others that are primarily involved in growth and development (Crozier, 1926).

The yolk of fish embryos has been shown to be more than just a static nutritional resource. In fact, experiments have shown that the yolk carries determinants for the establishment of the fish body plan, thus playing an active role in development and patterning (Devillers, 1961; Oppenheimer, 1936; Tung et al., 1945; Tung and Tung, 1943, 1944). Specifically, the yolk has been shown to confer dorsal specification cues to developing fish embryos (Mizuno et al., 1999, 1997; Ober and Schulte-Merker, 1999). According to the paradigm defined in zebrafish (*Danio rerio*), shortly after egg activation and fertilization and prior to the first cleavage a parallel array of microtubules forms at the vegetal side of the yolk. This is crucial for the proper asymmetric transport of maternal dorsalizing determinants to the developing embryo’s prospective dorsal organizer (Jesuthasan and Strähle, 1997; Tran et al., 2012). An analogous supply and asymmetrical partitioning of maternal axial determinants has been similarly inferred in goldfish (Mizuno et al., 1997; Tung et al., 1955a; Tung and Tung, 1943).

Interestingly, a wide variety of viable DV patterning phenotypes has been observed in fish. This is the case of the wide variety of median-fin-related morphological phenotypes (i.e. median fin morphotypes) of domesticated goldfish strains, which have been shown to arise from altered DV patterning during embryonic development. While there is evidence for an active role of the acellular yolk in this process, most studies on the matter have focused on underlying genetic and molecular mechanisms. It is now known that twin-tail goldfish carry a mutation in one of their chordin genes (i.e. *ChdS*), which results in a non-functional truncated ChdS protein (Abe et al., 2014). Loss of function of said chordin gene, which naturally inhibits ventralization of the body (Piccolo et al., 1996; Sasai et al., 1994), is sufficient to cause caudal fin bifurcation in wild-type goldfish (Abe et al., 2014; Lee et al., 2023), and this mutation has been also recently implicated in loss of dorsal fin in dorsal finless goldfish strains (Chen et al., 2022). The spatiotemporal dynamics of the contractile goldfish yolk remains understudied in goldfish development and evolution, in particular, and in cell-developmental biology, in general.

Here, we aim to investigate the dynamic, emergent behavior of the goldfish yolk, hoping to learn more about its origin, its maintenance, and its role. In this manuscript, we share findings from our work addressing the following questions:

- On origin and maintenance of yolk contractions: How do the rhythmic yolk contractions emerge? How are they maintained?
- On mechanism and function of yolk contractions: Do the rhythmic yolk contractions have a function? Do they permit the emergence of diverse median fin morphotypes (DV patterning phenotypes) in domesticated goldfish?

## About this version

This is the second version of this manuscript, which we here publish as a living communication (as described in Gnaiger (2021)). We intend to share findings when they are available, encourage feedback and discussion, and invite knowledge exchange and collaboration.

Since the last version, we have substantially increased the sample numbers, prioritizing throughput over image resolution. We have also significantly extended the duration of timelapse imaging to see how the period of the yolk contractions changes over time. To minimize confounding effect of slight differences in environmental conditions, whenever possible, we recovered eggs and embryos from the same clutch and we imaged different conditions and respective controls simultaneously using multi-well dishes. Here, we included some new experiments / quantifications on the onset of yolk contractions in fertilized and unfertilized goldfish eggs, on yolk contractions in dechorionated goldfish embryos, on yolk contractions in bisected embryos separating animal and vegetal poles, on yolk contractions in pharmacological perturbations of actomyosin and calcium, and on yolk contractions in goldfish embryos with different *ChdS* genotypes. We here also avoid discussing about the amplitude and phase of the yolk contractions (which we could not precisely quantify with our simple quantification of yolk contractions) but instead focus on their period. We confirm here the main findings we shared in the first version, except for the period of contractions in embryos treated with microtubule depolymerizing drug, nocodazole. Previously, we reported that the yolk contractions are faster in nocodazole-treated embryos. Our new experiment, with much greater sample numbers, shows an opposite result where the nocodazole-treated samples are actually slower than controls.

In this version, we also cited more older references (including those in non-English languages) to refer to earlier work that investigated the rhythmic movements of the yolk in different teleost fishes. With this, we invite the reader to explore a rich body of literature on the topic, which piqued the interest of many for some time but since then has remained understudied.

## Other versions

First version: 10.1101/2023.11.02.564871v1

## Results

### Extracting the mean pixel value of an egg / embryo over time allows quantification of the period of yolk contractions from simple timelapse stereomicroscopy

We imaged developing goldfish embryos under the stereomicroscope and observed rhythmic yolk contractions starting at the 4-cell stage and persisting until later stages (Supplementary Movie M1), as reported by Yamamoto (1934). To quantify these contractions, we segmented the yolk and determined its circularity, its perimeter, and its projected area over time. However, we immediately realized that quantifying yolk contractions from measurements of the shape of the yolk requires precise image segmentation (with the yolk as foreground) over long periods of time (spanning > 12 hours post-fertilization, from cleavage to epiboly stages). This is challenging in our simple experimental and imaging conditions as the yolk has low contrast compared to the background and to the ooplasm / embryo proper, the yolk changes its shape and position in 3D, and the entire egg / embryo is not fixed in one axis. Accordingly, we also extracted the timeseries from the mean pixel value of the entire egg / embryo, circumscribed by its chorion, over time (Figure 1, Supplementary Movie M2), reasoning that still-frames of an egg / embryo after a full contraction of the yolk would look most similar, and hence would have most similar mean pixel values (Figure 1 1 and 3, 2 and 4, 5 and 8). For the analysis, we detrended the extracted timeseries via sinc-filter detrending after specifying a cut-off period. We subjected the detrended timeseries to continuous wavelet transform using the Morlet wavelet as mother wavelet (Mönke et al., 2020), allowing us to recover the instantaneous period along a ridge tracing wavelet with maximum power for every timepoint. Comparing the timeseries and the evolution of period over time, we noted that the mean pixel value of the embryo is a proxy of the projected area of the segmented yolk. However, as it does not rely on image segmentation, extracting the timeseries of yolk contractions from the mean pixel value of the entire egg / embryo is additionally more robust, at least in quantifying the period of contractions (which does not rely on sample orientation, unlike quantifying phase and amplitude), compared to measurements of the perimeter, circularity, or projected area of the yolk when some segmentation is possible. As exemplified in Figure 1, the timeseries extracted from the mean pixel value provides a suitable approximation of the period of yolk contractions, matching the actual period of a full contraction (e.g. Figure 1, timepoints 1 and 3, or 2 and 4, or 5 and 8) determined from manual inspection of timelapse imaging (i.e. the ground truth). Thus, the method from mean pixel value provides a simple and flexible measure of the yolk contractions, and we henceforth used this method in our quantification.

**Fig. 1.**
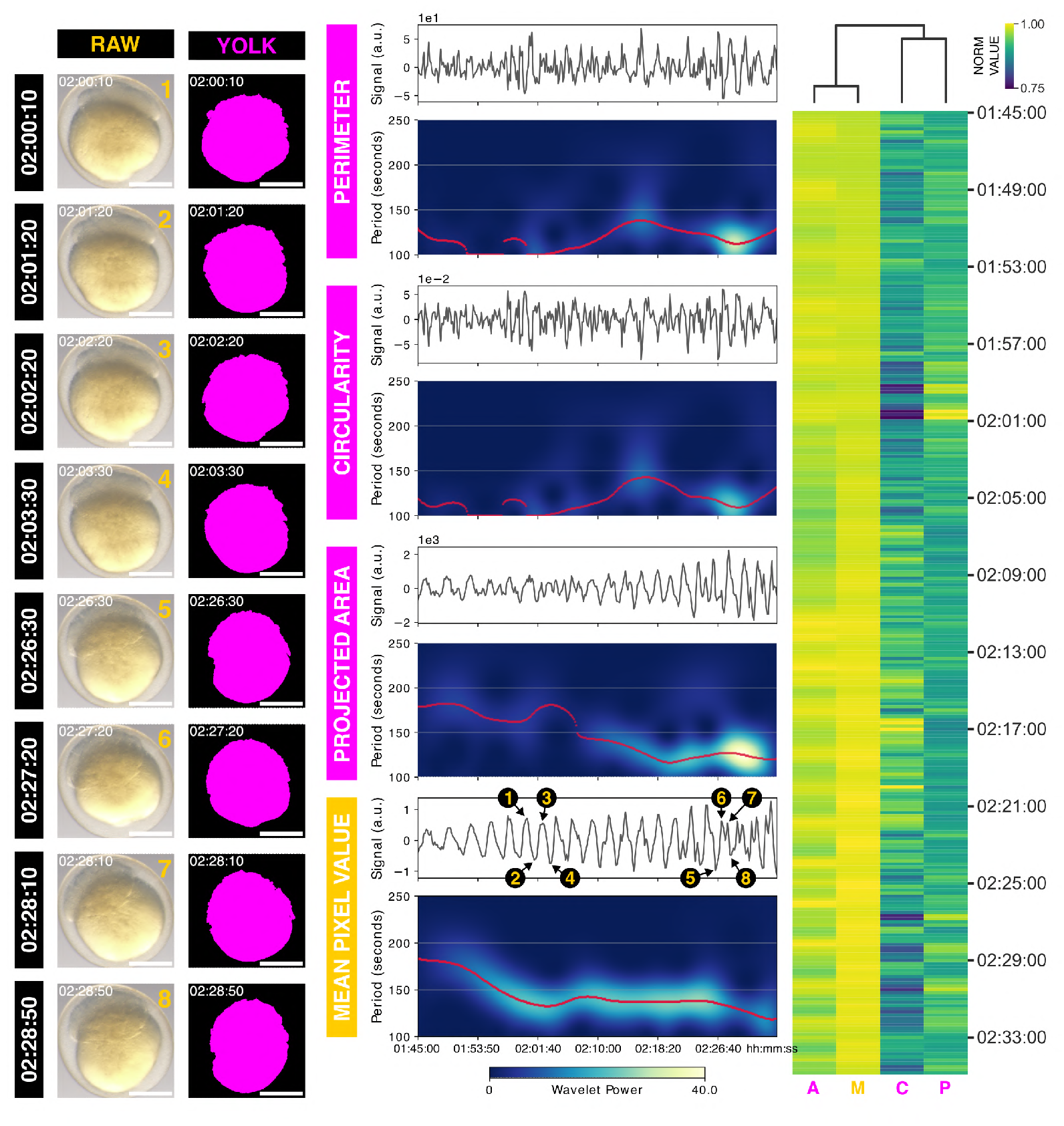
Extracting the mean pixel value of an embryo over time allows quantification of yolk contractions from simple timelapse stereomicroscopy. (**Left**) Snapshots of a goldfish embryo and its yolk (magenta, after image segmentation) over time (hh:mm:ss post-fertilization). Scale bar = 500 *µ*m. (**Middle**) Timeseries of yolk perimeter, yolk circularity, yolk projected area, and embryo mean pixel value over time after sinc-filter detrending with cut-off period = 250 seconds, alongside instantaneous period after subjecting the detrended timeseries to continuous wavelet transform. Red marks the ridge tracing the maximum wavelet power for each timepoint. A high power (power > 3) indicates strong correlation of the wavelet with the signal versus white noise. Numbers (1–8) mark corresponding timepoints of snapshots shown in the left panel. (**Right**) Clustering of normalized timeseries of circularity of the yolk (C), perimeter of the yolk (P), projected area of the yolk (A), and mean pixel value of the embryo (M). Time is in hh:mm:ss post-fertilization. Raw timeseries are shown alongside the timelapse of the goldfish embryo and its yolk (after image segmentation) in Supplementary Movie M2.

**Fig. 2.**
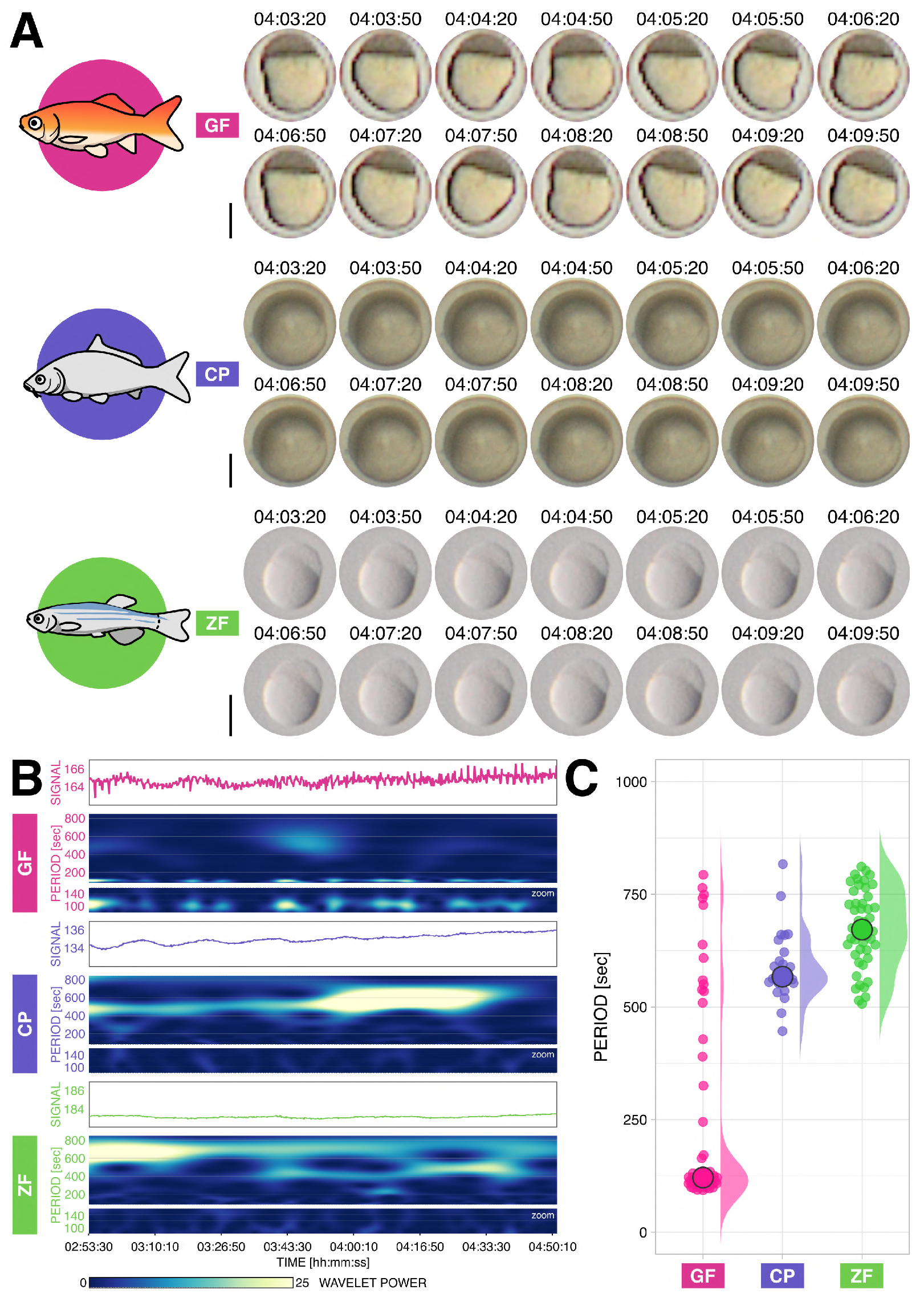
Goldfish exhibit persistent yolk contractions during embryonic development, unlike closely-related common carp and zebrafish. (**A**) Snapshots of goldfish *Carassius auratus* (GF), common carp *Cyprinus carpio* (CP), and zebrafish *Danio rerio* (ZF) embryos for a subset of imaging time, at 04:03:20–04:09:50 hh:mm:ss post-fertilization. Scale bar = 500 *µ*m. (**B**) Raw timeseries of representative fish embryos and corresponding instantaneous period of contractions (if present) after continuous wavelet transform of detrended timeseries (via sinc-filter detrending, cut-off period = 850 seconds). Shown is at 02:53:30–04:51:20 hh:mm:ss post-fertilization, a subset of the entire imaging time where all conditions show median wavelet power > 3, i.e. greater than the minimum wavelet power corresponding to 95% confidence interval in case of white noise. Samples are the same as those in panel A. Note high wavelet power for short period (fast) rhythm in goldfish, which is not present in common carp nor in zebrafish. Also note long period (slow) rhythmic trend present in the timeseries of goldfish. (**C**) Median period of goldfish (GF, n = 61), common carp (CP, n = 25), and zebrafish (ZF, n = 40) embryos at 02:53:30–04:51:20 hh:mm:ss post-fertilization. Individual samples are represented as color-coded dots, while the median of all the samples is denoted as a color-coded circle with black margin. The distribution of the data is also plotted. These data are summarized in Table 1. For temporal evolution of the period and of the wavelet power for the entire duration of the experiment, see Supplementary Figure F1A and Supplementary Figure F2A, respectively. Timelapse of some of the fish embryos are shown in Supplementary Movie M3.

**Fig. 3.**
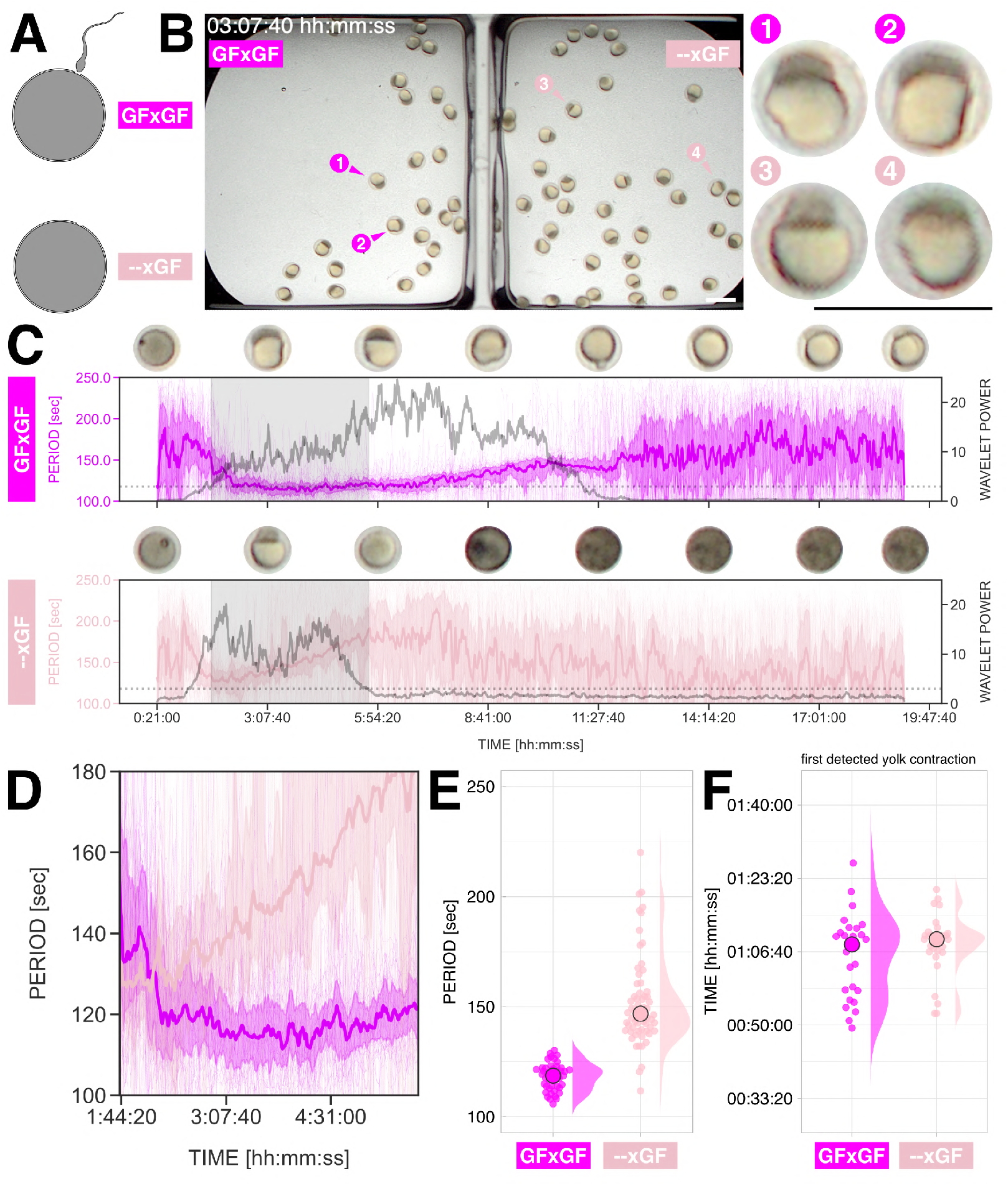
The rhythmic contractions of the goldfish yolk do not require fertilization and emerge at a precise time. (**A**) Schematic of experiment comparing fertilized (GFxGF, magenta) and unfertilized (–xGF, light pink) goldfish eggs. (**B**) Snapshot of simultaneous imaging of fertilized (GFxGF, left) and unfertilized (–xGF, right) goldfish eggs at 03:07:40 hh:mm:ss post-exposure of egg (and sperm, for fertilized sample) to water. Representative fertilized (1 and 2, magenta arrowheads) and unfertilized (3 and 4, light pink arrowheads) samples are magnified on the right. Scale bar = 2 mm. Simultaneous timelapse of fertilized and unfertilized goldfish eggs are shown in Supplementary MovieM4. (**C**) Temporal evolution of period of yolk contractions for the entire duration of the experiment, obtained from wavelet analysis of detrended timeseries (via sincfilter detrending, cut-off period = 250 seconds). The period evolution for each sample and the median of the periods are represented as a color-coded dashed line and a color-coded solid line, respectively. The color-coded shaded area corresponds to the interquartile range. Snapshots of representative samples (samples 2 and 3 in panel B for fertilized and unfertilized conditions, respectively) are shown above each plot. Median wavelet power is also plotted in gray, with a horizontal dotted gray line marking wavelet power threshold = 3, which corresponds to 95% confidence interval in case of white noise. Shaded region corresponds to subset time duration considered for comparison of periods, i.e. 01:43:10–05:40:30 hh:mm:ss post-exposure of egg (and sperm, for fertilized sample) to water, where both conditions show median wavelet power > 3. (**D**) Temporal evolution of period of yolk contractions in fertilized (magenta) and unfertilized (light pink) goldfish eggs for the specified subset of imaging time. The period evolution for each sample and the median of the periods are represented as a color-coded dashed line and a color-coded solid line, respectively. The color-coded shaded area corresponds to the interquartile range. (**E**) Median period of yolk contractions in fertilized (GFxGF, n = 45) and unfertilized (–xGF, n = 62) eggs at 01:43:10–05:40:30 hh:mm:ss post-exposure of egg (and sperm, for fertilized sample) to water. Individual samples are represented as color-coded dots, while the median of all the samples is denoted as a color-coded circle with black margin. The distribution of the data is also plotted. These data are summarized in Table2. (**F**) Onset of yolk contractions in fertilized (GFxGF, n = 27) and unfertilized (–xGF, n = 30) eggs. Individual samples are represented as color-coded dots, while the median of all the samples is denoted as a color-coded circle with black margin. The distribution of the data is also plotted. Time is in hh:mm:ss post-exposure of egg (and sperm, for fertilized sample) to water. These data are summarized in Table3.

**Fig. 4.**
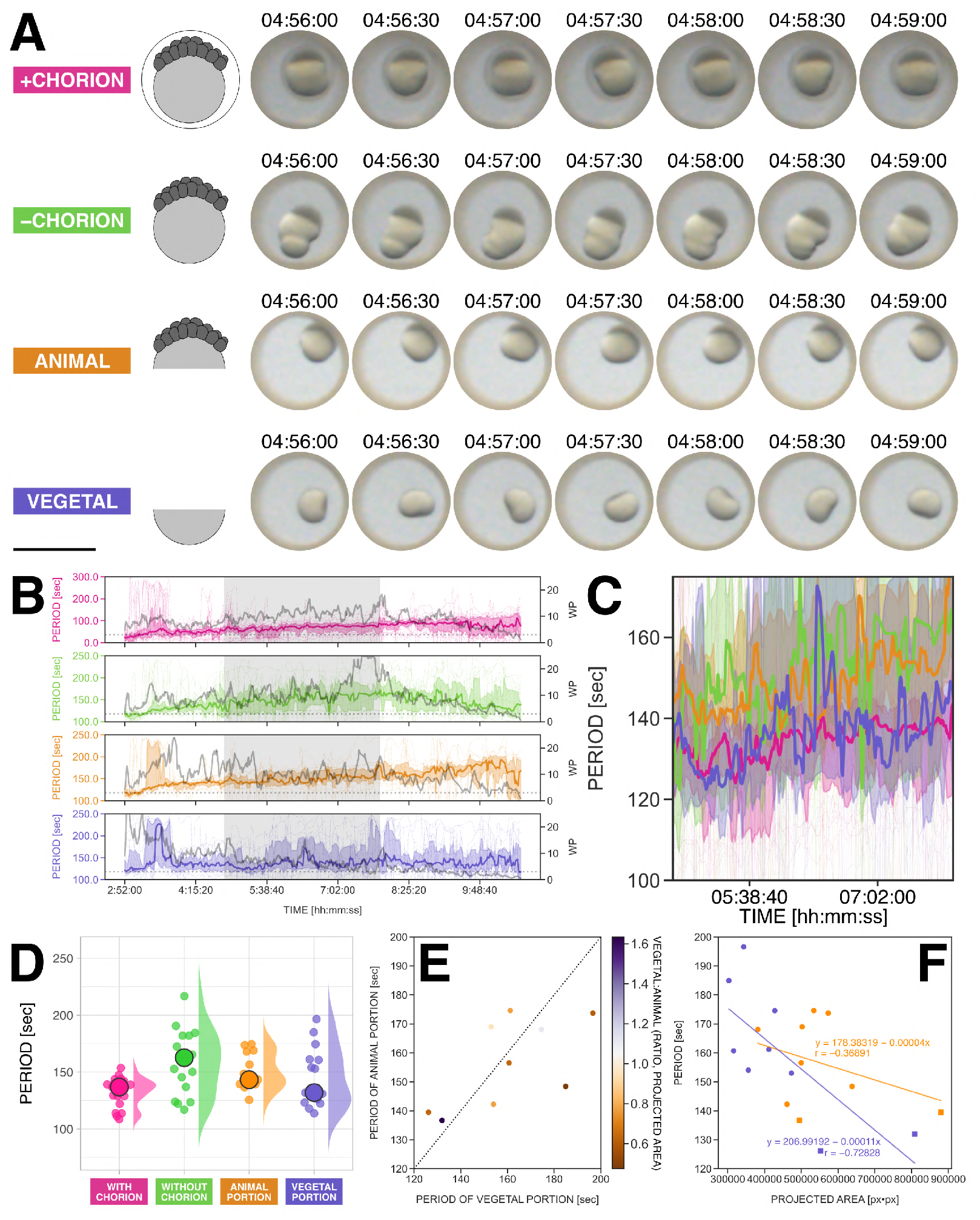
The rhythmic contractions are an emergent property of the goldfish yolk, persisting even after dechorionation and equatorial bisection. (**A**) Snapshots of an intact goldfish embryo with chorion (+CHORION, pink), an intact goldfish embryo without chorion (–CHORION, green), the animal portion of a bisected goldfish embryo (ANIMAL, orange), and the vegetal portion of same bisected goldfish embryo (VEGETAL, purple) for a subset of imaging time, at 04:56:00–04:59:00 hh:mm:ss post-fertilization. Scale bar = 2 mm. Simultaneous timelapse of multiple +CHORION and –CHORION samples are shown in Supplementary Movie M5. Bisection of a goldfish embryo to isolate the animal and vegetal portions is shown in Supplementary Movie M6. Simultaneous timelapse of the different conditions and detrended timeseries of representative samples in this figure are shown in Supplementary Movie M7. (**B**) Temporal evolution of period of yolk contractions for the entire duration of the experiment, obtained from wavelet analysis of detrended timeseries (via sinc-filter detrending, cut-off period = 250 seconds). The period evolution for each sample and the median of the periods are represented as a color-coded dashed line and a color-coded solid line, respectively. The color-coded shaded area corresponds to the interquartile range. Median wavelet power (WP) is also plotted in gray, with a horizontal dotted gray line marking wavelet power threshold = 3, which corresponds to 95% confidence interval in case of white noise. Shaded region corresponds to subset time duration considered for comparison of periods, i.e. 04:49:00–07:51:00 hh:mm:ss post-fertilization. From top to bottom: intact goldfish embryos with chorion (pink), intact goldfish embryos without chorion (green), animal portions of bisected goldfish embryos (orange), and vegetal portions of bisected goldfish embryos (purple). (**C**) Temporal evolution of period of yolk contractions in intact goldfish embryos with chorion (pink), intact goldfish embryos without chorion (green), animal portions of bisected goldfish embryos (orange), and vegetal portions of bisected goldfish embryos (purple) for the specified subset of imaging time. The period evolution for each sample and the median of the periods are represented as a color-coded dashed line and a color-coded solid line, respectively. The color-coded shaded area corresponds to the interquartile range. (**D**) Median period of yolk contractions in intact goldfish embryos with chorion (WITH CHORION, n = 23), intact goldfish embryos without chorion (WITHOUT CHORION, n = 17), animal portions of bisected goldfish embryos (ANIMAL PORTION, n = 13), and vegetal portions of bisected goldfish embryos (VEGETAL PORTION, n = 15) at 04:49:00–07:51:00 hh:mm:ss post-fertilization. Individual samples are represented as color-coded dots, while the median of all the samples is denoted as a color-coded circle with black margin. The distribution of the data is also plotted. These data are summarized in Table 4. (**E**) Period of yolk contraction in animal portion of bisected goldfish embryo plotted against the period of yolk contraction in its corresponding vegetal partner. Each dot corresponds to animal-vegetal partners (i.e. animal and vegetal portions recovered from the same bisected goldfish embryo). Colors correspond to the ratio of the projected areas of the two portions. More orange (or purple) means that the animal (or vegetal) portion is bigger than the vegetal (or animal) portion. Dotted line marks the diagonal where the periods of yolk contractions in animal-vegetal partners are equal. (**F**). Period of yolk contraction in animal (orange) and vegetal (purple) portions plotted against their projected area, with corresponding color-coded results from linear regression analysis.

**Fig. 5.**
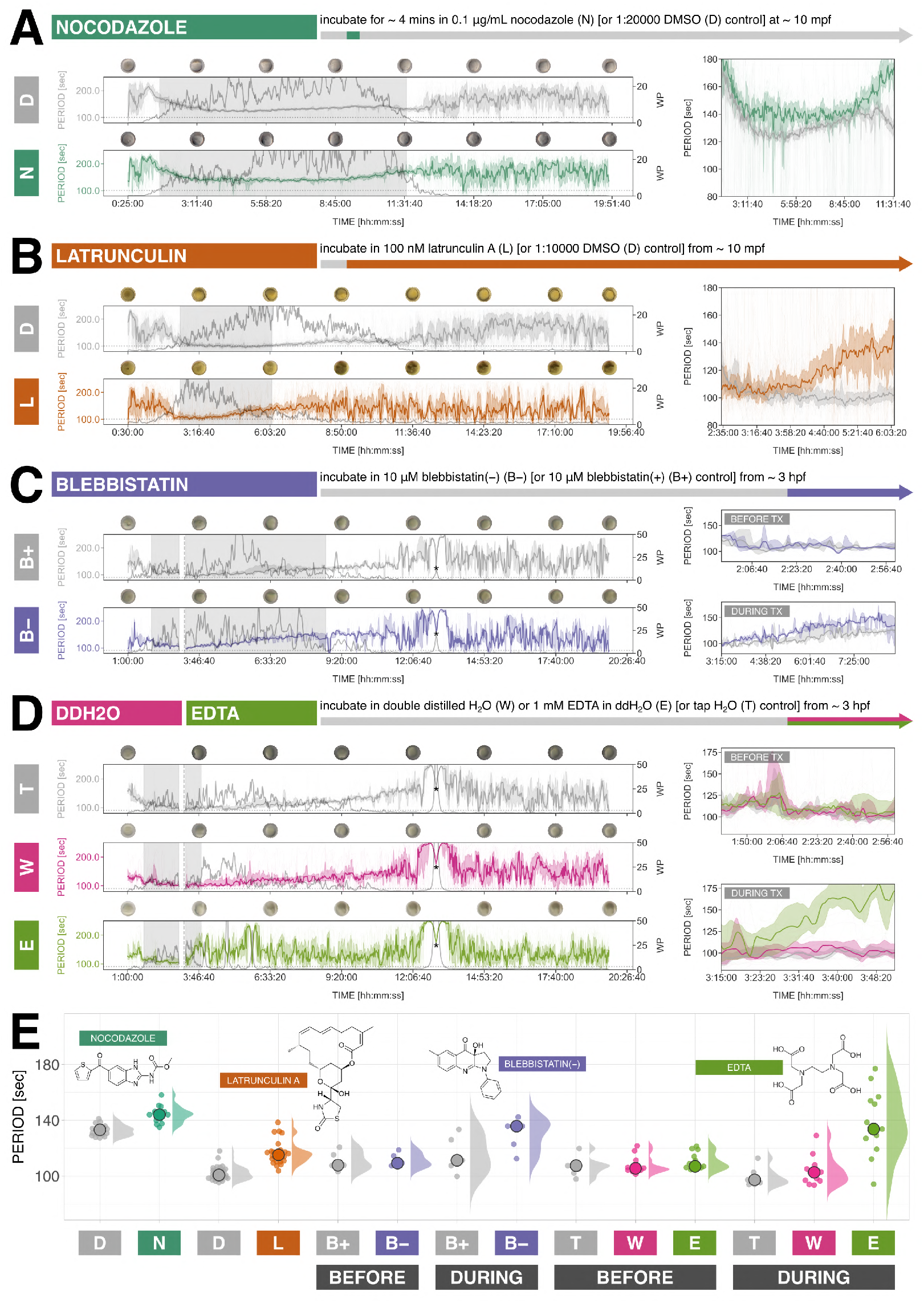
Rhythmic contractions of the yolk in goldfish embryos exposed to pharmacological perturbation of microtubules, actomyosin, and calcium. (**A**) To perturb microtubule polymerization prior to the first cleavage, goldfish embryos were incubated in either 0.1 *µ*g/mL nocodazole (N, dark green, a microtubule-depolymerizing drug) or 1:20000 DMSO control (D, light gray) for *∼* 4 mins at *∼* 10 mins post-fertilization. Simultaneous timelapse of nocodazole-treated goldfish embryos and controls is shown in Supplementary MovieM8. (**B**) To perturb actin, goldfish embryos were incubated in either 100 nM latrunculin A (L, orange, an actin polymerization inhibitor) or 1:10000 DMSO control (D, light gray) from *∼* 10 mins post-fertilization. Simultaneous timelapse of latrunculin-treated goldfish embryos and controls is shown in Supplementary Movie M10. (**C**) To perturb myosin, goldfish embryos were incubated in either 10 *µ*M blebbistatin(–) (B–, purple, a myosin inhibitor) or 10 *µ*M blebbistatin(+) control (B+, light gray) from *∼* 3 hours post-fertilization. Due to unavailability of spawning wild-type (ZWJ-*ChdS*(+/+) strain) goldfish, ZWJ-*ChdS*(–/–)-singletail goldfish was used. Simultaneous timelapse of blebbistatin(–)-treated goldfish embryos and controls, before and during treatment, is shown in Supplementary Movie M11. To perturb calcium, goldfish embryos were incubated in either double distilled water (ddH_2_ O, W, pink, Ca^2+^ free water), 1 mM EDTA in ddH_2_ O (E, light green, a Ca^2+^ chelating agent), or tap water control (T, light gray) from *∼* 3 hours post-fertilization. Due to unavailability of spawning wild-type (ZWJ-*ChdS*(+/+) strain) goldfish, ZWJ-*ChdS*(– /–)-singletail goldfish was used. Simultaneous timelapse of ddH_2_ O/EDTA-treated goldfish embryos and controls, before and during treatment, is shown in Supplementary Movie M12. For panels **A**–**D**: Left: Temporal evolution of period of yolk contractions for the entire duration of the experiment, obtained from wavelet analysis of detrended timeseries (via sinc-filter detrending, cut-off period = 250 seconds). The period evolution for each sample and the median of the periods are represented as a color-coded dashed line and a color-coded solid line, respectively. The color-coded shaded area corresponds to the interquartile range. Snapshots of representative samples are shown above each plot. Median wavelet power (WP) is also plotted in gray, with a horizontal dotted gray line marking wavelet power threshold = 3, which corresponds to 95% confidence interval in case of white noise. Shaded region corresponds to subset time duration considered for comparison of periods, where the conditions show median wavelet power > 3. Asterisks mark timepoint where ambient light was briefly disturbed during the timelapse imaging. For panels **A**–**B**: Right: Temporal evolution of period of yolk contractions in drug-treated goldfish embryos (color-coded) and their respective controls (light gray) for the specified subset of imaging time. The period evolution for each sample and the median of the periods are represented as a color-coded dashed line and a color-coded solid line, respectively. The color-coded shaded area corresponds to the interquartile range. For panels **C**–**D**: Right: Temporal evolution of period of yolk contractions in drug-treated goldfish embryos (color-coded) and their respective controls (light gray) for the specified subset of imaging time, until 03:00:00 hh:mm:ss post-fertilization (top, before treatment) and from 03:15:00 hh:mm:ss post-fertilization (bottom, during treatment). The period evolution for each sample and the median of the periods are represented as a color-coded dashed line and a color-coded solid line, respectively. The color-coded shaded area corresponds to the interquartile range. (**E**) Median period of yolk contractions in drug-treated goldfish embryos (color-coded) and their respective controls (light gray) at specified subset of imaging time. Individual samples are represented as color-coded dots, while the median of all the samples is denoted as a color-coded circle with black margin. The distribution of the data is also plotted. These data are summarized in Table 5 for the nocodazole experiment, Table 6 for the latrunculin experiment, Table 7 for the blebbistatin(–) experiment, and Table 8 for the ddH_2_ O and EDTA experiment. The phenotypes of drug-treated goldfish embryos and their respective controls at 1 day post-fertilization are shown in Supplementary Figure F3.

### Goldfish, unlike closely-related common carp and zebrafish, exhibit persistent rhythmic yolk contractions during embryonic development

Having established a pipeline for the quantification and analysis of yolk contractions from timelapse images, we compared goldfish yolk dynamics with those of a closely-related species, common carp (*Cyprinus carpio*), and of another cypriniform species, zebrafish (*Danio rerio*). Persistent and periodic yolk contractions were evident in goldfish embryos, but not in common carp nor in zebrafish (Figure 2A, Supplementary Movie M3). As summarized in Table 1 and shown in Figure 2B-C (and Supplementary Figure F1A), these contractions had a stable period of 121.31*±*14.98 seconds (113.23–123.33 seconds) (n = 61) or ∼2 mins at 02:53:30–04:51:20 hh:mm:ss post-fertilization). In common carp and zebrafish embryos, we instead only noted longer-period (i.e. slower) rhythms with period of 567.11 ± 31.86 seconds (555.00–601.66 seconds) or ∼ 9.5 mins for common carp (n = 25) and 671.51 ± 60.59 seconds (643.98–717.61 seconds) or ∼ 11.2 mins for zebrafish (n = 46) at 02:53:30–04:51:20 hh:mm:ss post-fertilization.

**Table 1.**
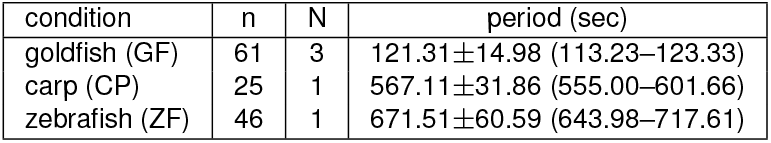
Period of detected rhythms in goldfish *Carassius auratus* (GF), common carp *Cyprinus carpio* (CP), and zebrafish *Danio rerio* (ZF) at 02:53:30–04:51:20 hh:mm:ss post-fertilization, expressed as median *±* median absolute deviation (95% confidence interval of the median). n and N correspond to the number of samples and to the number of clutches, respectively.

**Table 2.** Period of yolk contractions in fertilized and unfertilized goldfish eggs at 01:43:10–05:40:30 hh:mm:ss post-exposure of egg (and sperm, for fertilized sample) to water, expressed as median *±* median absolute deviation (95% confidence interval of the median). n and N correspond to the number of samples and to the number of clutches, respectively.

| condition | n | N | period (sec) |
| --- | --- | --- | --- |
| fertilized | 45 | 2 | 118.66±4.12 (116.23–120.73) |
| unfertilized | 62 | 2 | 146.82±7.83 (142.81–152.16) |

**Table 3.** Onset of yolk contractions in fertilized and unfertilized goldfish eggs, expressed as median *±* median absolute deviation (95% confidence interval of the median) time when the first yolk contraction was detected. Time is in hh:mm:ss post-exposure of egg (and sperm, for fertilized sample) to water. n and N correspond to the number of samples and to the number of clutches, respectively.

| condition | n | N | time (hh:mm:ss) |
| --- | --- | --- | --- |
| fertilized | 27 | 2 | 01:08:20±00:05:10 (00:58:40–01:10:30) |
| unfertilized | 30 | 2 | 01:09:30±00:02:20 (01:07:30–01:10:20) |

**Table 4.** Period of yolk contractions in intact goldfish embryos with chorion, intact goldfish embryos without chorion (dechorionated), animal portions of bisected goldfish embryos, and vegetal portions of bisected goldfish embryos at 04:49:00– 07:51:00 hh:mm:ss post-fertilization, expressed as median *±* median absolute deviation (95% confidence interval of the median). n and N correspond to the number of samples and to the number of clutches, respectively.

| condition | n | N | period (sec) |
| --- | --- | --- | --- |
| with chorion | 23 | 3 | 136.84±06.73 (129.52–139.14) |
| without chorion | 17 | 3 | 167.67±19.39 (137.00–182.02) |
| animal portion | 13 | 3 | 143.34±06.64 (139.50–168.05) |
| vegetal portion | 15 | 3 | 131.97±18.30 (123.23–161.20) |

**Table 5.**
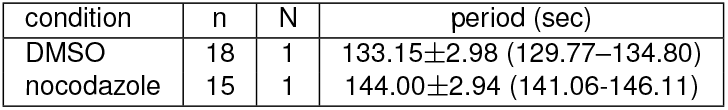
Period of yolk contractions in nocodazole-treated goldfish embryos and their respective controls at 01:41:20–11:36:10 hh:mm:ss post-fertilization, expressed as median *±* median absolute deviation (95% confidence interval of the median). Goldfish embryos were incubated in either 0.1 *µ*g/mL nocodazole or 1:20000 DMSO control for *∼* 4 mins at *∼* 10 mins post-fertilization. n and N correspond to the number of samples and to the number of clutches, respectively.

| condition | $n$ | $N$ | period (sec) |
| --- | --- | --- | --- |
| DMSO | 18 | 1 | 133.15±2.98 (129.77–134.80) |
| nocodazole | 15 | 1 | 144.00±2.94 (141.06–146.11) |

**Table 6.** Period of yolk contractions in latrunculin-treated goldfish embryos and their respective controls at 02:32:40–06:07:20 hh:mm:ss post-fertilization, expressed as median *±* median absolute deviation (95% confidence interval of the median). Goldfish embryos were incubated in either 100 nM latrunculin A or 1:10000 DMSO control from *∼* 10 mins post-fertilization. n and N correspond to the number of samples and to the number of clutches, respectively.

| condition | $n$ | $N$ | period (sec) |
| --- | --- | --- | --- |
| DMSO | 29 | 1 | 100.73±2.90 (99.07–104.05) |
| latrunculin A | 26 | 1 | 115.19±3.14 (112.43–117.12) |

**Table 7.** Period of yolk contractions in blebbistatin(–)-treated goldfish embryos and their respective controls at 01:55:20–03:00:00 hh:mm:ss (before treatment) and 03:15:00–08:42:30 hh:mm:ss (during treatment) post-fertilization, expressed as median *±* median absolute deviation (95% confidence interval of the median). Goldfish embryos were incubated in either 10 *µ*M blebbistatin(–) or 10 *µ*M blebbistatin(+) control from *∼* 3 hours post-fertilization. Due to unavailability of spawning wild-type (ZWJ-*ChdS*(+/+) strain) goldfish, ZWJ-*ChdS*(–/–)-singletail goldfish was used. n and N correspond to the number of samples and to the number of clutches, respectively.

| condition | $n$ | $N$ | period (sec) |
| --- | --- | --- | --- |
| before, blebbistatin(+) | 5 | 1 | 107.77±04.60 (103.16–120.66) |
| before, blebbistatin(–) | 5 | 1 | 109.25±02.47 (106.78–118.68) |
| during, blebbistatin(+) | 5 | 1 | 111.28±11.52 (99.77–133.44) |
| during, blebbistatin(–) | 5 | 1 | 135.73±06.55 (112.44–142.28) |

**Table 8.** Period of yolk contractions in goldfish embryos treated with either double distilled water (ddH_2_ O) or EDTA and their respective controls at 01:37:50–03:00:00 hh:mm:ss (before treatment) and 03:15:00–03:52:20 hh:mm:ss (during treatment) post-fertilization, expressed as median *±* median absolute deviation (95% confidence interval of the median). Goldfish embryos were incubated in either ddH_2_ O, 1 mM EDTA in ddH_2_ O, or tap H_2_ O control from *∼* 3 hours post-fertilization. Due to unavailability of spawning wild-type (ZWJ-*ChdS*(+/+) strain) goldfish, ZWJ-*ChdS*(–/–)-singletail goldfish was used. n and N correspond to the number of samples and to the number of clutches, respectively.

| condition | $n$ | $N$ | period (sec) |
| --- | --- | --- | --- |
| before, tap H<sub>2</sub>O | 7 | 1 | 107.44±02.42 (102.19–108.34) |
| before, ddH<sub>2</sub>O | 11 | 1 | 105.53±01.20 (104.33–111.34) |
| before, EDTA | 13 | 1 | 106.94±02.31 (106.09–113.98) |
| during, tap H<sub>2</sub>O | 7 | 1 | 097.28±03.34 (093.95–102.67) |
| during, ddH<sub>2</sub>O | 11 | 1 | 102.66±05.72 (096.55–107.93) |
| during, EDTA | 13 | 1 | 133.70±17.98 (126.95–153.98) |

The rhythms we had detected for the three fish species had high wavelet powers (Supplementary Figure F2A), correlating well with considered wavelets and not with white noise (Mönke et al., 2020) at least at specific timepoints during development (see differences in temporal profile of wavelet power between the three fish species as represented by black bars in Supplementary Figure F1A). Comparison with timeseries from background regions (in close proximity to analyzed fish embryos) verified that the detected embryo rhythms were not an artefact of the image acquisition nor were due to background effects (Figure 2B versus Supplementary Figure F2B). Interestingly, we also observed a trend in the timeseries of goldfish embryos (see for e.g. raw timeseries of representative goldfish sample in Figure 2B) seemingly matching the slow (long-period) rhythms detected in common carp and zebrafish. For some goldfish samples (e.g. 15/61 goldfish samples plotted in Figure 2C), this slow rhythm could even register higher wavelet power than the faster yolk contractions when the cut-off period was set at 850 seconds. We are still not certain what these slow rhythms are. Nonetheless, such rhythms are clearly not the same as the persistent yolk contractions seen in goldfish. Notably, we also recorded rhythmic movements near the animal pole of common carp embryos (Supplementary Movie M3), which was present shortly after fertilization (Supplementary Figure F1B) and lasted only until just before the first cleavage was completed (Supplementary Figure F1C). This indicates that common carp samples are deformable at least up to some point and yet somehow do not show goldfish-like persistent rhythmic yolk contractions. In summary, we find that goldfish embryos exhibit unique dynamics in the form of persistent rhythmic yolk contractions that are not seen in embryos of common carp and zebrafish.

### The rhythmic contraction of the goldfish yolk is independent from fertilization or cell division, and emerge at a precise time

We then investigated whether fertilization of the egg is required to trigger rhythmic yolk contractions in goldfish. To address this question, we recovered goldfish eggs from one clutch but exposed only some to goldfish sperm (Figure 3A). To minimize any effect from slight differences in environmental / imaging conditions, we imaged both conditions simultaneously using a multi-well dish (Figure 3B). We also extended the duration of timelapse imaging to see how the period of the contractions (if present) changed over time. For fertilized samples, we noted rhythmic yolk contractions from the 4-cell stage which became fairly stable during the early blastula stages but then gradually slowed down before ceasing towards the end of epiboly (Figure 3C). Unfertilized samples also notably exhibited rhythmic yolk contractions that lasted for many hours but slowed down and ceased more quickly than fertilized controls (Figure 3C). This corresponded with a slower period of yolk contractions in unfertilized eggs (Table 2, Figure 3D-E), possibly reflecting loss of viability in these samples.

Remarkably, as actually previously reported by Yamamoto (1954), unfertilized goldfish eggs not only showed rhythmic yolk contractions; these contractions also emerged at around the same time as they did in fertilized eggs, which started at ∼4-cell stage (Table 3, Figure 3F, Supplementary Movie M4). Considering there is no real “4-cell” stage in unfertilized samples, as they fail to undergo cell divisions (Supplementary Movie M4) (Yamamoto, 1954), these results hint to an intrinsic timing mechanism to the beginning of yolk contractions relative to a certain reference “time 0” that could be egg activation, which in turn is triggered by contact with water (Lee et al., 1999; Yamamoto, 1954). These data reveal that the contractions of the yolk do not require sperm entry / fertilization, and that they are also not due to cell division, further supporting the idea that the goldfish egg is an independently irritable system, i.e. it is excitable (Newman, 2009). Upon passing a threshold, possibly initiated at egg activation, goldfish eggs are able to exhibit emergent contractile dynamics that persist for a long time.

### The rhythmic contractions are an emergent property of the goldfish yolk, persisting even after dechorionation and equatorial bisection

We wondered if the yolk contraction is a response to mechanical constraints set by the chorion and if either the animal pole or the vegetal pole is necessary for its maintenance. To address this, we dechorionated goldfish embryos and bisected some of them at the equator from the 32-cell stage, separating the animal (i.e. the embryo proper and some yolk) and the vegetal (i.e. the isolated yolk) portions. We noted that rhythmic yolk contractions persisted in all conditions considered (Figure 4A, Supplementary Movie M5, Supplementary Movie M6, Supplementary Movie M7). We recorded more apparent yolk deformations in samples lacking a chorion than in controls, implying that the chorion is actually constraining yolk contractions as noted before by Wülker (1953) in pike and trout whose yolks also contract. For surgically manipulated samples, we observed persistent yolk contractions in both animal and vegetal portions derived from equatorial bisection of goldfish embryos, suggesting that the maintenance of these contractions do not rely on either the animal or the vegetal poles. These data indicate that the rhythmic contractions are an emergent property of the goldfish yolk, independent of the chorion and importantly independent of the embryo proper. Interestingly, we recorded apparent rotations of isolated yolk samples (i.e. vegetal portion), which we did not observe in samples that contain the embryo proper, possibly revealing intrinsic dynamics of the yolk which is instead “anchored” by the embryo proper in intact samples. The period of yolk contractions between the different conditions showed no clear difference and was within sample variability, i.e. with overlapping 95% confidence intervals of the median (Table 4, Figure 4B-D). Without the chorion, however, the rhythmic yolk contractions tended to have slower periods, possibly reflecting longer time for a full contraction of a higher-order deformation than for a full contraction inside spatial restrictions of the chorion.

There was no clear match in period between animal-vegetal partners, i.e. animal and vegetal portions recovered from the same bisected goldfish embryo (Figure 4E). However, we saw a negative correlation between size (i.e. projected area) of the sample and the period of rhythmic contractions, i.e. the smaller the sample, the longer the period, especially for the vegetal portions (Figure 4F). That is, we observe a re-lationship opposite to what might be expected if there was scaling between sample size and yolk contraction period. As we observed peculiar dynamics in these samples (e.g. apparent rotations in vegetal portions), more sophisticated imaging modalities (e.g. to image in 3D) and quantification methods (e.g. to capture shape, area, and volume changes) will be be needed to describe them more precisely.

### The rhythmic contractions of the yolk persist, but are slower, in goldfish embryos acutely treated with a microtubule-depolymerizing drug prior to first cleavage

The formation of parallel array of microtubules at the vegetal yolk immediately after egg activation and fertilization, prior to the first cleavage, is known to be crucial to the proper DV patterning of fish embryos (Jesuthasan and Strähle, 1997; Tran et al., 2012). To investigate if goldfish yolk contractions are linked to this process, we treated goldfish embryos with nocodazole, a microtubule depolymerizing drug (Hoebeke et al., 1976). More precisely, at ∼ 10 mins post-fertilization, we incubated goldfish embryos in 0.1 *µ*g/mL nocodazole for ∼ 4 mins, as previously done in zebrafish (Jesuthasan and Strähle, 1997). Yolk contractions persisted even after acute drug treatment (Figure 5A, Supplementary Movie M8), but were however slower than in controls (Table 5, Figure 5A,E). Incidentally, we also noted persistent contractions in yolk that pinched off from a dying treated embryo (n = 4, Supplementary Figure F4, Supplementary Movie M9). To verify the effectiveness of our drug treatment, we screened embryonic phenotypes at 1 day post-fertilization (Supplementary Figure F3). As reported in zebrafish (Jesuthasan and Strähle, 1997), acute treatment of goldfish embryos with nocodazole prior to first cleavage results in axis patterning defects and mostly ventralized phenotypes (Supplementary Figure F3B). We have not verified the presence of a parallel array of microtubules in the vegetal yolk of goldfish, as described in zebrafish. Nonetheless, these results indicate that the contractions of the goldfish yolk do not rely on a similar microtubule assembly prior to the first cleavage. More interestingly to our investigation, these experiments reveal that rhythmic yolk contractions *per se* do not ensure fidelity of axis patterning and that the effect of the acute nocodazole treatment to DV patterning supersedes influence (if any) from the yolk contractions.

### The rhythmic contractions of the yolk persist, but are slower, in goldfish embryos treated with chemicals affecting actomyosin or calcium

We also performed pharmacological pertubations of actomyosin and calcium, as these were previously linked to periodic contractions in other embryological contexts.

To perturb actin, we treated goldfish embryos with 100 nM latrunculin (Özgüç et al., 2022), an actin polymerization inhibitor (Spector et al., 1983). We recorded persistent contractions in embryos treated with the drug from *∼* 10 mins post-fertilization (Figure 5B, Supplementary Movie M10). However these contractions were slower than in controls (Table 6, Figure 5B,E), and the drug treatment resulted in dead embryos (Figure F3B).

To perturb myosin, we treated goldfish embryos with 10 *µ*M blebbistatin(–) (Özgüç, 2021), a myosin inhibitor (Straight et al., 2003), from 3 hours post-fertilization and used its inactive enantiomer, 10 *µ*M blebbistatin(+), as control. We noted persistent contractions in both conditions (Figure 5C, Supplementary Movie M11), but the contractions in embryos treated with blebbistatin(–) were slower than in controls (Table 7, Figure 5C,E).

To perturb calcium, we incubated goldfish embryos in either double distilled water (ddH_2_O, Ca^2+^ free water) or 1 mM EDTA in ddH_2_O (Ca^2+^ chelating agent) from 3 hours post-fertilization. We observed persistent contractions in both ddH_2_O and EDTA conditions (Figure 5D, Supplementary Movie M12), but the EDTA treatment resulted in quick dying of goldfish embryos (Supplementary Movie M12, Supplementary Figure F3B). Notably, the period of contractions in embryos incubated in ddH_2_O is similar to tap water controls (Table 8, Figure 5D-E), as previously reported by Yamamoto (1954), implying that extracellular calcium might not be affecting the rhythmic yolk contractions. In contrast, the period of contractions in embryos incubated in EDTA is slower than controls (Table 8, Figure 5D-E).

For all pharmacological perturbations (including nocodazole treatment in the previous section but excluding incubation in ddH_2_O), the period of the yolk contractions were slower than in respective controls (Figure 5E). While the period quantification was taken at timepoints when the contractions were still robust (i.e. median wavelet power > 3, greater than the minimum wavelet power corresponding to 95% confidence interval in case of white noise (Mönke et al., 2020)), it is still unclear if the slower period in treated samples reflected the importance of the perturbed molecular machinery in the maintenance of the rhythmic yolk contractions or if it was a consequence of loss of embryo viability upon drug treatment. Overall, we find that the persistence of yolk contractions is highly robust to pharmacological perturbation of pathways driving contractions in most other studied contractile systems / eggs.

### The period of yolk contractions is faster in embryos of an established goldfish strain with atypical phenotype but is independent of *ChdS* genotype

We were curious if the yolk contractions permitted emergence of different morphotypes in goldfish, and thus if certain yolk dynamics are correlated with now established goldfish strains and if this relates to strain-specific genotypes. To address this, we first compared wild-type goldfish with the twin-tail goldfish *Oranda* strain, which is naturally more ventralized (Abe et al., 2014) and has a bifurcated caudal fin. We found that the yolk contractions in *Oranda* embryos are faster than in wild-type controls (Table 9, Figure 6, Supplementary Movie M13) with periods of 103.02 *±* 5.94 seconds (94.75–104.27 seconds, n = 25) and 111.88 *±* 6.72 seconds (109.45–115.27 seconds, n = 56), respectively. This shows that embryos from a population of goldfish with atypical, more ventralized, phenotype exhibit faster yolk contractions than their typical wild-type counterparts.

**Table 9.** Period of yolk contractions in wild-type and twin-tail *Oranda* goldfish embryos at 02:14:20–09:43:10 hh:mm:ss post-fertilization, expressed as median *±* median absolute deviation (95% confidence interval of the median). n and N corre-spond to the number of samples and to the number of clutches, respectively.

| condition | $n$ | $N$ | period (sec) |
| --- | --- | --- | --- |
| wild-type (WT) | 56 | 2 | 111.88±6.72 (109.45–115.27) |
| twin-tail *Oranda* (OR) | 25 | 2 | 103.02±5.94 (94.75–104.27) |

**Fig. 6.**
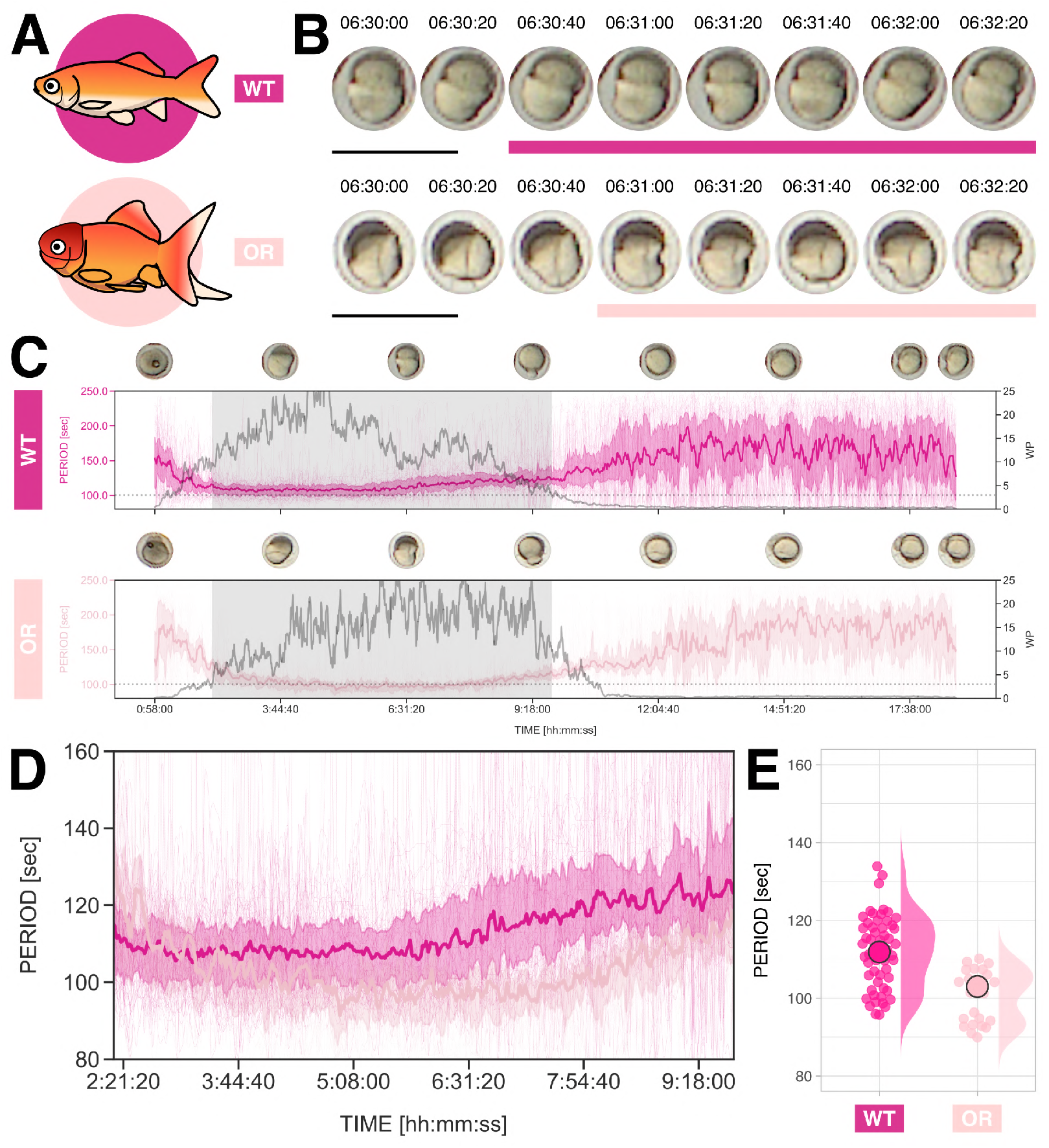
Embryos of *Oranda*, a twin-tail goldfish strain, exhibit faster rhythmic yolk contractions than wild-type. (**A**) Illustrations of wild-type (WT, dark pink) and twin-tail *Oranda* (OR, light pink) goldfish strains. (**B**) Snapshots of representative wild-type (top) and twin-tail *Oranda* (bottom) goldfish embryos for a subset of imaging time, at 06:30:00–06:32:20 hh:mm:ss post-fertilization. Solid color-coded bars mark snapshots spanning one full contraction. Scale bar = 2 mm. Timelapse of some of the wild-type and twin-tail *Oranda* goldfish embryos are shown in Supplementary MovieM13. (**C**) Temporal evolution of period of yolk contractions for the entire duration of the experiment, obtained from wavelet analysis of detrended timeseries (via sinc-filter detrending, cut-off period = 250 seconds). The period evolution for each sample and the median of the periods are represented as a color-coded dashed line and a color-coded solid line, respectively. The color-coded shaded area corresponds to the interquartile range. Snapshots of representative samples (same as those in panel B) are shown above each plot. Median wavelet power (WP) is also plotted in gray, with a horizontal dotted gray line marking wavelet power threshold = 3, which corresponds to 95% confidence interval in case of white noise. Shaded region corresponds to subset time duration considered for comparison of periods, i.e. 02:14:20–09:43:10 hh:mm:ss post-fertilization, where both conditions show median wavelet power > 3. (**D**) Temporal evolution of period of yolk contractions in wild-type (dark pink) and twin-tail *Oranda* (light pink) goldfish embryos for the specified subset of imaging time. The period evolution for each sample and the median of the periods are represented as a color-coded dashed line and a color-coded solid line, respectively. The color-coded shaded area corresponds to the interquartile range. (**E**) Median period of yolk contractions in wild-type (WT, n = 56) and twin-tail *Oranda* (OR, n = 25) goldfish embryos at 02:14:20–09:43:10 hh:mm:ss post-fertilization. Individual samples are represented as color-coded dots, while the median of all the samples is denoted as a color-coded circle with black margin. The distribution of the data is also plotted. These data are summarized in Table9.

Previously, it was found that the more ventralized twin-tail goldfish strains carry a *ChdS* allele that results in a non-functional truncated ChdS protein (Abe et al., 2014), i.e. *ChdS*(–) allele. This has been described as responsible for their atypical phenotype, and has been implicated even in the emergence of the dorsal finless goldfish strain (Abe et al., 2014; Chen et al., 2022; Lee et al., 2023). We then asked if the difference in period of yolk contractions between wildtype and twin-tail embryos is due to the difference in *ChdS* genotype. To tackle this, we analyzed yolk contractions in goldfish embryos of *ChdS*(+/–) x *ChdS*(+/–) matings, which results in embryos of all possible *ChdS* genotypes (Figure 7A-B). We performed simultaneous timelapse imaging of sibling embryos and subsequently genotyped them for *ChdS* (Figure 7C-D). We noted persistent yolk contractions across all genotypes (Figure 7E, Supplementary Movie M14). Strikingly, we did not record any marked difference between periods (Table 10, Figure 7F-H), indicating that the period of yolk contractions is independent of *ChdS* genotype. Remarkably, while we did not see significant difference in period between the three genotypes, we found that the period of contractions differs between clutches (Figure 7I). Since contractions could persist even in the absence of sperm (Figure 3, Supplementary Movie M4), the differences in the period between clutches could very likely be determined by the batch of eggs that was each recovered from different mothers. Incidentally, period variability across batches was also recovered in wild-type goldfish (not shown, but summarized in Table 1, Table 2, and Table 9).

**Table 10.** Period of yolk contractions in goldfish embryos with different *ChdS* genotypes at 02:45:50–11:44:10 hh:mm:ss post-fertilization, expressed as median *±* median absolute deviation (95% confidence interval of the median). n and N corre-spond to the number of samples and to the number of clutches, respectively.

| condition | $n$ | $N$ | period (sec) |
| --- | --- | --- | --- |
| *ChdS*(+/+) | 19 | 3 | 116.48±04.87 (112.83–123.87) |
| *ChdS*(+/–) | 28 | 3 | 117.93±05.90 (113.12–124.00) |
| *ChdS*(–/–) | 17 | 3 | 120.26±12.26 (113.95–137.11) |

**Fig. 7.**
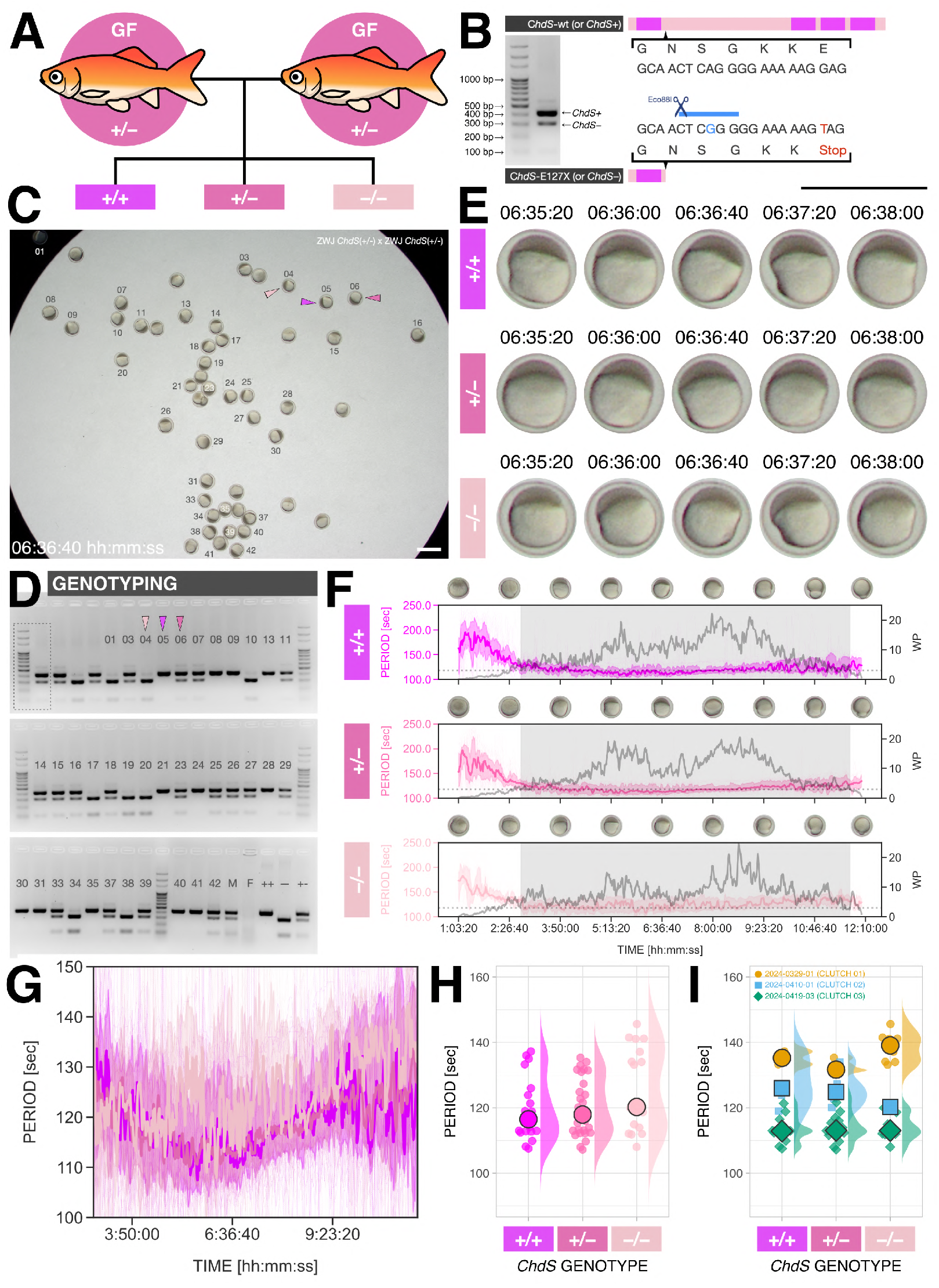
The period of yolk contractions in goldfish embryos is independent of *ChdS* genotype. (**A**) Schematic of experiment comparing the rhythmic yolk contractions in goldfish embryos with different *ChdS* genotypes recovered from mating of ZWJ-*ChdS*(+/–) goldfish. (**B**) Twin-tail goldfish strains have mutations in their *ChdS* gene, which result in a premature stop codon and the presence of an Eco88I restriction enzyme site, i.e. the *ChdS*(–) allele (Abe et al.,2014). For *ChdS* genotyping, *∼* 400 bp of the *ChdS* gene is amplified and is then treated with the Eco88I restriction enzyme, which recognizes the *ChdS*(–) allele-specific restriction enzyme site resulting in a shorter *∼* 300 bp (and *∼* 100 bp) band. (**C**) Snapshot of simultaneous imaging of goldfish embryos from a ZWJ-*ChdS*(+/–) x ZWJ-*ChdS*(+/–) mating at 06:36:40 hh:mm:ss post-fertilization. Numbers denote sample ID for genotyping. Color-coded arrowheads mark representative samples. Scale bar = 2 mm. Simultaneous timelapse of these goldfish embryos is shown in Supplementary MovieM14. (**D**) Result of *ChdS* genotyping of samples in panel C. Dotted rectangle marks lanes used to illustrate typical result of *ChdS* genotyping in panel B. Color-coded arrowheads mark genotyping results of representative samples in panels C. Numbers correspond to the sample IDs indicated in panel C. Unlabeled lanes refer to samples from another clutch. ++, —-, and +– lanes refer to *ChdS*(+/+), *ChdS*(–/–), and *ChdS*(+/–) controls, respectively. M and F lanes refer to male and female ZWJ-*ChdS*(+/–) goldfish, respectively, which were used for mating. Note that while no bands were seen for lane F, the combinations of *ChdS* genotypes in the embryos would only be possible if F was also *ChdS*(+/–). Snapshots of representative *ChdS*(+/+), *ChdS*(+/–), and *ChdS*(–/–) goldfish embryos for a subset of imaging time, at 06:35:20–06:38:00 hh:mm:ss post-fertilization. Scale bar = 2 mm. These representative samples are the same as those marked with color-coded arrowheads in panels C and D. (**F**) Temporal evolution of period of yolk contractions for the entire duration of the experiment, obtained from wavelet analysis of detrended timeseries (via sinc-filter detrending, cut-off period = 250 seconds). The period evolution for each sample and the median of the periods are represented as a color-coded dashed line and a color-coded solid line, respectively. The color-coded shaded area corresponds to the interquartile range. Snapshots of representative samples (same as those in panels C-E) are shown above each plot. Median wavelet power (WP) is also plotted in gray, with a horizontal dotted gray line marking wavelet power threshold = 3, which corresponds to 95% confidence interval in case of white noise. Shaded region corresponds to subset time duration considered for comparison of periods, i.e. 02:45:50–11:44:10 hh:mm:ss post-fertilization, where all conditions show median wavelet power > 3. (**G**) Temporal evolution of period of yolk contractions in *ChdS*(+/+) (magenta), *ChdS*(+/–) (dark pink), and *ChdS*(–/–) (light pink) goldfish embryos for the specified subset of imaging time. The period evolution for each sample and the median of the periods are represented as a color-coded dashed line and a color-coded solid line, respectively. The color-coded shaded area corresponds to the interquartile range. (**H**) Median period of yolk contractions in *ChdS*(+/+) (n = 19), *ChdS*(+/–) (n = 28), and *ChdS*(–/–) (n = 17) goldfish embryos at 02:45:50–11:44:10 hh:mm:ss post-fertilization. Individual samples are represented as color-coded dots, while the median of all the samples is denoted as a color-coded circle with black margin. The distribution of the data is also plotted. These data are summarized in Table10. (**I**) Median period of yolk contractions in different clutches of goldfish embryos at 02:45:50–11:44:10 hh:mm:ss post-fertilization. Data are categorized based on *ChdS* genotype, as in panel H, but the colors denote different embryo clutches instead. Individual samples are represented as color-coded dots, while the median of all the samples from the same clutch within a genotype is denoted as a color-coded circle with black margin. The distribution of the data is also plotted.

**Fig. 8.**
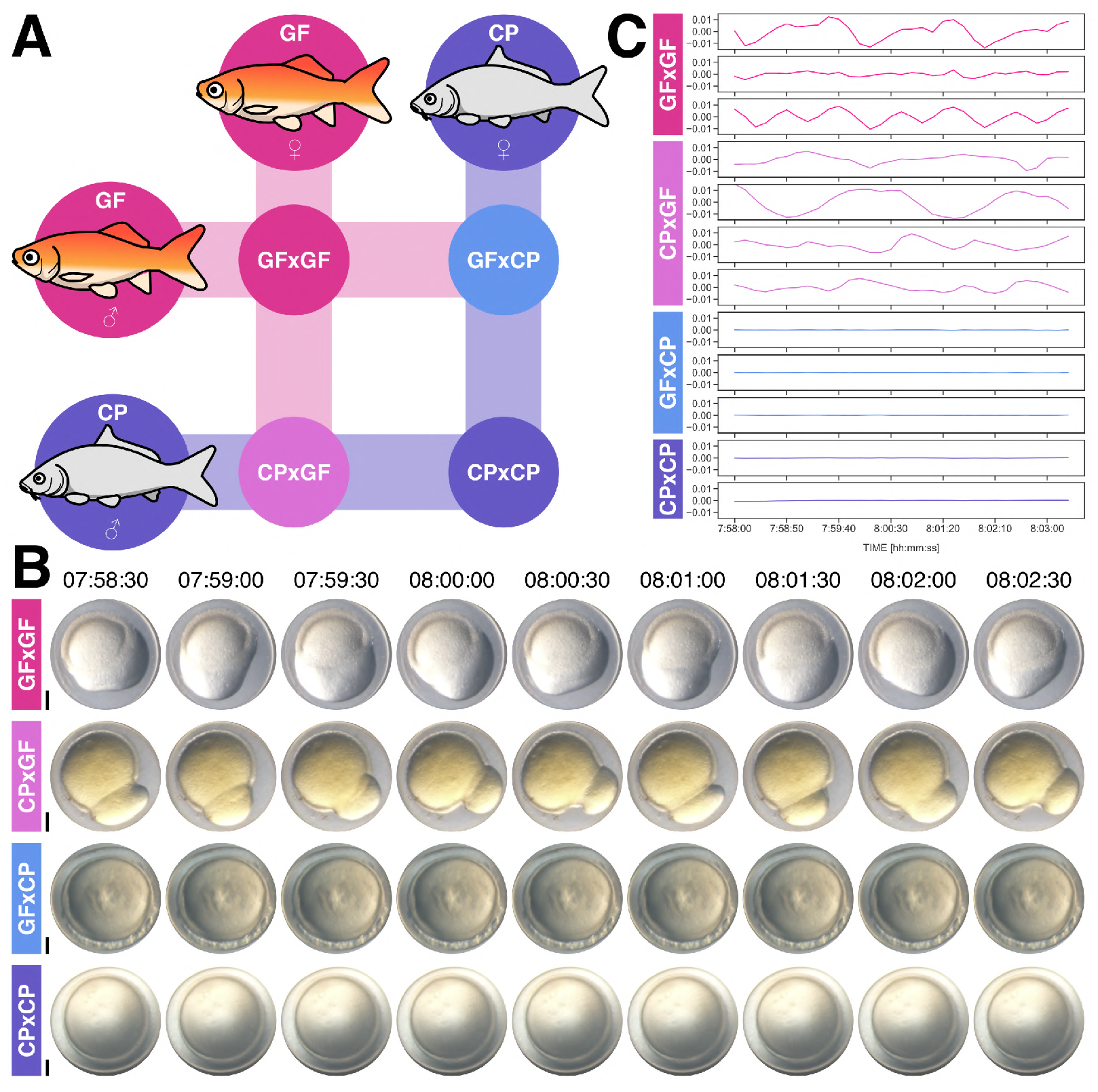
The rhythmic contraction of the yolk is a trait that is maternal in origin. (**A**) Schematic of experiment comparing embryos of goldfish (GFxGF, dark pink), of common carp (CPxCP, dark blue), and of their reciprocal hybrids: one that is from common carp sperm and goldfish egg (CPxGF, light pink), and another that is from goldfish sperm and common carp egg (GFxCP, light blue). (**B**) Snapshots of representative GFxGF, CPxGF, GFxCP, and CPxCP fish embryos for a subset of imaging time, at 07:58:30–08:02:30 hh:mm:ss post-fertilization. Scale bar = 250 *µ*m. Timelapse of these fish embryos are shown in Supplementary MovieM15. (**C**) Detrended timeseries (via sinc-filter detrending, cut-off period = 850 seconds) of GFxGF (n = 3), CPxGF (n = 4), GFxCP (n = 3) and CPxCP (n = 2) at 07:58:00–08:03:20 hh:mm:ss post-fertilization.

### The rhythmic contraction of the yolk is a trait that is maternal in origin

We tested the maternal origin of the contractions using reciprocal hybrids of goldfish and common carp (Figure 8). Specifically, we generated reciprocal goldfish-carp hybrids by artificial fertilization (Tsai et al., 2013) (Figure 8A): (a) in one type of hybrids the egg was derived from goldfish and the sperm from common carp, while (b) in the other type of hybrids the egg was derived from common carp and the sperm from goldfish. We observed that yolk contractions were only detectable in embryos where the egg had been derived from goldfish (Figure 8B-C, Supplementary Movie M15). We conclude that the contractions of the yolk are of maternal origin. In other words, and as supported by the results of the previous section, the presence and the period of rhythmic yolk contractions in fish embryos depend on the egg.

## Discussion and open questions

The egg exhibits a range of dynamic behaviors during development and patterning such as cytoskeletal reorganisation, calcium signaling oscillations, waves, and contractile behavior (Brownlee and Dale, 1990; Deguchi et al., 2000; Ishii and Tani, 2021; Just, 1919, 1939; Kyozuka et al., 2008; Limatola et al., 2022a; Santella and Chun, 2022; Sardet et al., 1998; Stricker, 1999) most evident immediately after egg activation and fertilization. Remarkably, in goldfish eggs and embryos (but not in those of closely-related common carp and zebrafish) rhythmic contractions of the yolk emerge after some time elapsed from egg activation and fertilization and persist until gastrulation. These contractions are maternal and do not require sperm entry / fertilization nor cell division.

That contractions emerge in unfertilized goldfish eggs at the same time as in fertilized ones was actually already observed by Yamamoto (1954). Interestingly, in trout eggs that also exhibit rhythmic yolk contractions (Wülker, 1953; Yamamoto, 1940), periodic impedance changes also emerge at the same time between fertilized and unfertilized samples (Hubbard and Rothschild, 1939). Egg activation does not require sperm and will be initiated when goldfish eggs contact water (Lee et al., 1999; Yamamoto, 1954), thus possibly functioning as a trigger and “time 0” reference for the emergence of yolk contractions even in unfertilized eggs. We currently do not know why yolk contractions do not begin immediately after egg activation but instead become apparent only at a precise time (i.e. the 4-cell stage of fertilized eggs). This could perhaps be related to the timing of the 4-cell stage as a milestone in DV axis specification. For instance, maternal *sqt* RNA (*squint/nodal-related 1*), which is involved in DV patterning (Lim et al., 2012), is distributed evenly in the egg but is transported to the animal pole via microtubules upon egg activation, even in the absence of sperm (Gore and Sampath, 2002). In fertilized eggs, it eventually becomes asymmetrically localized in 4-cell zebrafish embryos (Gore et al., 2005). Additionally, removal of the vegetal yolk after the 4-cell stage via surgical bisection along the equator of fish embryos does not result in severe ventralized phenotypes in goldfish (as is instead the case when the vegetal yolk was removed by Tung et al. (1945) and Mizuno et al. (1997) at the 2-cell stage), suggesting that a sufficient amount of the maternal dorsal determinants have been transported to the prospective dorsal organizer only by the 4-cell stage (Tung et al., 1955b). Whether or not this timing is linked to the appearance of rhythmic yolk contractions is still unclear. Do the yolk contractions need the entire time from egg activation to organize their machinery (provided there is any)? Or do they already have the material/physical/chemical conditions for their execution early on, but “wait” until time equivalent to the 4-cell stage (e.g. until triggering of an activator and/or decay of an inhibitor)?

We hypothesize that the yolk contractions in goldfish are maintained by the cortical actomyosin network and/or calcium signaling. Both of these factors correlate spatiotemporally with contractile behavior in the ascidian egg after egg activation and fertilization (Brownlee and Dale, 1990; Ishii and Tani, 2021). Waves of calcium signaling are also involved in the deformation of mouse oocytes (Deguchi et al., 2000) and in the rhythmic contraction of the blastoderm in medaka embryos (Simon and Cooper, 1995). Softening of the cortical actomyosin network has also been recently implicated in the periodic cortical waves of contraction of mouse embryos prior to compaction (Özgüç et al., 2022). Periodic impedance changes were previously recorded in trout, which are believed to be linked to the periodic yolk contractions in this fish and are inferred to be more precisely due to the periodic thickening-thinning of the cortex (Hubbard and Rothschild, 1939; Rothschild, 1940, 1947; Wülker, 1953; Yamamoto, 1940). That the rhythmic contractions persist in isolated yolk, e.g. in vegetal yolk separated from the rest of the embryo (Figure 4, Supplementary Movie M6, Supplementary Movie M7, and as described by Mizuno et al. (1997) and by Wülker (1953) (but in trout and pike)) and in yolk pinched off from a dying embryo (Supplementary Figure F4, Supplementary Movie M9), implies that the yolk itself is sufficient to exhibit said dynamic behavior (and that the yolk contractions do not rely on the embryo proper). Current knowledge on the link between cortical actomyosin, calcium signaling, and contractile behavior mainly focus on these phenomena in cells. It is thus curious to further investigate if these three similarly act in concert in an acellular biological system like the yolk. Remarkably, surface deformations were seen in water-in-oil droplets encapsulating *Xenopus laevis* egg extracts, which among other things contain physiological levels of actomyosin (Sakamoto et al., 2022, 2020). Indeed, it has been shown that an actomyosin network is sufficient to cause contractions in minimal synthetic models of cells. By tuning the ratio between actin and myosin, contractions can also be induced in cortical actomyosin network reconstituted on a coverslip, in emulsion, or in lipid vesicles (Litschel et al., 2021; Miyazaki et al., 2015; Tsai et al., 2011). Contraction of the cortical actomyosin network in lipid vesicles could even result in surface deformations (Litschel et al., 2021). These deformations are however not periodic, suggesting that additional factors are likely involved in periodic contractions like those of the goldfish yolk. Indeed, linking actomyosin to calcium signaling in a theoretical model captures periodic deformations of non-adhering fibroblasts (Salbreux et al., 2007).

In terms of the function of the goldfish yolk contractions, could yolk contraction dynamics (e.g. period) confer patterning cues to the developing fish embryo? Considering that the yolk carries material crucial for embryonic patterning, could yolk contractions serve as a “stirrer” as already suggested by Yamamoto (1934)? If the contractions serve as a stirrer, then they could be spreading maternally-derived dorsal determinants resulting in their dilution at the prospective dorsal organizer. This could be lowering the density of dorsalizing molecules available to counteract specification of ventral fates, e.g. via maternal *radar/gdf6a* (Goutel et al., 2000; Rissi et al., 1995; Sidi et al., 2003), therefore priming embryos for ventralization. In this “stirrer” framework (Figure 9), faster and/or stronger yolk contractions lead to further dilution of dorsal determinants and are thus correlated with more ventralized phenotypes. However, we found here that goldfish embryos acutely treated with nocodazole prior to the first cleavage actually exhibit slower yolk contractions despite being more ventralized than controls. This suggests that the effect of the drug treatment, which likely disrupted the microtubule assembly important for asymmetric distribution of dorsal determinants as in zebrafish (Jesuthasan and Strähle, 1997), supersedes any patterning influence from the yolk contractions. Interestingly, faster calcium transients have been recorded in *hecate/grip2a* zebrafish mutants, which are ventralized (Ge et al., 2014; Gingerich et al., 2005). To the opposite, slowing down calcium transients via pharmacological perturbation of Ca^2+^ release results in ectopic expression of the dorsal marker *chordin* in zebrafish (Gingerich et al., 2005; Westfall et al., 2003). In fact, the zebrafish dorsal organizer has been shown to be a calcium pacemaker at the gastrula stage (Chen et al., 2017; Gilland et al., 1999). If calcium signaling is involved in goldfish yolk contractions, could these contractions be acting as an early determinant or an early readout of DV patterning in these fish? In the presence of contractions, could there be actual spatial spreading of maternally-derived dorsal determinants at the prospective dorsal organizer? And/or are there altered temporal dynamics between said molecules and target cells? How would this be orchestrated with cytoskeleton-dependent transport?

**Fig. 9.**
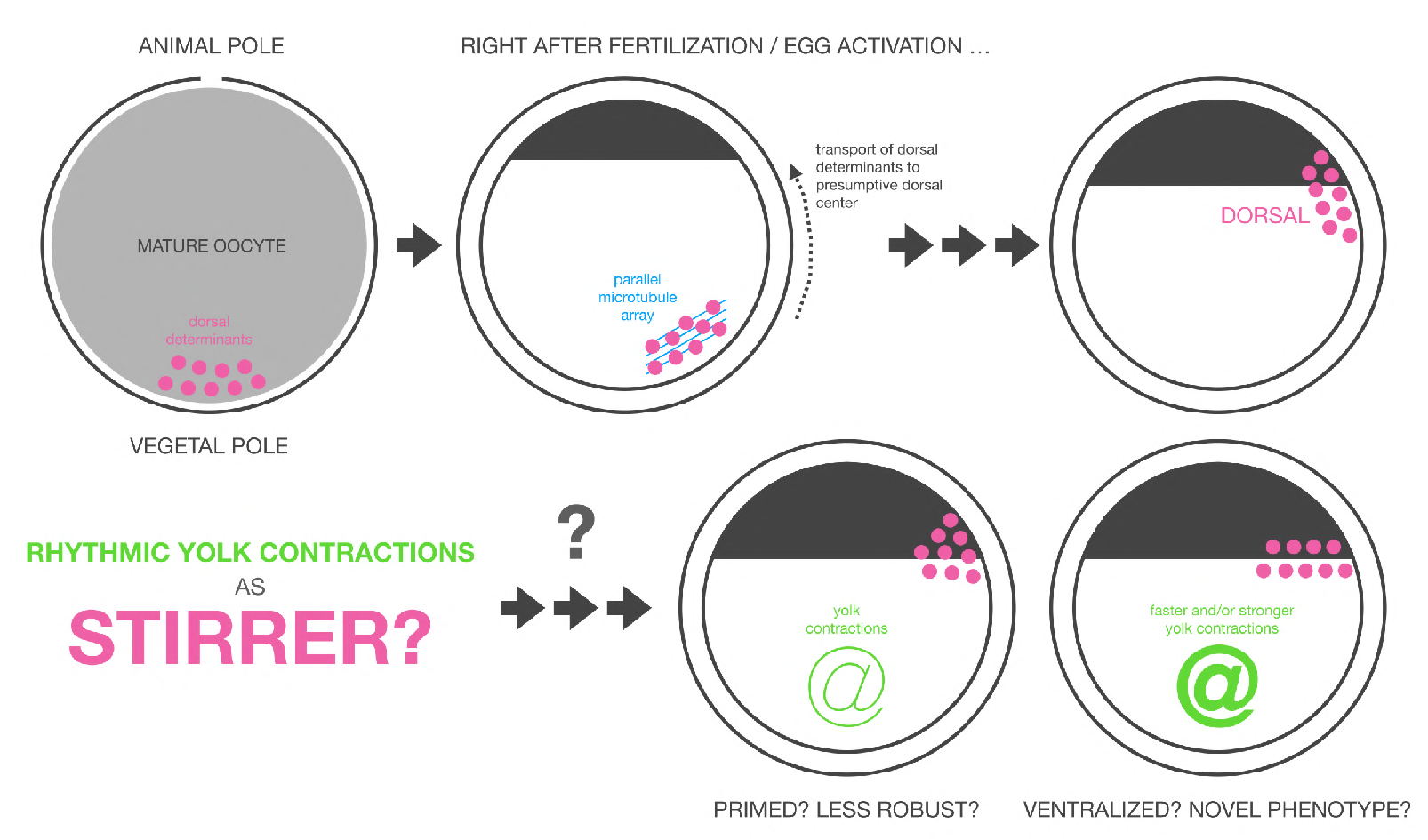
Stirrer hypothesis: could the rhythmic yolk contractions serve as a “stirrer” during early fish embryo development and patterning? After egg activation / fertilization, dorsal determinants deposited at the vegetal pole of mature fish oocytes are directed at an angle via a parallel microtubule array that forms at the vegetal side of the egg prior to the first cleavage (Jesuthasan and Strähle, 1997; Mizuno et al., 1999, 1997; Ober and Schulte-Merker, 1999; Tran et al., 2012). These determinants are then transported towards the animal pole, at the prospective dorsal center (Ge et al., 2014; Jesuthasan and Strähle, 1997; Tran et al., 2012). In the presence of rhythmic yolk contractions, as in goldfish embryos, the transport of these dorsal determinants from the vegetal pole to their target site are likely disturbed. If yolk contractions serve as a “stirrer”, as previously suggested by Yamamoto (1934), then could they be spreading the determinants to a broader area, which would result in dilution at the prospective dorsal region and could permit the emergence of more ventralized phenotypes? More generally, could yolk contractions perturb early fish embryo development and patterning, thus lessening its robustness? Could this then permit emergence of novel phenotypes under strong selective pressures (e.g. the initial emergence of twin-tail and dorsal finless phenotypes during the domestication of goldfish), which could later then be fixed by acquisition of genetic mutations (e.g. the *ChdS*(-) allele in twin-tail goldfish strains)? Scheme on top row is adapted and modified from Ge et al. (2014); Hibi et al. (2002, 2018); Langdon and Mullins (2011); Nojima et al. (2010).

Strikingly, altered DV patterning is implicated in the variation of median fin morphotypes in goldfish, such as twin-tail (Abe et al., 2014) and dorsal-finless (Chen et al., 2022). The yolk, and not the genetic material *per se*, was previously demonstrated to be important in the emergence different morphotypes in goldfish strains. In particular, Cai (1989) was able to recover single-tail goldfish from twin-tail goldfish embryos only by yolk suction. This could be interpreted as due to the removal of dorsal determinant-poor yolk, resulting in increase of concentration of the dorsal determinants and rescue of the ventralized twin-tail phenotype. Here, we documented that embryos of *Oranda*, a twin-tail goldfish strain, exhibit faster yolk contractions than wild-type goldfish embryos, correlating faster contractions to atypical, more ventralized, phenotypes in alignment to the current “stirrer” framework. Periodic contractions of the yolk were documented in embryos of other fishes, e.g. those of the Japanese icefish *Salangichthys microdon* (Yamamoto, 1938), those of the pike *Esox lucius* (Kasansky, 1936; Ransom, 1867; Rusconi, 1840; Wülker, 1953), those of the trout *Onchorhynchus masou* and *Salmo* (Lereboullet, 1862; Wülker, 1953; Yamamoto, 1940), and those of the stickleback *Gasterosteus aculeatus* (Ransom, 1867; Wintrebert and Yung, 1926). However, when considering a closely-related species with the same ecological niche, we noted that common carp does not show rhythmic yolk contractions as goldfish. Could wildtype goldfish therefore already be more ventralized than common carp? Interestingly, while it is easy to induce bifurcated tail fin in wild-type goldfish (e.g. by injecting morpholinos targeting *ChdS* in embryos), similar ventralized phenotypes could not be achieved with common carp (Abe et al., 2014, 2016). Notably, when investigating reciprocal goldfish-carp hybrids, we noted that rhythmic yolk contractions are present in hybrids from goldfish eggs. The egg cytoplasm was previously determined to impact traits (e.g. vertebra number) in goldfish-carp hybrids (Sun et al., 2005). Could hybrids exhibiting rhythmic yolk contractions then be more sensitive to perturbations resulting in bifurcated tail fin? Are these embryos primed for ventralization?

Embryos of the even closer related *Carassius auratus indigentiaus* subsp. nov. have been shown to also exhibit rhythmic yolk contractions (Zhang et al., 2023). It thus seem likely that the ancestral *Carassius auratus* goldfish initially used in domestication also exhibited rhythmic yolk contractions. Could yolk contractions then be permissive of the emergence of median fin morphotypes in goldfish? We observed a distribution of periods of yolk contractions in each clutch of goldfish eggs, with some contracting slightly faster than the rest. It could be that development and patterning is less robust in these fast-contracting embryos, and the lowering of robustness could then allow emergence of atypical morphotypes especially when these embryos are subjected to selective pressures like domestication. We found here that the presence and the period of yolk contractions is a trait that is maternal in origin. If the period of contractions is something that can be passed on from one generation to the next, then the fast contractions in present-day twin-tail *Oranda* embryos reflect the faster yolk contractions among goldfish that first showed the twin-tail morphotype versus their wild-type counterparts / siblings. That novel morphotypes could have a higher likelihood of emergence in goldfish that have fast yolk contractions implies that faster-contracting twin-tail goldfish strains could further acquire other novel morphotypes. Indeed, it has been documented that goldfish strains showing the even more ventralized dorsal finless morphotype (e.g. the *Ranchu* strain) were actually derived from twin-tail goldfish (Chen et al., 2022; Matsui, 1935; Ota, 2021). Considering this, more generally, could yolk contractions be perturbing the robustness of early fish development and patterning (e.g. disrupting early DV patterning), permitting the emergence of novel phenotypes (e.g. twin-tail morphotype) under strong selective pressures (e.g. goldfish domestication) which are then only later fixed by acquisition of genetic mutations (e.g. *ChdS*(–) allele)?

## Future perspectives

We here propose further experiments to address open questions mentioned in the current study, and aimed to further elucidate the origin, the maintenance, and the role of rhythmic yolk contractions in goldfish.

### On investigating yolk contraction and its dynamics

It would be curious to investigate the role of egg activation on the onset of yolk contractions. Some goldfish eggs could be acutely treated with protease inhibitors (e.g. aprotinin) or ovarian fluid to delay their egg activation (Ma and Carney, 2024; Minin and Ozerova, 2008). The onset of rhythmic yolk contractions in these treated eggs could then be compared with those in control conditions. Another prospective experiment could involve tying or bisecting goldfish embryos along the first cleavage plane prior to the onset of yolk contractions, as previously done by Mizuno et al. (1997); Tung et al. (1955a); Tung and Tung (1943) and as recently suggested by Saunders (2024). The separated samples could then be imaged and the dynamics of yolk contractions (if present) between the pairs be compared. Additionally, it would be insightful to investigate the period of yolk contractions across generations. For instance, it is interesting to compare the period of yolk contractions in eggs of siblings that show markedly different periods of yolk contractions when they were embryos. Simultaneous timelapse and quantification of yolk contractions in wild-type, twin-tail *Oranda*, and the even more ventralized dorsal finless *Ranchu* goldfish strains would also be very informative.

Alternative ways to extract the timeseries could be considered, especially those able to describe the yolk contractions in 3D. Through these more complete approaches, and perhaps in tandem with particle image velocimetry (Özgüç et al., 2022; Pereyra et al., 2021, 2024), one could more clearly relate the propagation and number of the waves of contractions to the position of the prospective dorsal organizer. This also enables more thorough quantification of the effect of yolk size on the yolk deformations and the period of contractions, e.g. in yolk isolated from the rest of the embryo. This necessitates imaging the embryo at higher spatial (i.e. 3D) and temporal resolutions (contractions are at the 10^2^ seconds range), requiring more sophisticated imaging modalities, likely at the expense of throughput. Quantitative measurements of the deformability of the yolk and of other viscoelastic properties would also be informative and could be achieved through micropipette aspiration (Guevorkian and Maître, 2017). In collaboration with theoreticians, modeling and numerical experiments could be done, e.g. on linking dynamic phenomena to transfer of material from one part of a contracting/deformable sphere to another and on the propagation of deformation along the yolk surface.

### On investigating cytoskeleton and its dynamics

To elucidate the contribution of the cytoskeleton on the rhythmic yolk contractions, it would be insightful to image and to quantify yolk contraction dynamics in goldfish embryos treated with low concentrations of either latrunculin, which interferes with actin polymer assembly (Spector et al., 1983), or blebbistatin, which disrupts interactions of actin and myosin (Straight et al., 2003). While this drug treatments were already done in this study, it is still unclear if the effects we see are due to the perturbation of actomyosin or due to the loss of sample viability. Titration of the drug concentrations would be important to determine the concentration of drug that would have a detectable effect on the period of contractions (if any) without drastically affecting viability of the embryos. Generation of transgenic goldfish expressing an actin reporter, e.g. fluorescent protein fused to actin-binding domain of utrophin (Burkel et al., 2007), would be advantageous to study actin dynamics during yolk contractions. While CRISPR-Cas has been done in goldfish (Lee et al., 2023; Yu et al., 2022), generation of transgenic lines remains challenging due to the restricted spawning season of this fish. Alternatively, goldfish embryos could be injected with fluorescent reporters of actin (co-injected with inert fluorescent dye as counterstain) prior to timelapse imaging, e.g. fluorescently labeled phalloidin as done by Kyozuka et al. (2008) in starfish oocytes, or mRNA of LifeAct fused to a fluorescent protein as done by Foster et al. (2022) also in starfish oocytes, or LifeAct-GFP protein as done by (Limatola et al., 2022b) in sea urchin eggs. Immunostaining (e.g. with β-tubulin antibody and labeled phalloidin) of fixed goldfish embryos could also be done to evaluate the organization of the cytoskeleton at different phases of the yolk contractions.

### On investigating calcium signaling and its dynamics

To elucidate the contribution of calcium signaling on the rhythmic yolk contractions, it would be insightful to image and to quantify yolk contraction dynamics in goldfish embryos treated with either phorbol 12-myristate 13-acetate (PMA) or bisindolylmaleimide I (BIM), which have been shown to speed up and slow down Ca^2+^ oscillations, respectively (Halet et al., 2004). Generation of transgenic goldfish expressing reporter of calcium signaling, e.g. GCaMP6s (Chen et al., 2013), would be advantageous to study calcium signaling dynamics during yolk contractions. As an alternative to generating a transgenic reporter strain e.g. via CRISPR-Cas, goldfish embryos could be injected with fluorescent calcium indicators (co-injected with inert fluorescent dye as counterstain) prior to timelapse imaging, e.g. Calcium Green Dextran (CGD) as done by Mohri and Kyozuka (2022) in starfish oocytes, or fluo-3 as done by Tsuruwaka et al. (2007) in zebrafish embryos.

### On investigating early DV patterning in goldfish embryos in relation to their yolk contractions

To relate the contraction of the yolk to DV patterning, and resolve their role as determinants or readouts, markers of DV patterning (e.g. *goosecoid, chordin*, nuclear β-catenin) could be used as additional readouts in the experiments cited above. The distribution of maternal dorsal determinants (e.g. *wnt8a* RNA) could be monitored either via in-situ hybridization of fixed embryos at different cleavage stages or via imaging of fluorescently-labeled *wnt8a* injected into live embryos, as done by Tran et al. (2012) in zebrafish embryos. Systematic comparison of the dorsal organizer domain in contractile goldfish and non-contractile common carp would also be informative. Injection of *ChdS* morpholino into goldfish-carp hybrid embryos would resolve whether a bifurcated tail fin can be induced in carp (hybrids) provided rhythmic yolk contractions occur.

## Methods

### Fish strains

Wild-type (ZWJ-*ChdS*(+/+) strain, locally known as 朱文錦) and twin-tail goldfish (*Carassius auratus*) were obtained from an aquarium supplier in Taiwan. Common carp (*Cyprinus carpio*) were obtained from local breeders or directly caught from the Erlong (二龍) river system in Yilan, Taiwan. Zebrafish (*Danio rerio*) were obtained from the Taiwan Zebrafish Core Facility at Academia Sinica (TZCAS). All experiments were conducted according to guidelines and upon approval of the Institutional Animal Care and Use Committee (IACUC) of Academia Sinica (Protocol #19-11-1351 and Protocol #22-11-1922).

### Artificial fertilization of goldfish and common carp eggs

During spawning season (March to June), artificial fertilization was performed as described in Tsai et al. (2013). Briefly, on the night before artificial fertilization, mature goldfish were injected with Ovaprim (Syndel, US) to stimulate production and maturation of gametes: ∼ 0.5 mL Ovaprim per kg of female fish and ∼ 0.1 mL Ovaprim per kg of male fish. After 12-16 hours, sperm and egg were collected. To collect sperm, male fish were gently squeezed near the cloaca and sperm was collected into labeled syringe prefilled with Modified Kurokawa’s extender 2 solution. Sperm activity was assessed by diluting a drop of sperm with water on a coverslip and visualizing sperm motility under the microscope. After confirming sperm activity, sperm was kept at 4*^o^*C until use. To collect mature eggs, female fish was squeezed near the cloaca, which had been pat-dried, and eggs were collected onto PTFE dish. Adult fish were anesthetized in MS-222 (Supelco, #A5040) before handling, and were immediately allowed to recover in fresh tap water afterwards.

Drops of sperm were mixed with the eggs and the mixture was transferred to labeled plastic wells / dishes (pre-coated with “Cha-Li-Wang” green tea (茶裏王, Uni-President Corp., Taiwan) to ease detachment of embryos from dish) containing tap water. For experiments involving unfertilized eggs, eggs were immediately transferred to dish containing tap water, without addition of sperm. After around 5 mins, water was removed and the eggs were washed at least 5 times with tap water. Optionally, the eggs were bleached with 0.1% (v/v, 1 mL in 1 L) bleach (Magic Amah, Taiwan) for 5 mins and quickly neutralized with 0.05% (w/v, 0.5 g in 1 L) sodium thiosulfate prior to washing. Eggs and embryos were kept in tap water at room temperature (*∼*24*^o^*C).

### Mating and embryo recovery of zebrafish

On the late afternoon before mating, mature male (1) and female (1-2) zebrafish were put in a mating tank and were separated by a physical barrier. At dawn, when the lights were turned on, the barrier was removed and fish were allowed to mate. Embryos were recovered ∼ 5-10 mins after egg release, washed with E3 medium, transferred to labeled plastic dish containing E3, and incubated at room temperature (*∼*24*^o^*C).

### Crude DNA extraction and genotyping

After timelapse imaging, at 1 day post-fertilization, goldfish embryos were given sample IDs, carefully transferred to separate wells, and were kept at 24*^o^*C. From 3 days post-fertilization, crude DNA extraction was performed as described by Kosuta et al. (2018). Briefly, each sample was sacrificed by incubation with ice for at least 20 minutes and was then transferred to a labeled 1.5 mL tube. Water was removed and 50 *µ*L of 50 mM NaOH (J.T.Baker, #3722-01) was added. The sample was subsequently incubated at 95*^o^*C for 1 hour with gentle rocking. Afterwards, it was vortexed and then was placed in ice for at least 10 mins. After cooling down, 6 *µ*L of 1 M Tris-HCl (pH = 8) was added. The sample was vortexed and then centrifuged at 1500 rcf for 1 min at room temperature. The supernatant was transferred to a new tube and was used for genotyping. Genotyping for the goldfish *ChdS* allele was done as described by Abe et al. (2014). In a 50 *µ*L reaction mixture, 3 *µ*L of crude DNA was mixed with 25 *µ*L of KOD One PCR Master Mix - Blue - (Toyobo, #KMM-201), 0.15 *µ*L of 10 *µ*M forward primer, 0.15 *µ*L of 10 *µ*M reverse primer, and 21.7 *µ*L of distilled water (Invitrogen, #10977-015). The sequences of the primers are: 5’-TAACGCACAGATGCAGACGTGTG-3’ (forward) and 5’-TGCTGTTCTCCTCAGAGCTGATGTAGG-3’ (reverse).

PCR was performed with (a) an initial denaturation at 98*^o^*C for 3 mins, (b) 35 cycles of 98*^o^*C for secs, 62*^o^*C for 5 secs, and 68*^o^*C for 1 sec, and (c) a final elongation at 68*^o^*C for 10 mins. Samples were kept at 4*^o^*C until use. To 20 *µ*L of PCR product, 1 *µ*L of Eco88I restriction enzyme (Thermo Scientific, #FD0384) was added. The mixture was incubated for 1 hour at 37*^o^*C to facilitate enzyme digestion and then for an additional 20 mins at 65*^o^*C to inactivate the enzyme. Eco88I recognizes a restriction enzyme site in the *ChdS*(–) allele, digesting the ∼ 400 bp PCR product to shorter ∼ 300 bp and ∼100 bp fragments (Abe et al., 2014), as illustrated in Figure 7B. To visualize the fragments, 2% agarose was first prepared by dissolving agarose (VWR Life Science, #0710) in TAE buffer and then adding 5 *µ*L DNA stain (Protech, #PT-D1001) per 100 mL mixture. 10 *µ*L of the enzyme-treated sample or 100 bp MW ladder (Purigo, #PU-GDM-101) was loaded into a well of the agarose gel that was submerged in TAE buffer. Agarose gel electrophoresis was done at 135 V for 30 mins. DNA bands were visualized using a UV transilluminator (Gel Doc XR+, Bio-Rad) and photos were taken using the Quantity One software (Bio-Rad).

### Dechorionation and equatorial bisection of goldfish embryos

To minimize adhesiveness of goldfish embryos and improve ease of handling, embryos were coated with milk prior to dechorionation. Isolated goldfish egg and sperm were mixed on PTFE dish and then transferred to a 1 L plastic conical beaker (Kartell, Italy) with 200-300 mL of ∼ 0.2 g/mL skim milk (安佳脫脂奶粉, Fonterra, Taiwan) in tap water. The conical beaker was constantly swirled for 5 mins and then 400-500 mL of tap water was added. Embryos were incubated in the milk mixture for at least 1 hour at room temperature (∼24*^o^*C) with constant stirring. After incubation, these were washed with tap water and subsequently transferred to dishes coated with 1% agarose (in tap water). These samples were then treated with 10 mg/mL pronase (Sigma-Aldrich, #P5147) for at most 5 mins. After pronase treatment, embryos were immediately washed for at least two times with E3 medium and kept in fresh E3 until fully dechorionated. At times, the chorion had to be gently peeled off from the embryo using forceps. Some of the milk-coated embryos were not dechorionated and were kept as controls (i.e. the with-chorion condition). After dechorionation, dechorionated embryos were transferred to an agarose-coated dish filled with tap water. Some of the dechorionated samples were bisected equatorially (i.e. separating the vegetal and the animal poles) using an eyelash tool. Because of challenges in dechorionation of early goldfish embryos, surgical manipulations were performed only from the 32-cell stage.

### Pharmacological perturbation

Stock solutions of the drugs were prepared: 2 mg of nocodazole (Sigma-Aldrich, #M1404) was mixed with 1 mL of DMSO (J.T.Baker, #9224-01 0121 24) to prepare 2 mg/mL nocodazole stock, 100 *µ*g of latrunculin A (Tocris, #3973) was mixed with 237.22 *µ*L of DMSO to prepare 1000 *µ*M latrunculin A stock, 1 mg of blebbistatin(–) (Tocris, #1852) was mixed with 34.207 *µ*L of DMSO to prepare 100 mM blebbistatin(–) stock, and 1 mg of blebbistatin(+) (Tocris, #1853) was mixed with 34.207 *µ*L of DMSO to prepare 100 mM blebbistatin(+) stock. Working solutions were freshly prepared: (a) 2 *µ*L of 2 mg/mL nocodazole in 40 mL solution with tap water (final concentration = 0.1 *µ*g/mL nocodazole) and 2 *µ*L of DMSO in 40 mL solution with tap water (as control), (b) 4 *µ*L of 1000 *µ*M latrunculin A in 40 mL solution with tap water (final concentration = 100 nM) and 4 *µ*L of DMSO in 40 mL solution with tap water (as control), (c) 4 *µ*L of 100 mM blebbistatin(–) in 40 mL solution with double distilled water (final concentration = 10 *µ*M) and 4 *µ*L of 100 mM blebbistatin(+) in 40 mL solution with double distilled water (final concentration = 10 *µ*M, as control), and (d) 80 *µ*L of 0.5 M EDTA (pH = 8) in 40 mL solution with double distilled water (final concentration = 1 mM).

For experiment with nocodazole, as in Jesuthasan and Strähle (1997), at ∼ 10 mins post-fertilization, goldfish embryos were incubated in 0.1 *µ*g/mL nocodazole (or 1:20000 DMSO) for ∼ 4 mins. After incubation, embryos were washed at least 5 times with tap water. Embryos were then kept in tap water.

For experiment with latrunculin, 100 nM latrunculin A was used as in Özgüç et al. (2022). Goldfish embryos were incubated in 100 nM latrunculin A (or 1:10000 DMSO control) from *∼* 10 mins post-fertilization.

For experiment with blebbistatin, 10 uM blebbistatin was used as in Özgüç (2021). Goldfish embryos were incubated in tap water and then kept in 10 *µ*M blebbistatin(–) (or 10 *µ*M blebbistatin(+) control) from 3 hours postfertilization. Due to unavailability of spawning wild-type (ZWJ-*ChdS*(+/+) strain) goldfish, ZWJ-*ChdS*(–/–)-singletail goldfish was used.

For experiment with double distilled water (ddH_2_O) and EDTA, goldfish embryos were incubated in tap water and then kept in either ddH_2_O or 1 mM EDTA in ddH_2_O (or tap water control) from ∼ 3 hours post-fertilization. Due to unavailability of spawning wild-type (ZWJ-*ChdS*(+/+) strain) goldfish, ZWJ-*ChdS*(–/–)-singletail goldfish was used.

For all pharmacological perturbations, some embryos were subjected to timelapse imaging for analysis of yolk contrac-tion dynamics. The rest of the embryos were kept at 24*^o^*C and their phenotypes were analyzed at 1 day post-fertilization. The phenotypes were categorized as: 0 = no axis defect, 1 = has well developed posterior, but shows head phenotype, 2 = has two axes, 3 = has head phenotype and missing or truncated posterior, 4 = has anterior mass and some segmentation in posterior, 5 = no anterior mass, but has some segmentation in posterior, 6 = no axis, or dead.

### Timelapse imaging

Simultaneous timelapse imaging was done except when working with different fish species or strains, which were unfortunately not spawning at the same time. For simultaneous timelapse imaging, eggs and embryos were recovered from the same clutch, and samples of different conditions were incubated in separate wells of the same multi-well dish (ibidi, Germany). For experiments involving dechorionated and surgically manipulated samples, samples (including control embryos with chorion) were placed in dish coated with 1% agarose (in tap water). Individual samples were put in separate agarose wells. To minimize evaporation of water, plastic dishes containing the eggs or embryos were either (a, for 9 cm plastic dishes) covered with plastic lid that has a hole just on the field of view or (b, for multi-well plastic dishes) covered with a plastic lid and placed in a bigger plastic dish containing water. Eggs and embryos were imaged at room temperature (∼ 24*^o^*C) using either an Olympus SZX16 microscope equipped with a DP80 digital camera (Olympus) and a 1.0x (SDF-PLAPO-1XPF, Olympus, #1-SX950) or a 0.5x objective (SDF-PLAPO-0.5XPF, Olympus, #1-SX605), an Olympus SZ61 microscope equipped with a DP27 digital camera (Olympus) and a 0.5x objective (110AL0.5X-2, Olympus, #1-S819-2), an Olympus SZ61 microscope equipped with an SC50 digital camera (Olympus) and a 0.5x objective (110AL0.5X-2, Olympus, #1-S819-2), or an Olympus SZ61 microscope equipped with an SC180 digital camera (Olympus) and a 0.5x objective (110AL0.5X-2, Olympus, #1-S819-2). Timelapse images were taken using the Process Manager of cellSens Standard software (Olympus) with a temporal resolution of 10 seconds. Access to metadata of timelapse imaging is available in `DATA-ImagingTimeseriesV2` at https://github.com/PGLSanchez/yolk-contractions.

### Data analysis

To extract the timeseries of yolk perimeter, circularity, and projected area, segmentation was first done using Simple Interactive Object Extraction (SIOX) plugin (Friedland et al., 2006) in Fiji (Schindelin et al., 2012) [in Fiji: `Plugins > Segmentation > SIOX: Simple Interactive Object Extraction`]. Using one frame, multiple regions within the yolk were specified as foreground [in SIOX: `Foreground`] and segmented [in SIOX: `Segment`]. The segmentation information was then saved [in SIOX: `Save segmentator`]. This segmentation was then applied to the rest of the stack [in Fiji: `Plugins > Segmentation > Apply saved SIOX segmentator`]. A binary mask, corresponding to the yolk, was then created for each frame of the stack. The circularity, perimeter, and area of this mask over time was analyzed using a Fiji macro (.ijm) available at https://github.com/PGLSanchez/yolk-contractions, which iterates `Analyze > Measure` over all frames in a stack.

For most of the analyses, the timeseries of each sample was extracted using Fiji by specifying a circular or polygonal ROI marking the chorion and plotting the z-axis profile [in Fiji: `Image > Stacks > Z Project`], which plots the mean pixel value over time. When comparing conditions, samples with same hh:mm:ss post-fertilization (or hh:mm:ss after exposure to water) were considered which unfortunately restricted the sample numbers. Access to raw timeseries is available in `DATA-ImagingTimeseriesV2` at https://github.com/PGLSanchez/yolk-contractions.

Raw timeseries was detrended using sinc-filter detrending after specifying a cut-off period. The cut-off period was determined empirically, which was set to be 250 seconds for all analyses except when comparisons were done between goldfish, common carp, and zebrafish. Then, a cut-off period of 850 seconds was used to account for high-period rhythms in common carp and zebrafish. Detrended timeseries was subjected to continuous wavelet transform (with the Morlet wavelet as mother wavelet) using a wavelet analysis workflow (Mönke et al., 2020), which is also implemented as a Python-based standalone software available at https://github.com/tensionhead/pyBOAT (pyBOAT 0.9.11). A ridge tracing maximum wavelet power for each timepoint was detected and the instantaneous period was extracted from this ridge. A high power indicates strong correlation of the wavelet with the signal versus white noise, with wavelet power = 3 corresponding to 95% confidence interval in case of white noise (Mönke et al., 2020). The Python code used in data analysis is available as a Jupyter notebook (.ipynb) at https://github.com/PGLSanchez/yolk-contractions. This code uses Matplotlib (Hunter, 2007), NumPy (Harris et al., 2020; Van Der Walt et al., 2011), pandas (McKinney, 2010), scikit-image (Van der Walt et al., 2014), SciPy (Virtanen et al., 2020), and seaborn (Waskom, 2021).

For comparing the timeseries of embryo mean pixel value with the timeseries of yolk circularity, yolk perimeter, and yolk projected area, clustering was done on normalized timeseries using `seaborn.clustermap`.

To determine the size of surgically manipulated samples (i.e. animal and vegetal portions of equatorially bisected goldfish embryos), a polygonal ROI marking the boundary of the sample at frame = 30, around 5 mins (i.e. 290 seconds) from the start of timelapse imaging, was specified and the projected area of this ROI was then quantified [in Fiji: `Analyze > Measure`].

For plots specifying one period per sample, median period of each sample within a specified time range was considered. Time range is defined to be when conditions that are being compared all have a median wavelet power > 3. Plots were generated- and summary statistics were calculated using PlotsOfData (Postma and Goedhart, 2019) or SuperPlotsOfData (Goedhart, 2021), which are available as Shiny apps at https://huygens.science.uva.nl/PlotsOfData and https://huygens.science.uva.nl/SuperPlotsOfData/, respectively. Periods were expressed as median ± median absolute deviation (95% confidence interval of the median) when indicated in the text.

## ACKNOWLEDGEMENTS

We thank (in alphabetical order) Hsiao-Chian Chen 陳筱茜, Jong Tai Chun, Mayu A. Fukuda 福田茉由, Tetsuya Hiraiwa 平岩徹也, Asano Ishikawa 石川麻乃, Simon Knoblich, Vincent Laudet, Shu-Hua Lee 李淑華, Nunzia Limatola, Daniel Ríos Barrera, Luigia Santella, Christian Sardet, Dillan Saunders, Stephan Q. Schneider, Benjamin Steventon and lab, Stefano Davide Vianello, Jr-Kai Yu 游智凱, former and present members of the Laboratory of Aquatic Zoology at the Yilan Marine Research Station (Yilan MRS), and all members of the EcoEvoDevo Internal Group of the Institute of Cellular and Organismic Biology (ICOB), Academia Sinica for the in-depth discussions and feedback. We likewise thank (in alphabetical order) Chen-Hui Chen 陳振輝, Kuo-Chiang Hsia 夏國強, Pou-Long Kuan 關寶龍, Shu-Hua Lee 李淑華, Athira Saju, Stephan Q. Schneider, Grace Sonia, and Stefano Davide Vianello for the sharing of reagents and equipment. We are grateful to Wei-Chen Chu 朱韋臣 of the Imaging Core Facility of ICOB and to all the administrative staff and aquaculture specialists of Yilan MRS. We are also thankful to Ricardo Henriques for kindly sharing the template that was used to format this manuscript, and we are grateful to Daniel Ríos Barrera and Stefano Davide Vianello for critical reading and editing. This work was supported by an Academia Sinica Postdoctoral Scholarship (#235g), Academia Sinica Career Development Award (CDA-103-L05), a Japan Society for the Promotion of Science Grant (JSPS KAKENHI Grant JP16K18546), and Taiwan National Science and Technology Council (formerly Ministry of Science and Technology) Grants (MOST Grant 109-2311-B-001-027-MY3 and NSTC Grant 112-2311-B-001-033).

## AUTHOR CONTRIBUTIONS

**Conceptualization**: Paul Gerald Layague Sanchez (P). **Data curation**: P. **Formal Analysis**: P. **Funding Acquisition**: P, Kinya G. Ota 太田欽也 (K). **Investigation**: P, K. **Methodology**: P, Chen-Yi Wang 王貞懿 (C), Ing-Jia Li 李穎佳 (I), K. **Project Administration**: P, K. **Resources**: C, I, K. **Software**: P. **Supervision**: P, K. **Validation**: P, C, I, K. **Visualization**: P. **Writing – Original Draft**: P. **Writing – Review & Editing**: P.

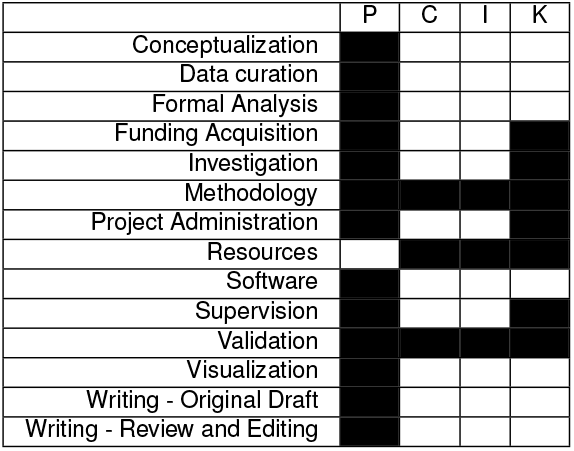

## Supplementary Note 1: Supplementary Figures

**Supplementary Figure F1: Goldfish embryos, unlike embryos of common carp and of zebrafish, exhibit persistent, fast rhythmic yolk contractions.** (**A**) Temporal evolution of period for the entire duration of the experiment, obtained from wavelet analysis of detrended timeseries (via sinc-filter detrending, cut-off period = 850 seconds). The period evolution for each sample and the median of the periods are represented as a color-coded dashed line and a color-coded solid line, respectively. The color-coded shaded area corresponds to the interquartile range. Note fast periodic rhythm in goldfish that persists for many hours. Snapshots of representative fish embryos are shown above each period evolution plot. Solid dark gray bars denote regions where median wavelet power (MWP) > 3, i.e. greater than the minimum wavelet power corresponding to 95% confidence interval in case of white noise. Shaded region corresponds to subset time duration considered for comparison of periods as in Figure 2B-C. Asterisk marks timepoint where ambient light was briefly disturbed during the timelapse imaging of common carp embryos. Similar plots are shown with the corresponding temporal evolution of wavelet power in Supplementary Figure F2A. (**B**) Snapshots of common carp embryo showing contractions shortly after fertilization. Scale bar = 500 *µ*m. (**C**) Snapshots of same common carp embryo showing contractions that last until just before the completion of the first cleavage. Scale bar = 500 *µ*m. These contractions can be seen in Supplementary MovieM3.

**Supplementary Figure F2: Rhythms recorded in goldfish, common carp, and zebrafish are not an artefact of image acquisition or are due to the background.** (**A**) Temporal evolution of wavelet power for the entire duration of the experiment, obtained from wavelet analysis of detrended timeseries (via sinc-filter detrending, cut-off period = 850 seconds). The median of the wavelet powers and the interquartile range are represented as a color-coded solid line and color-coded shaded area, respectively. Horizontal dotted line marks wavelet power = 3, which corresponds to 95% confidence interval in case of white noise. Shaded region corresponds to subset time duration considered for comparison of periods as in Figure2B-C and Supplementary Figure F1A. Asterisk marks timepoint where ambient light was briefly disturbed during the timelapse imaging of common carp embryos. (**B**) Detrended timeseries and wavelet analysis of regions corresponding to background in close proximity to analyzed fish embryos. Number IDs denote clutch ID of fish embryos. Note low wavelet power for background regions, especially when compared with that for embryos as shown in Panel A and Figure 2B.

**Supplementary Figure F3: Phenotypes of drug-treated embryos at 1 day post-fertilization** (**A**) Representative images of surviving control and treated embryos showing the different phenotypes at 1 day post-fertilization: 0 = no axis defect, 1 = has well developed posterior, but shows head phenotype, 2 = has two axes, 3 = has head phenotype and missing or truncated posterior, 4 = has anterior mass and some segmentation in posterior, 5 = no anterior mass, but has some segmentation in posterior, and 6 = no axis. Scale bar = 500 *µ*m. (**B**) Phenotypes of treated embryos and their respective controls at 1 day post-fertilization. For experiment with nocodazole (microtubule depolymerizing drug), embryos were incubated for ∼ 4 mins in 1 *µ*g/mL nocodazole (N, n = 140) or 1:20000 DMSO (D, n = 123) control at ∼ 10 mins post-fertilization. For experiment with latrunculin (actin polymerization inhbitor), embryos were incubated in 100 nM latrunculin A (L, n = 144) or 1:10000 DMSO (D, n = 147) control from ∼ 10 mins post-fertilization. For experiment with blebbistatin(–) (myosin inhibitor), embryos were incubated in 10 *µ*M blebbistatin(–) (B–, n = 10) or 10 *µ*M blebbistatin(+) (B+, n = 12) control from ∼ 3 hours post-fertilization. For experiment with double distilled water (ddH_2_O, Ca^2+^ free water) and EDTA (Ca^2+^ chelating agent), embryos were incubated in ddH_2_O (W, n = 22), 1 mM EDTA in ddH_2_O (E, n = 24), or tap water (T, n = 13) control from ∼ 3 hours post-fertilization. Due to unavailability of spawning wild-type (ZWJ-*ChdS*(+/+) strain) goldfish, ZWJ-*ChdS*(–/–)-singletail goldfish was used for experiments with blebbistatin, ddH_2_O, and EDTA.

**Supplementary Figure F4: Rhythmic contractions persist even in yolk that is pinched off from dying embryos. (A)** Snapshots (top row) and magnified images (bottom row) of yolk that is getting pinched off from a dying embryo over time (06:07:10–06:08:30 hh:mm:ss post-fertilization). Embryo was treated with 0.1 *µ*g/mL nocodazole for ∼ 4 mins at ∼ 10 mins post-fertilization. Magenta arrowheads mark pinching off of the yolk. Scale bar = 500 *µ*m. (**B**) Snapshots (top row) and magnified images (bottom row) of persistent contractions of yolk that is pinched off from a dying embryo (same sample as in panel A) over time (06:13:00–06:17:00 hh:mm:ss post-fertilization). Magenta symbols mark change in the shape of the pinched off yolk between consecutive snapshots. Scale bar = 500 *µ*m. For more samples showing persistent contractions of yolk that is pinched off from a dying embryo, refer to Supplementary Movie M9.

**Fig. F1.**
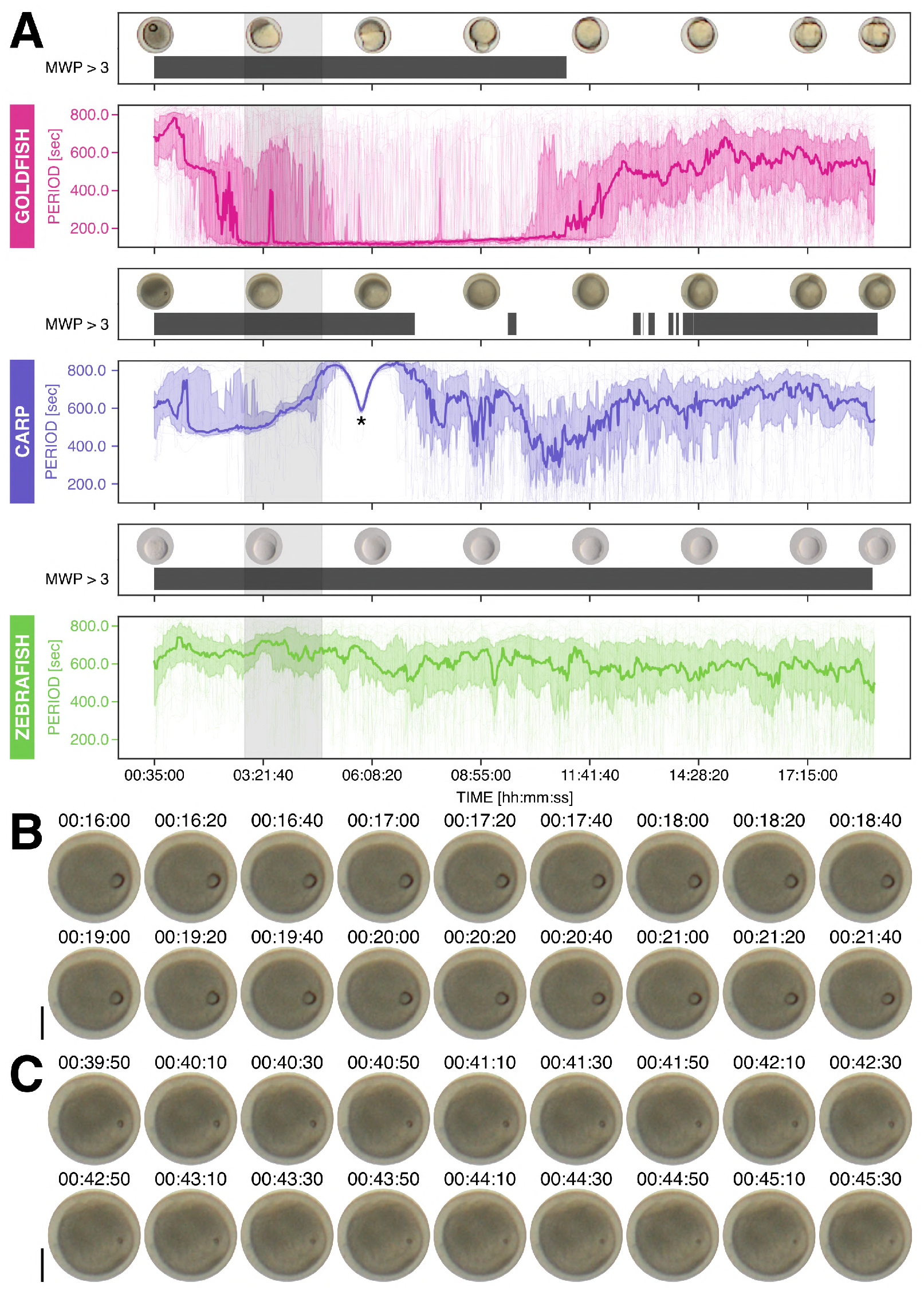
Goldfish embryos, unlike embryos of common carp and of zebrafish, exhibit persistent, fast rhythmic yolk contractions. (**A**) Temporal evolution of period for the entire duration of the experiment, obtained from wavelet analysis of detrended timeseries (via sinc-filter detrending, cut-off period = 850 seconds). The period evolution for each sample and the median of the periods are represented as a color-coded dashed line and a color-coded solid line, respectively. The color-coded shaded area corresponds to the interquartile range. Note fast periodic rhythm in goldfish that persists for many hours. Snapshots of representative fish embryos are shown above each period evolution plot. Solid dark gray bars denote regions where median wavelet power (MWP) > 3, i.e. greater than the minimum wavelet power corresponding to 95% confidence interval in case of white noise. Shaded region corresponds to subset time duration considered for comparison of periods as in Figure 2B-C. Asterisk marks timepoint where ambient light was briefly disturbed during the timelapse imaging of common carp embryos. Similar plots are shown with the corresponding temporal evolution of wavelet power in Supplementary Figure F2A. **(B)** Snapshots of common carp embryo showing contractions shortly after fertilization. Scale bar = 500 *µ*m. (**C**) Snapshots of same common carp embryo showing contractions that last until just before the completion of the first cleavage. Scale bar = 500 *µ*m. These contractions can be seen in Supplementary Movie M3.

**Fig. F2.**
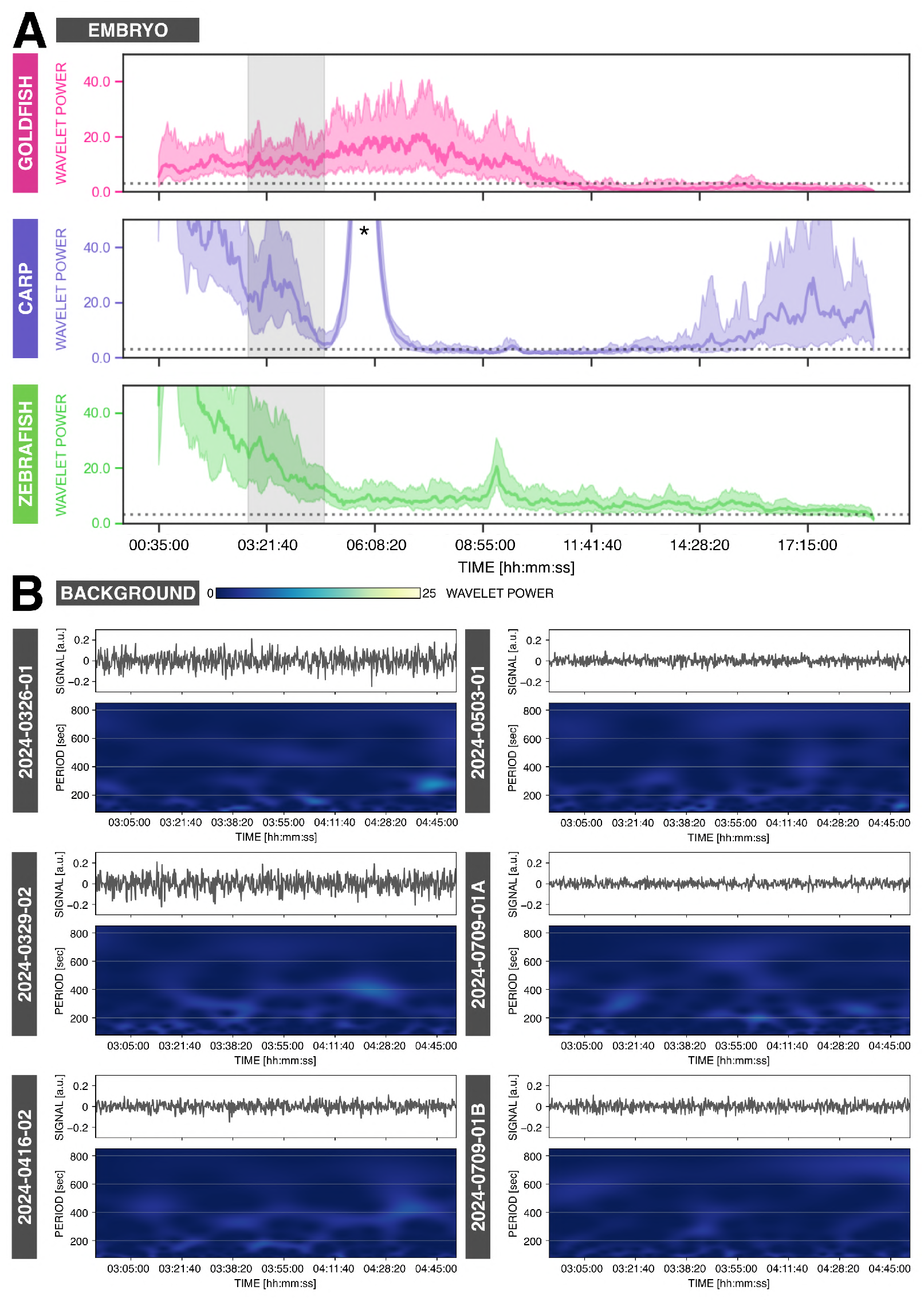
Rhythms recorded in goldfish, common carp, and zebrafish are not an artefact of image acquisition or are due to the background. (**A**) Temporal evolution of wavelet power for the entire duration of the experiment, obtained from wavelet analysis of detrended timeseries (via sinc-filter detrending, cut-off period = 850 seconds). The median of the wavelet powers and the interquartile range are represented as a color-coded solid line and color-coded shaded area, respectively. Horizontal dotted line marks wavelet power = 3, which corresponds to 95% confidence interval in case of white noise. Shaded region corresponds to subset time duration considered for comparison of periods as in Figure 2B-C and Supplementary Figure F1A. Asterisk marks timepoint where ambient light was briefly disturbed during the timelapse imaging of common carp embryos. (**B**) Detrended timeseries and wavelet analysis of regions corresponding to background in close proximity to analyzed fish embryos. Number IDs denote clutch ID of fish embryos. Note low wavelet power for background regions, especially when compared with that for embryos as shown in Panel A and Figure 2B.

**Fig. F3.**
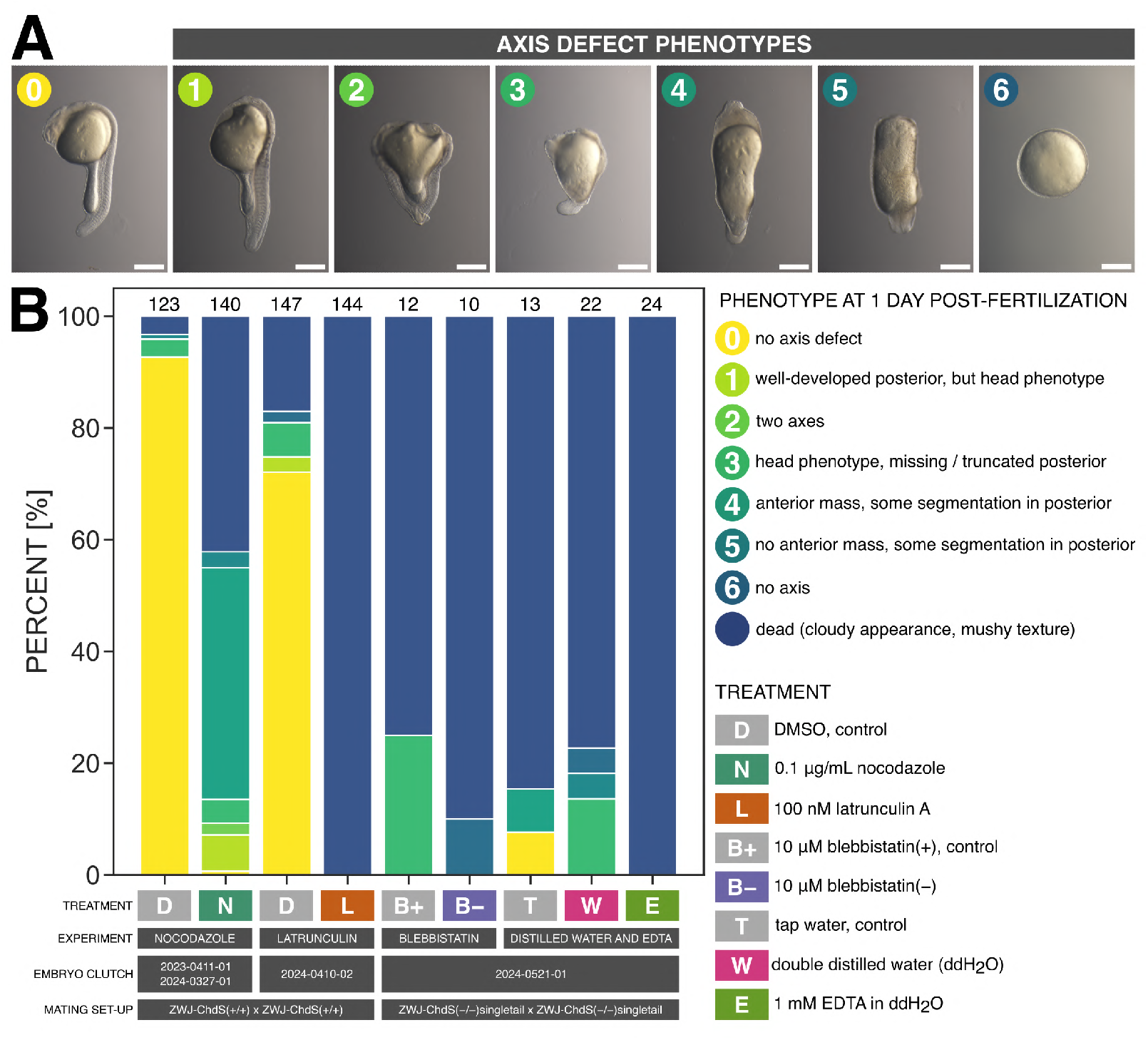
Phenotypes of drug-treated embryos at 1 day post-fertilization. (**A**) Representative images of surviving control and treated embryos showing the different phenotypes at 1 day post-fertilization: 0 = no axis defect, 1 = has well developed posterior, but shows head phenotype, 2 = has two axes, 3 = has head phenotype and missing or truncated posterior, 4 = has anterior mass and some segmentation in posterior, 5 = no anterior mass, but has some segmentation in posterior, and 6 = no axis. Scale bar = 500 *µ*m. (**B**) Phenotypes of treated embryos and their respective controls at 1 day post-fertilization. For experiment with nocodazole (microtubule depolymerizing drug), embryos were incubated for *∼* 4 mins in 1 *µ*g/mL nocodazole (N, n = 140) or 1:20000 DMSO (D, n = 123) control at *∼* 10 mins post-fertilization. For experiment with latrunculin (actin polymerization inhbitor), embryos were incubated in 100 nM latrunculin A (L, n = 144) or 1:10000 DMSO (D, n = 147) control from *∼* 10 mins post-fertilization. For experiment with blebbistatin(–) (myosin inhibitor), embryos were incubated in 10 *µ*M blebbistatin(–) (B–, n = 10) or 10 *µ*M blebbistatin(+) (B+, n = 12) control from *∼* 3 hours post-fertilization. For experiment with double distilled water (ddH_2_O, Ca^2+^ free water) and EDTA (Ca^2+^ chelating agent), embryos were incubated in ddH_2_O (W, n = 22), 1 mM EDTA in ddH_2_O (E, n = 24), or tap water (T, n = 13) control from *∼* 3 hours post-fertilization. Due to unavailability of spawning wild-type (ZWJ-*ChdS*(+/+) strain) goldfish, ZWJ-*ChdS*(–/–)-singletail goldfish was used for experiments with blebbistatin, ddH_2_O, and EDTA.

**Fig. F4.**
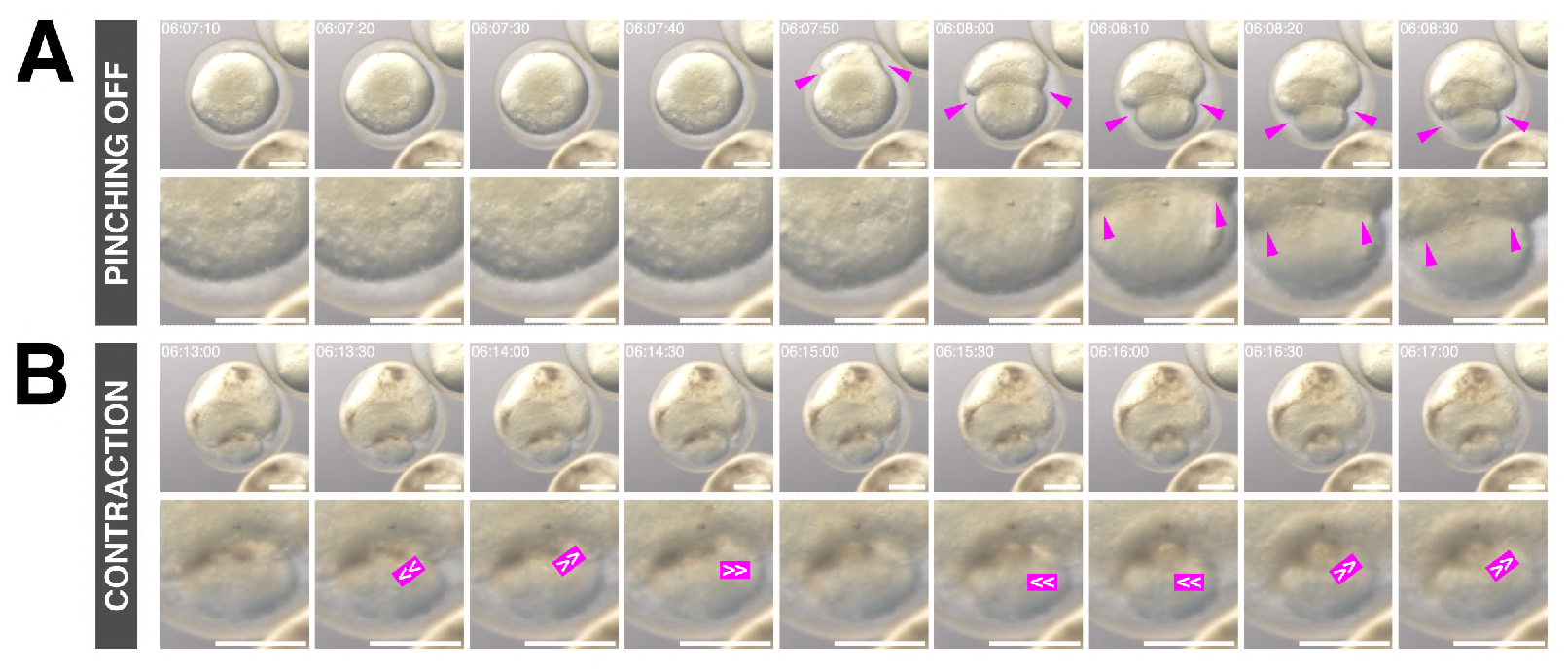
Rhythmic contractions persist even in yolk that is pinched off from dying embryos. (**A**) Snapshots (top row) and magnified images (bottom row) of yolk that is getting pinched off from a dying embryo over time (06:07:10–06:08:30 hh:mm:ss post-fertilization). Embryo was treated with 0.1 *µ*g/mL nocodazole for *∼* 4 mins at *∼* 10 mins post-fertilization. Magenta arrowheads mark pinching off of the yolk. Scale bar = 500 *µ*m. (**B**) Snapshots (top row) and magnified images (bottom row) of persistent contractions of yolk that is pinched off from a dying embryo (same sample as in panel A) over time (06:13:00–06:17:00 hh:mm:ss post-fertilization). Magenta symbols mark change in the shape of the pinched off yolk between consecutive snapshots. Scale bar = 500 *µ*m. For more samples showing persistent contractions of yolk that is pinched off from a dying embryo, refer to Supplementary Movie M9.

## Supplementary Note 2: Supplementary Movies

All the supplementary movies can be accessed via this playlist:Goldfish Yolk Contractions (v2) – Supplementary Movies.

**Supplementary Movie M1: Goldfish embryos exhibit rhythmic yolk contractions from the 4-cell stage to the gastrula (epiboly) stage.** Timelapse of goldfish embryos at different stages of development: zygote to cleavage stage at 00:21:00–01:43:00 hh:mm:ss post-fertilization, cleavage to blastula stage at 01:45:00–02:34:50 hh:mm:ss post-fertilization, blastula stage at 03:13:00–04:17:40 hh:mm:ss post-fertilization and at 04:31:00–05:10:10 hh:mm:ss post-fertilization, and gastrula (epiboly) stage at 07:28:30–09:06:10 hh:mm:ss post-fertilization. Scale bar = 500 *µ*m. Movie is available at https://youtu.be/96XY7i88lmY. See also our video on early embryonic development of goldfish available at https://youtu.be/TVMull5YEqw.

**Supplementary Movie M2: Extracting the mean pixel value of an embryo over time allows quantification of yolk contractions.** Timelapse of goldfish embryo and its yolk (after image segmentation) with raw timeseries of yolk circularity, yolk perimeter, yolk projected area, and embryo mean pixel value. Scale bar = 500 *µ*m. Embryo is the same as embryo at cleavage-blastula stage in Supplementary Movie M1. Movie is available at https://youtu.be/U7OM27 _3_*sQQ*.

**Supplementary Movie M3: Persistent and rhythmic contractions of the yolk are present in goldfish but not in closely-related common carp or zebrafish.** Timelapse of embryos of goldfish *Carassius auratus*, common carp *Cyprinus carpio*, and zebrafish *Danio rerio* at room temperature. Time is in hh:mm:ss post-fertilization. Scale bar = 500 *µ*m. Note periodic contractions of the yolk in goldfish that lasts for many hours. Also note contractions in common carp that starts shortly after fertilization but lasts just until around the first cleavage (from around 00:15:00 to around 00:45:00 hh:mm:ss post-fertilization). Movie is available at https://youtu.be/8KHQLgLWK1s.

**Supplementary Movie M4: The rhythmic contractions of the goldfish yolk are independent from fertilization or cell division, and emerge at a precise time.** (**First movie**) Timelapse of fertilized and unfertilized goldfish eggs at different time periods: cleavage stage at 00:33:00–01:53:40 hh:mm:ss post-exposure of egg (and sperm, for fertilized sample) to water, cleavage-blastula stage at 01:45:00–02:00:40 hh:mm:ss post-exposure of egg (and sperm, for fertilized sample) to water, and gastrula stage at 07:58:00–08:03:20 hh:mm:ss post-exposure of egg (and sperm, for fertilized sample) to water. Labels for each time period are based on developmental staging of fertilized samples. Scale bar = 500 *µ*m. Note emergence of rhythmic yolk contractions in both samples from around 01:30:00 hh:mm:ss post-exposure of egg (and sperm, for fertilized sample), when the fertilized sample is at 4-cell stage. Also note failure of unfertilized samples to undergo cell divisions. The fertilized sample at cleavage-blastula stage is the same as the goldfish embryo at cleavage-blastula stage in Supplementary Movie M1. The fertilized sample at gastrula stage is the same as the goldfish sample (GF x GF) in Supplementary Movie M15. (**Second movie**, in thumbnail) Simultaneous timelapse of fertilized (left) and unfertilized (right) goldfish eggs. Time is in hh:mm:ss post-exposure of egg (and sperm, for fertilized sample) to water. Scale bar = 2 mm. Movies available at https://youtu.be/hD8lzYZ2UQg.

**Supplementary Movie M5: The rhythmic contractions of the yolk persist even in dechorionated goldfish embryos.** Simultaneous timelapse of goldfish embryos with (left) or without (right) chorion, both incubated in agarose-coated dish. Time is in hh:mm:ss post-fertilization. Scale bar = 2 mm. Note more apparent yolk deformations in dechorionated embryos and what could be hydrodynamic interaction between these samples. Movie is available at https://youtu.be/EWuEfUnnvpA.

**Supplementary Movie M6: The rhythmic contractions of the yolk persist even in samples where the animal and vegetal poles are separated.** Left: Equatorial bisection of goldfish embryo using an eyelash tool to separate the animal and vegetal poles. Right-top: Isolated yolk (vegetal portion) showing persistent rhythmic movements that manifest as apparent rotations. Time is in hh:mm:ss post-fertilization. Scale bar = 1 mm. Right-bottom: Pair of animal and vegetal portions both showing persistent rhythmic movements. Movie is available at https://youtu.be/6HH3tdj-gaM.

**Supplementary Movie M7: The rhythmic contractions are an emergent property of the goldfish yolk, and are independent of the chorion and of the embryo proper (in the animal pole).** (**First movie**, in thumbnail) Simultaneous timelapse of intact (i.e. with and without chorion) and bisected (i.e. vegetal and animal portions) goldfish embryos in agarose wells. Time is in hh:mm:ss post-fertilization. Scale bar = 2 mm. Note more apparent yolk deformations in intact dechorionated embryos, apparent rotations in vegetal portions (i.e. isolated yolk), and persistent yolk contractions in animal portions (i.e. small amount of yolk with embryo proper). Right: Representative samples for each condition. Samples are the same as those in Figure 4A. (**Second movie**) Left: Simultaneous timelapse of representative intact (i.e. with and without chorion) and bisected (i.e. vegetal and animal portions) goldfish embryos in agarose wells. Right: Corresponding detrended timeseries of representative samples (via sinc-filter detrending, cut-off period = 250 seconds). Time is in seconds post-fertilization. Movies available at https://youtu.be/FmBmOlxjQvM.

**Supplementary Movie M8: Rhythmic contractions of the yolk in goldfish embryos acutely treated with a microtubule depolymerizing drug, nocodazole.** Simultaneous timelapse of goldfish embryos incubated in either 0.1 *µ*g/mL nocodazole (right) or DMSO control (left) for ∼ 4 mins at ∼ 10 mins post-fertilization. Time is in hh:mm:ss post-fertilization. Scale bar = 2 mm. Movie is available at https://youtu.be/Vnox05QuRJA.

**Supplementary Movie M9: The rhythmic contractions of the goldfish yolk persist even in yolk that is pinched off from dying embryos.** Timelapse of goldfish embryos showing persistent contractions of yolk that is pinched off from embryo. Embryos were treated with 0.1 *µ*g/mL nocodazole for ∼ 4 mins at ∼ 10 mins post-fertilization. White arrows mark samples showing persistent yolk contractions of pinched-off yolk. Specified numbers are representative timepoints when these contractions are visible. Time is in hh:mm:ss post-fertilization. Scale bar = 500 *µ*m. Movie is available at https://youtu.be/Fm296U5thaM.

**Supplementary Movie M10: Rhythmic contractions of the yolk in goldfish embryos treated with an actin polymerization inhibitor, latrunculin.** Simultaneous timelapse of goldfish embryos incubated in either 100 nM latrunculin A (right) or DMSO control (left) from ∼ 10 mins post-fertilization. Time is in hh:mm:ss post-fertilization. Scale bar = 2 mm. Movie is available at https://youtu.be/1-MOZORp3z0.

**Supplementary Movie M11: Rhythmic contractions of the yolk in goldfish embryos treated with a myosin inhibitor, blebbistatin(–).** Simultaneous timelapse of goldfish embryos incubated in either 10 *µ*M blebbistatin(–) (right) or 10 *µ*M blebbistatin(+) control (left) from ∼ 3 hours post-fertilization. Top row: Simultaneous timelapse until 03:00:00 hh:mm:ss post-fertilization, before treatment. Bottom row: Simultaneous timelapse from 03:15:00 hh:mm:ss post-fertilization, during treatment. Time is in hh:mm:ss post-fertilization. Scale bar = 2 mm. Due to unavailability of spawning wild-type (ZWJ-*ChdS*(+/+) strain) goldfish, ZWJ-*ChdS*(–/–)-singletail goldfish was used. Movie is available at https://youtu.be/ZkxuXLxUj *_A_*.

**Supplementary Movie M12: Rhythmic contractions of the yolk in goldfish embryos treated with Ca** ^2+^ **free water or Ca**^2+^ **chelating agent, EDTA.** Simultaneous timelapse of goldfish embryos incubated in either double distilled water (ddH_2_O, middle), 1 mM EDTA in ddH_2_O (right), or tap water control (left) from ∼ 3 hours post-fertilization. Top row: Simultaneous timelapse until 03:00:00 hh:mm:ss post-fertilization, before treatment. Bottom row: Simultaneous timelapse from 03:15:00 hh:mm:ss post-fertilization, during treatment. Time is in hh:mm:ss post-fertilization. Scale bar = 2 mm. Due to unavailability of spawning wild-type (ZWJ-*ChdS*(+/+) strain) goldfish, ZWJ-*ChdS*(–/–)-singletail goldfish was used. Movie is available at https://youtu.be/Ar1B1rbL6L0.

**Supplementary Movie M13: Rhythmic contractions of the yolk in twin-tail goldfish embryos are faster than those in wild-type.** Left: Timelapse of wild-type (top) and twin-tail (i.e. *Oranda*, bottom) goldfish embryos. Time is in hh:mm:ss post-fertilization. Scale bar = 2 mm. Simultaneous imaging was unfortunately not possible because the two goldfish strains were not spawning at the same time. Right: Temporal evolution of period of yolk contractions for the entire duration of the experiment, obtained from wavelet analysis of detrended timeseries (via sinc-filter detrending, cut-off period = 250 seconds). The period evolution for each sample and the median of the periods are represented as a color-coded dashed line and a color-coded solid line, respectively. The color-coded shaded area corresponds to the interquartile range. Snapshots of representative fish embryos are shown above each plot. Median wavelet power is also plotted in gray, with a horizontal dotted gray line marking wavelet power threshold = 3, which corresponds to 95% confidence interval in case of white noise. Note shorter (i.e. faster) period of yolk contractions in twin-tail *Oranda* goldfish embryos. These data are the same as those plotted in Figure 6C. Movie is available at https://youtu.be/XZ7K8HKpS0Y.

**Supplementary Movie M14: Rhythmic contractions of the yolk in goldfish embryos with different *ChdS* geno-types.** Left: Simultaneous timelapse of goldfish embryos from mating of ZWJ-*ChdS*(+/–) goldfish. Numbers denote sample ID for genotyping and X marks samples that were recovered but were not genotyped (due to loss during handling). Time is in hh:mm:ss post-fertilization. Scale bar = 2 mm. Right: *ChdS* genotyping of samples in the left. Embryos were recovered at 1 day post-fertilization, after timelapse imaging, and were subsequently genotyped from 3 days post-fertilization. For genotyping, ∼ 400 bp of the *ChdS* gene was amplified and then treated with the Eco88I restriction enzyme, which recognizes a restriction enzyme site in the *ChdS*(–) allele resulting in a shorter ∼ 300 bp (and ∼ 100 bp) band (Abe et al.,2014). Movie is available at https://youtu.be/nVYwIuxD964.

**Supplementary Movie M15: The rhythmic contraction of the yolk is a trait that is maternal in origin.** Timelapse of embryos of GF x GF goldfish, CP x CP common carp, and their hybrids: (1) CP x GF from common carp sperm and goldfish egg and (2) GF x CP from goldfish sperm and common carp egg. Time is in hh:mm:ss post-fertilization. Scale bar = 500 *µ*m. Samples are the same as those in Figure 8B. Note rhythmic contractions of the yolk in embryos where the egg had been derived from goldfish. Movie is available at https://youtu.be/8l5PfF2uOjE.

**Fig. M1.**
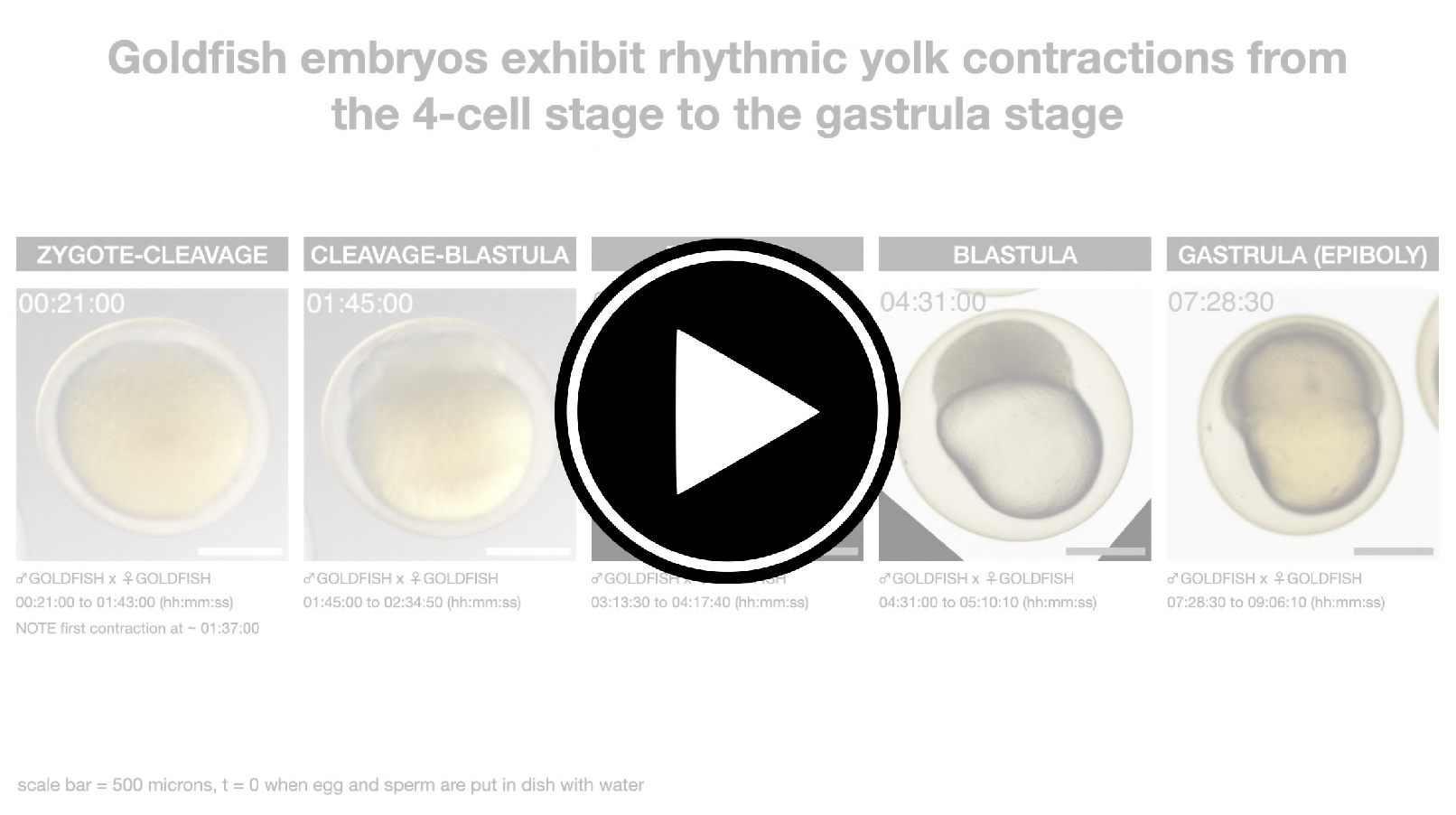
Goldfish embryos exhibit rhythmic yolk contractions from the 4-cell stage to the gastrula (epiboly) stage. Timelapse of goldfish embryos at different stages of development: zygote to cleavage stage at 00:21:00–01:43:00 hh:mm:ss post-fertilization, cleavage to blastula stage at 01:45:00–02:34:50 hh:mm:ss post-fertilization, blastula stage at 03:13:00–04:17:40 hh:mm:ss post-fertilization and at 04:31:00–05:10:10 hh:mm:ss post-fertilization, and gastrula (epiboly) stage at 07:28:30–09:06:10 hh:mm:ss post-fertilization. Scale bar = 500 *µ*m. Movie is available at https://youtu.be/96XY7i88lmY. See also our video on early embryonic development of gold-fish available at https://youtu.be/TVMull5YEqw.

**Fig. M2.**
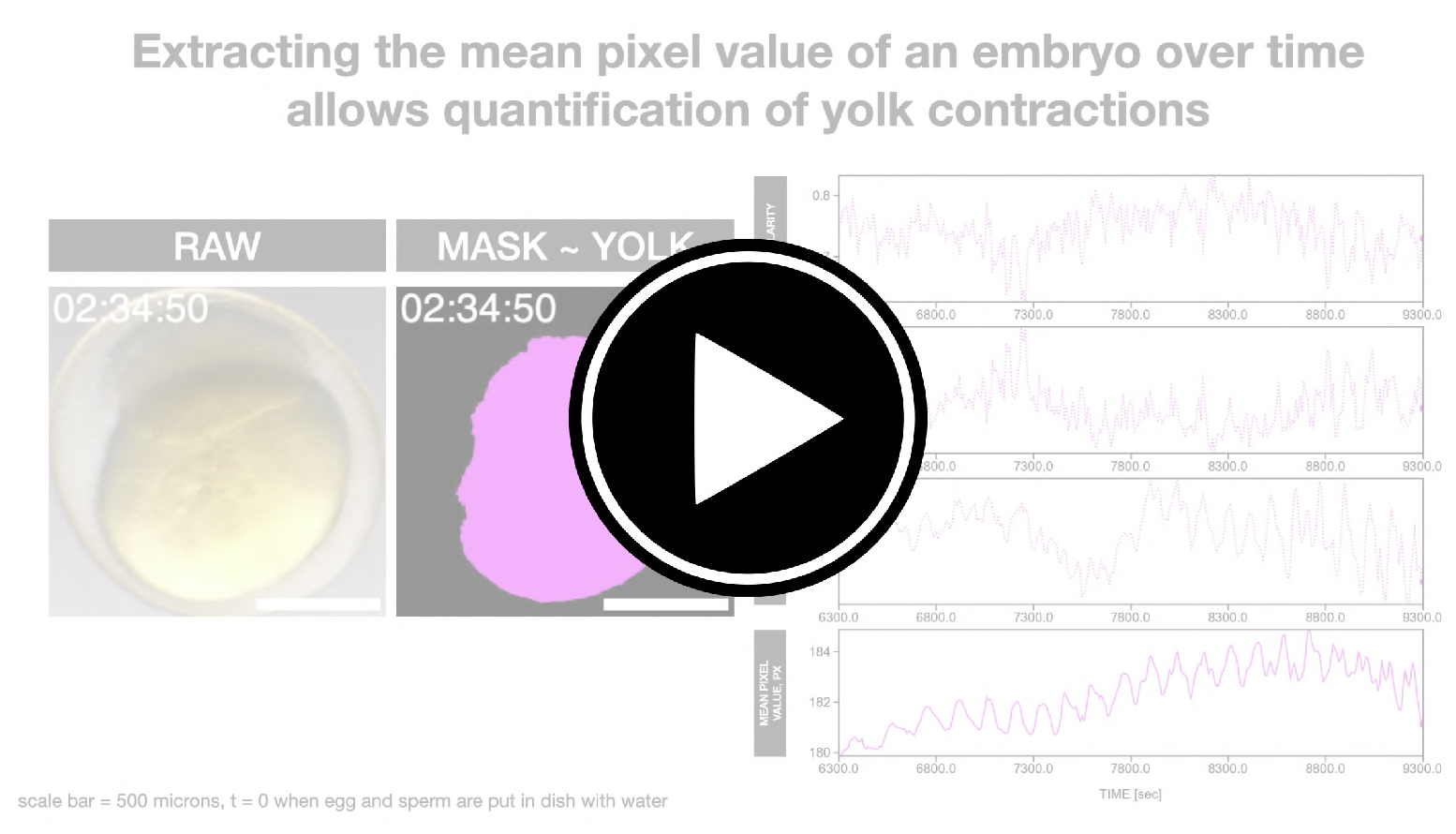
Extracting the mean pixel value of an embryo over time allows quantification of yolk contractions. Timelapse of goldfish embryo and its yolk (after image segmentation) with raw timeseries of yolk circularity, yolk perimeter, yolk projected area, and embryo mean pixel value. Scale bar = 500 *µ*m. Embryo is the same as embryo at cleavage-blastula stage in Supplementary Movie M1. Movie is available at https://youtu.be/U7OM27 _3_*sQQ*.

**Fig. M3.**
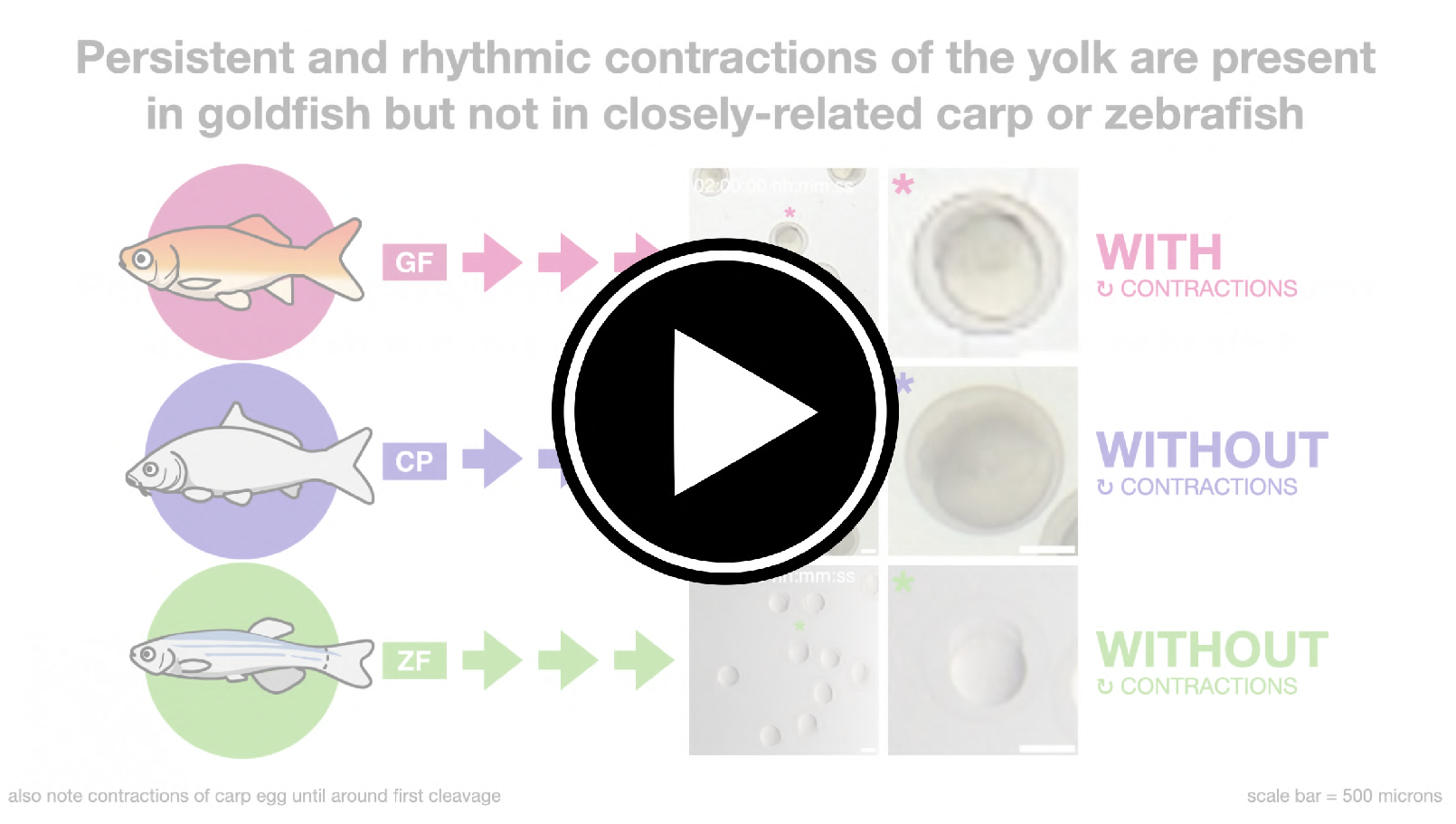
Persistent and rhythmic contractions of the yolk are present in goldfish but not in closely-related common carp or zebrafish. Timelapse of embryos of goldfish *Carassius auratus*, common carp *Cyprinus carpio*, and zebrafish *Danio rerio* at room temperature. Time is in hh:mm:ss post-fertilization. Scale bar = 500 *µ*m. Note periodic contractions of the yolk in goldfish that lasts for many hours. Also note contractions in common carp that starts shortly after fertilization but lasts just until around the first cleavage (from around 00:15:00 to around 00:45:00 hh:mm:ss post-fertilization). Movie is available at https://youtu.be/8KHQLgLWK1s.

**Fig. M4.**
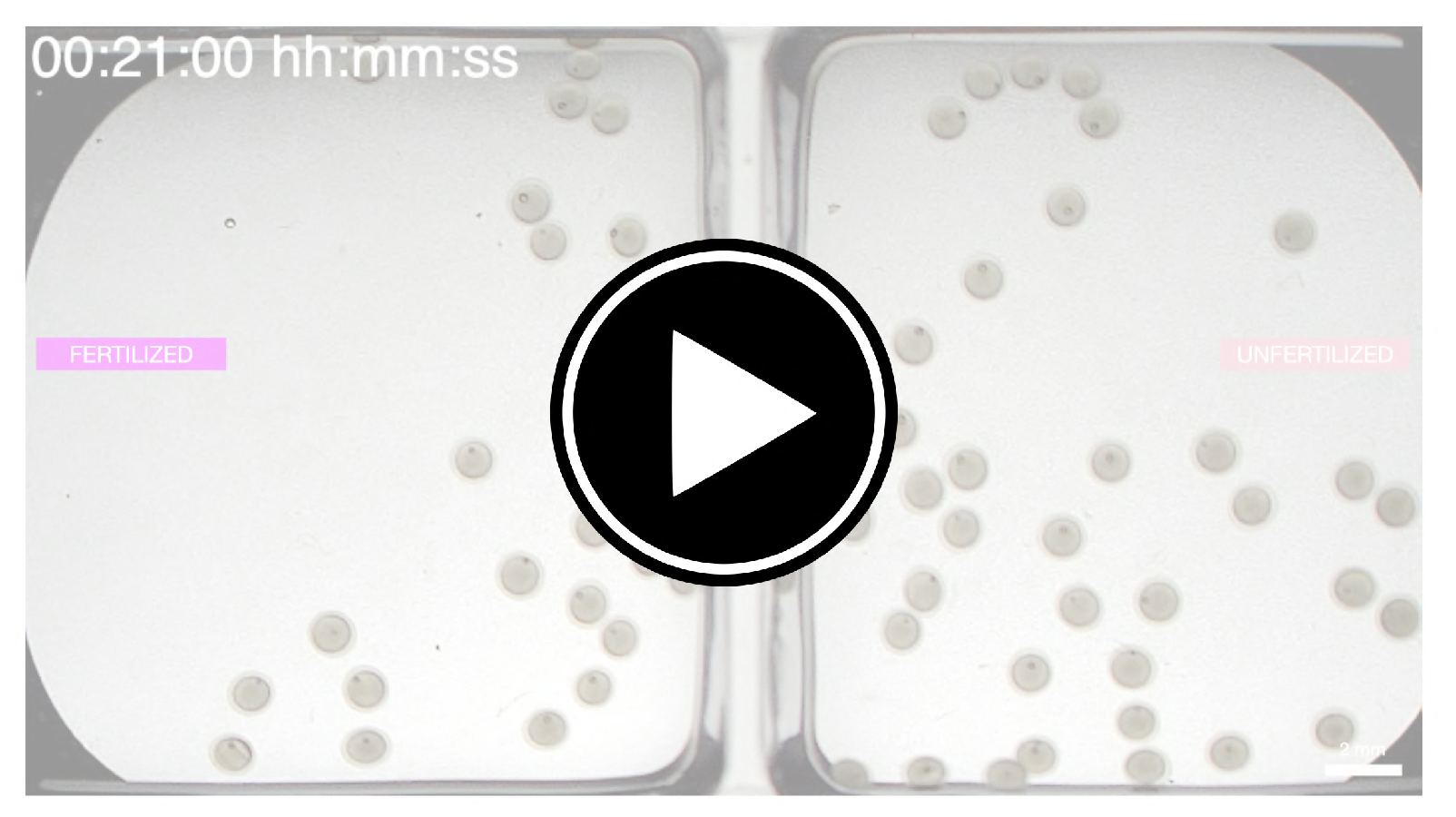
The rhythmic contractions of the goldfish yolk are independent from fertilization or cell division, and emerge at a precise time. (**First movie**) Timelapse of fertilized and unfertilized goldfish eggs at different time periods: cleavage stage at 00:33:00– 01:53:40 hh:mm:ss post-exposure of egg (and sperm, for fertilized sample) to water, cleavage-blastula stage at 01:45:00–02:00:40 hh:mm:ss post-exposure of egg (and sperm, for fertilized sample) to water, and gastrula stage at 07:58:00–08:03:20 hh:mm:ss post-exposure of egg (and sperm, for fertilized sample) to water. Labels for each time period are based on developmental staging of fertilized samples. Scale bar = 500 *µ*m. Note emergence of rhythmic yolk contractions in both samples from around 01:30:00 hh:mm:ss post-exposure of egg (and sperm, for fertilized sample), when the fertilized sample is at 4-cell stage. Also note failure of unfertilized samples to undergo cell divisions. The fertilized sample at cleavage-blastula stage is the same as the goldfish embryo at cleavageblastula stage in Supplementary MovieM1. The fertilized sample at gastrula stage is the same as the goldfish sample (GF x GF) in Supplementary MovieM 15. (**Second movie**, in thumbnail) Simultaneous timelapse of fertilized (left) and unfertilized (right) goldfish eggs. Time is in hh:mm:ss post-exposure of egg (and sperm, for fertilized sample) to water. Scale bar = 2 mm. Movies available at https://youtu.be/hD8lzYZ2UQg.

**Fig. M5.**
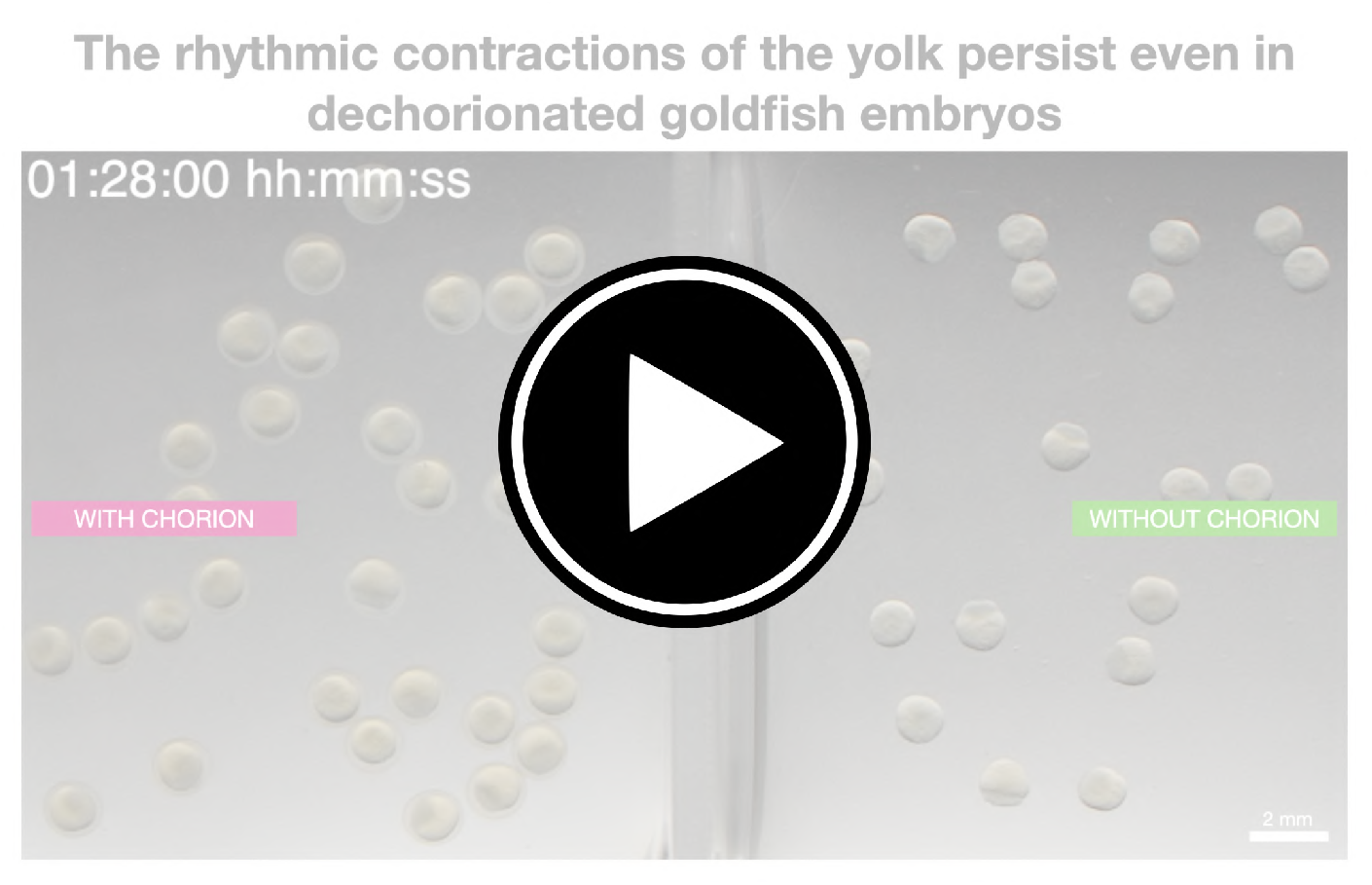
The rhythmic contractions of the yolk persist even in dechorionated goldfish embryos. Simultaneous timelapse of goldfish embryos with (left) or without (right) chorion, both incubated in agarose-coated dish. Time is in hh:mm:ss post-fertilization. Scale bar = 2 mm. Note more apparent yolk deformations in dechorionated embryos and what could be hydrodynamic interaction between these samples. Movie is available at https://youtu.be/EWuEfUnnvpA.

**Fig. M6.**
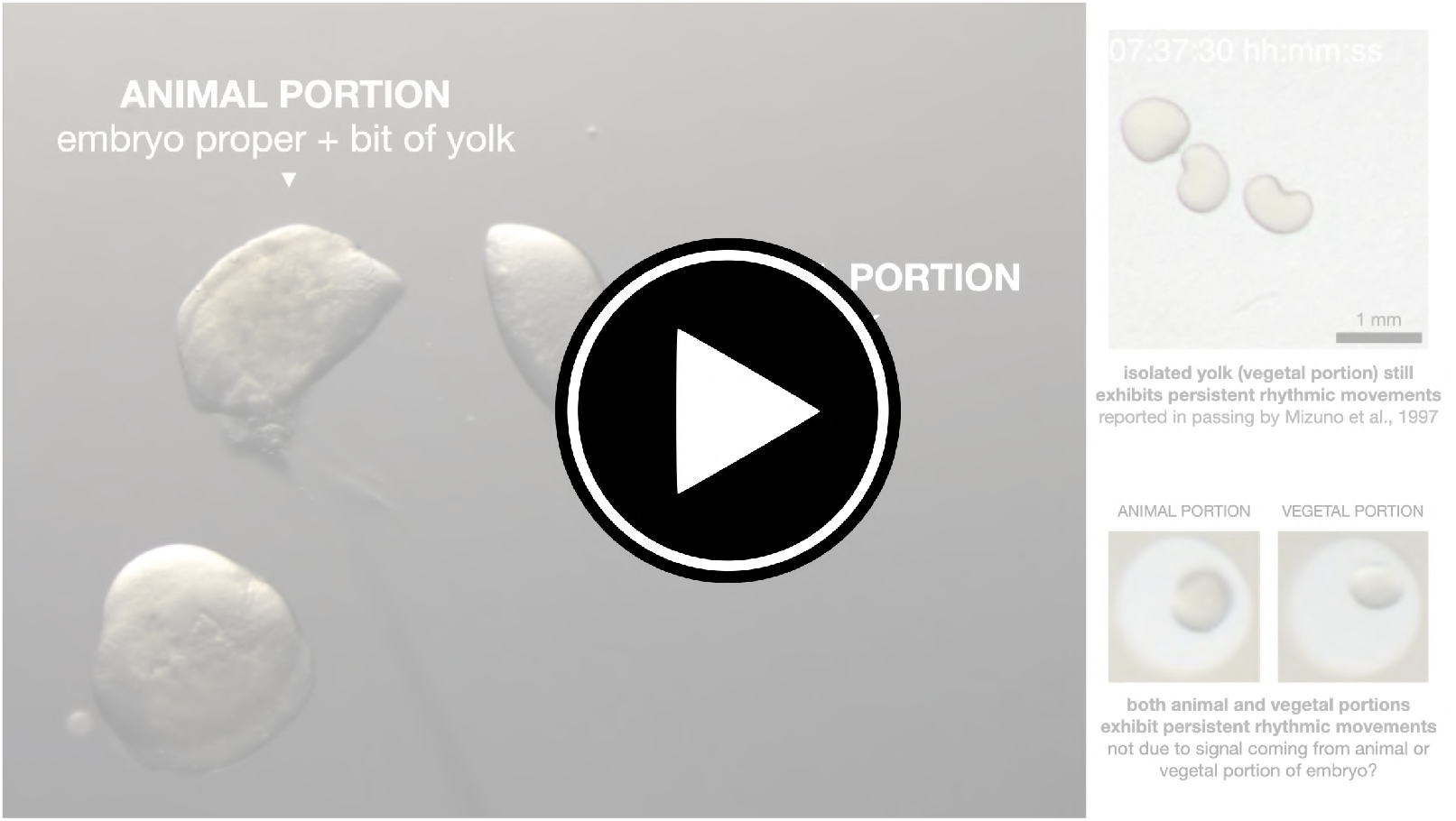
The rhythmic contractions of the yolk persist even in samples where the animal and vegetal poles are separated. Left: Equatorial bisection of goldfish embryo using an eyelash tool to separate the animal and vegetal poles. Right-top: Isolated yolk (vegetal portion) showing persistent rhythmic movements that manifest as apparent rotations. Time is in hh:mm:ss post-fertilization. Scale bar = 1 mm. Right-bottom: Pair of animal and vegetal portions both showing persistent rhythmic movements. Movie is available at https://youtu.be/6HH3tdj-gaM.

**Fig. M7.**
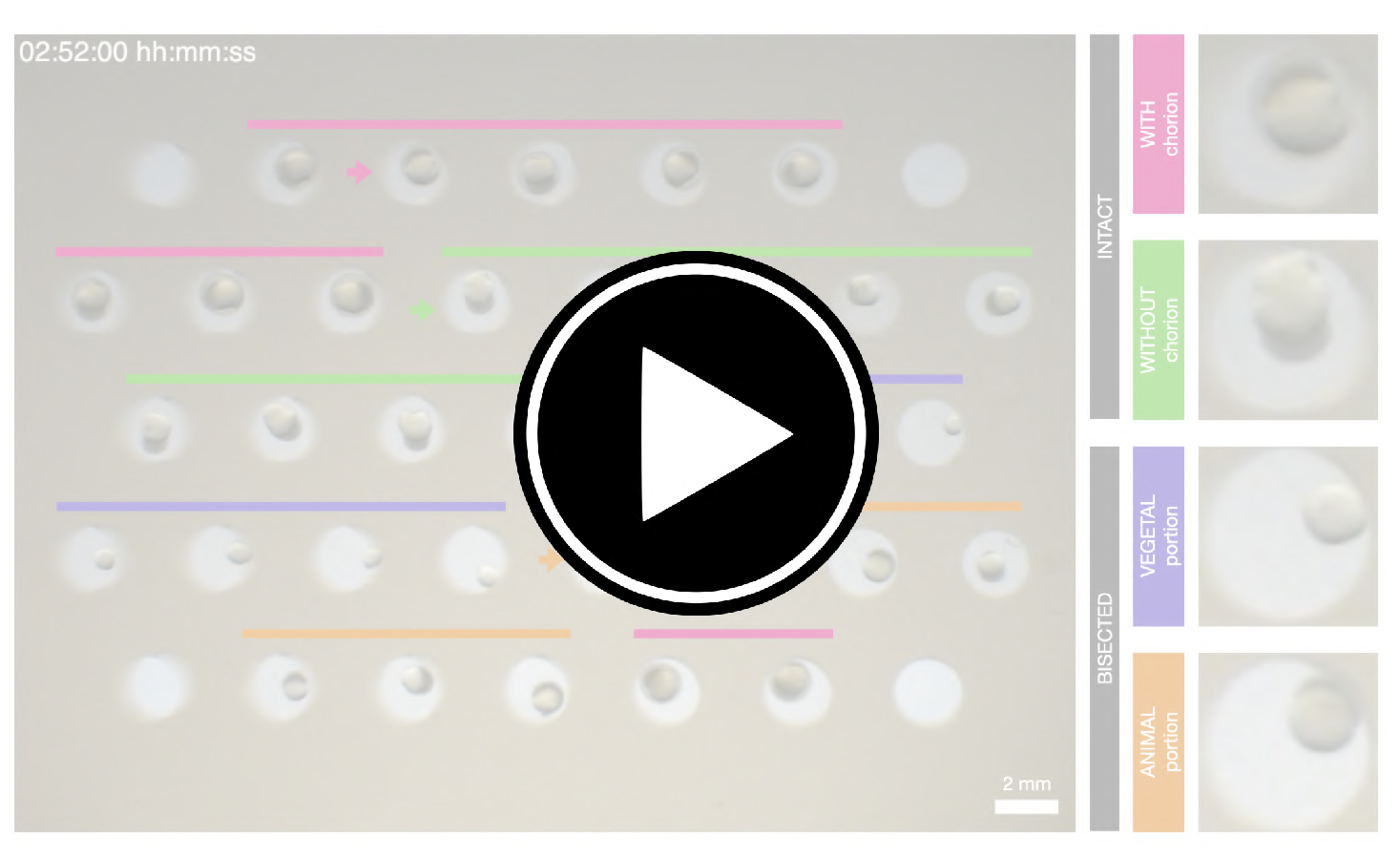
The rhythmic contractions are an emergent property of the goldfish yolk, and are independent of the chorion and of the embryo proper (in the animal pole). (**First movie**, in thumbnail) Simultaneous timelapse of intact (i.e. with and without chorion) and bisected (i.e. vegetal and animal portions) goldfish embryos in agarose wells. Time is in hh:mm:ss post-fertilization. Scale bar = 2 mm. Note more apparent yolk deformations in intact dechorionated embryos, apparent rotations in vegetal portions (i.e. isolated yolk), and persistent yolk contractions in animal portions (i.e. small amount of yolk with embryo proper). Right: Representative samples for each condition. Samples are the same as those in Figure 4A. (**Second movie**) Left: Simultaneous timelapse of representative intact (i.e. with and without chorion) and bisected (i.e. vegetal and animal portions) goldfish embryos in agarose wells. Right: Corresponding detrended timeseries of representative samples (via sinc-filter detrending, cut-off period = 250 seconds). Time is in seconds post-fertilization. Movies available at https://youtu.be/FmBmOlxjQvM.

**Fig. M8.**
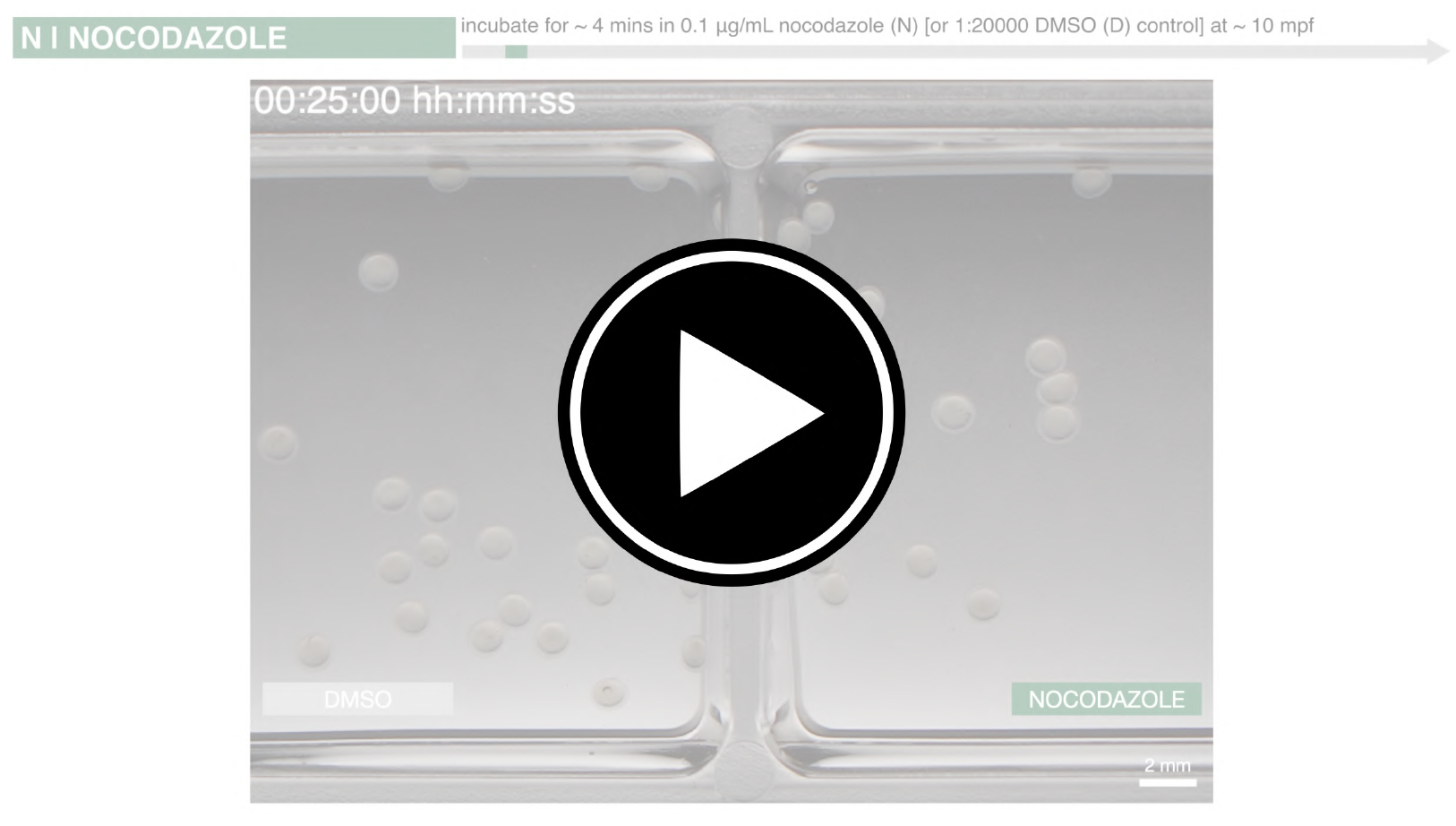
Rhythmic contractions of the yolk in goldfish embryos acutely treated with a microtubule depolymerizing drug, nocodazole. Simultaneous timelapse of goldfish embryos incubated in either 0.1 *µ*g/mL nocodazole (right) or DMSO control (left) for *∼* 4 mins at *∼* 10 mins post-fertilization. Time is in hh:mm:ss post-fertilization. Scale bar = 2 mm. Movie is available at https://youtu.be/Vnox05QuRJA.

**Fig. M9.**
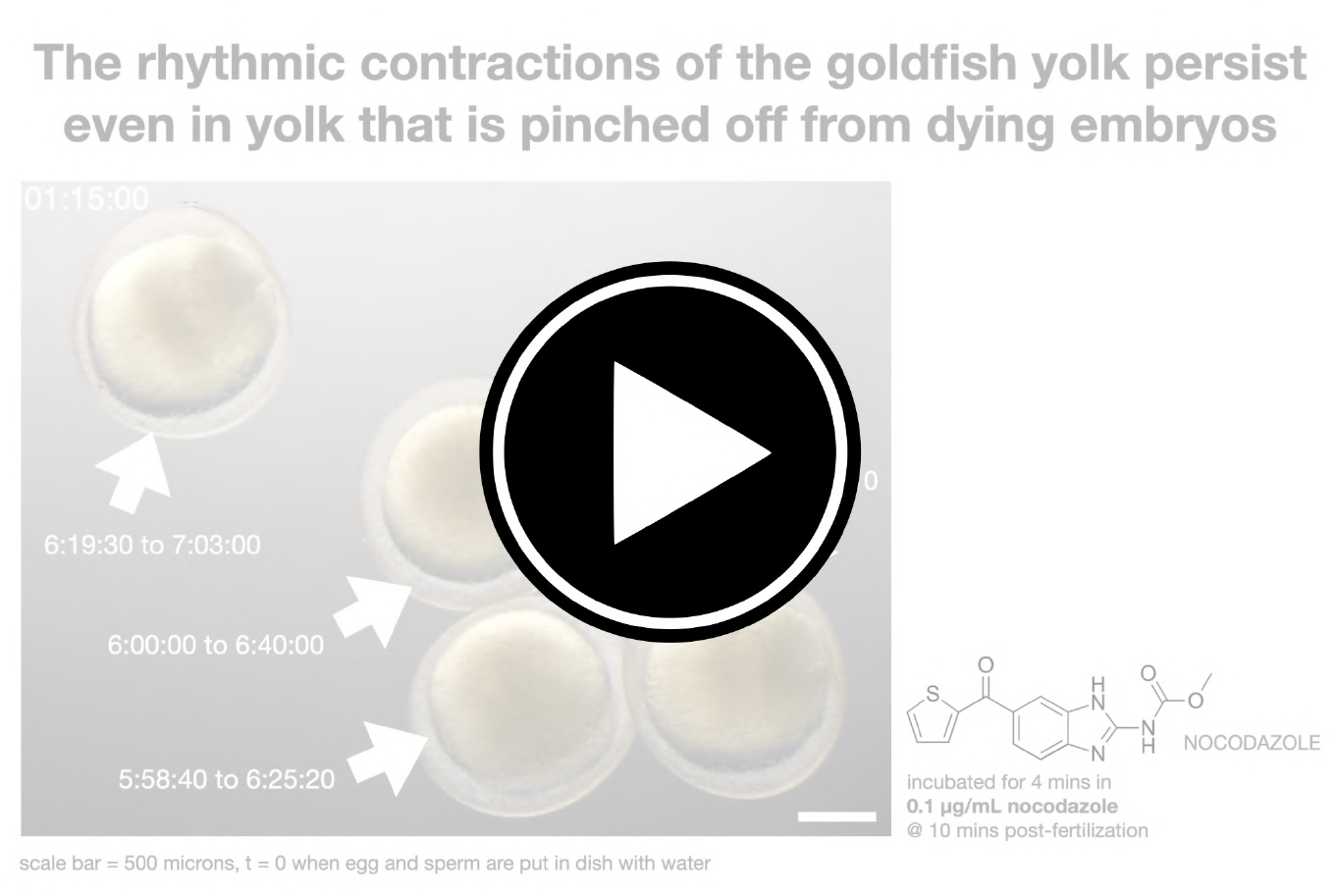
The rhythmic contractions of the goldfish yolk persist even in yolk that is pinched off from dying embryos. Timelapse of goldfish embryos showing persistent contractions of yolk that is pinched off from embryo. Embryos were treated with 0.1 *µ*g/mL nocodazole for *∼* 4 mins at *∼* 10 mins post-fertilization. White arrows mark samples showing persistent yolk contractions of pinched-off yolk. Specified numbers are representative timepoints when these contractions are visible. Time is in hh:mm:ss post-fertilization. Scale bar = 500 *µ*m. Movie is available at https://youtu.be/Fm296U5thaM.

**Fig. M10.**
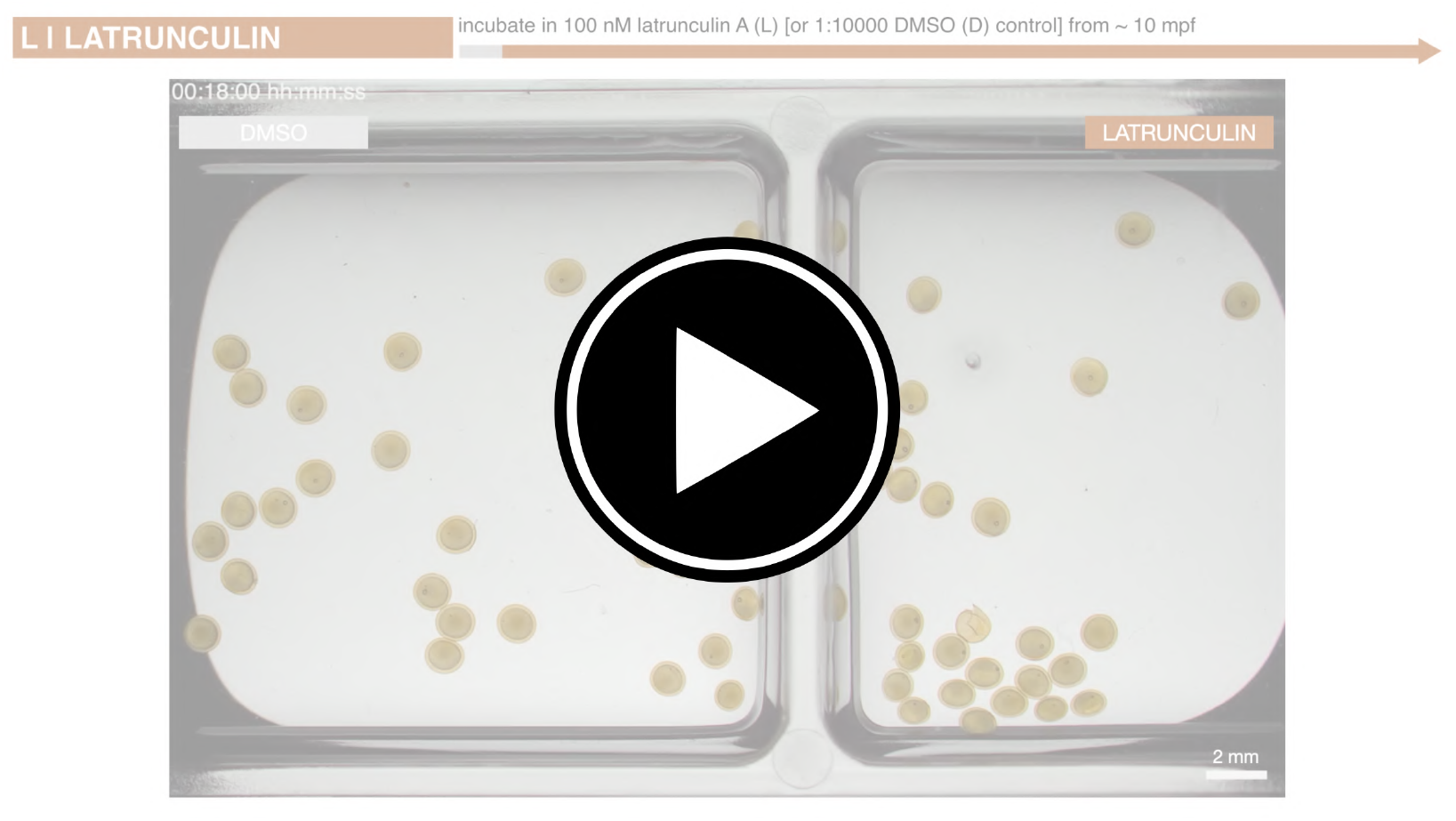
Rhythmic contractions of the yolk in goldfish embryos treated with an actin polymerization inhibitor, latrunculin. Simultaneous timelapse of goldfish embryos incubated in either 100 nM latrunculin A (right) or DMSO control (left) from *∼* 10 mins post-fertilization. Time is in hh:mm:ss post-fertilization. Scale bar = 2 mm. Movie is available at https://youtu.be/1-MOZORp3z0.

**Fig. M11.**
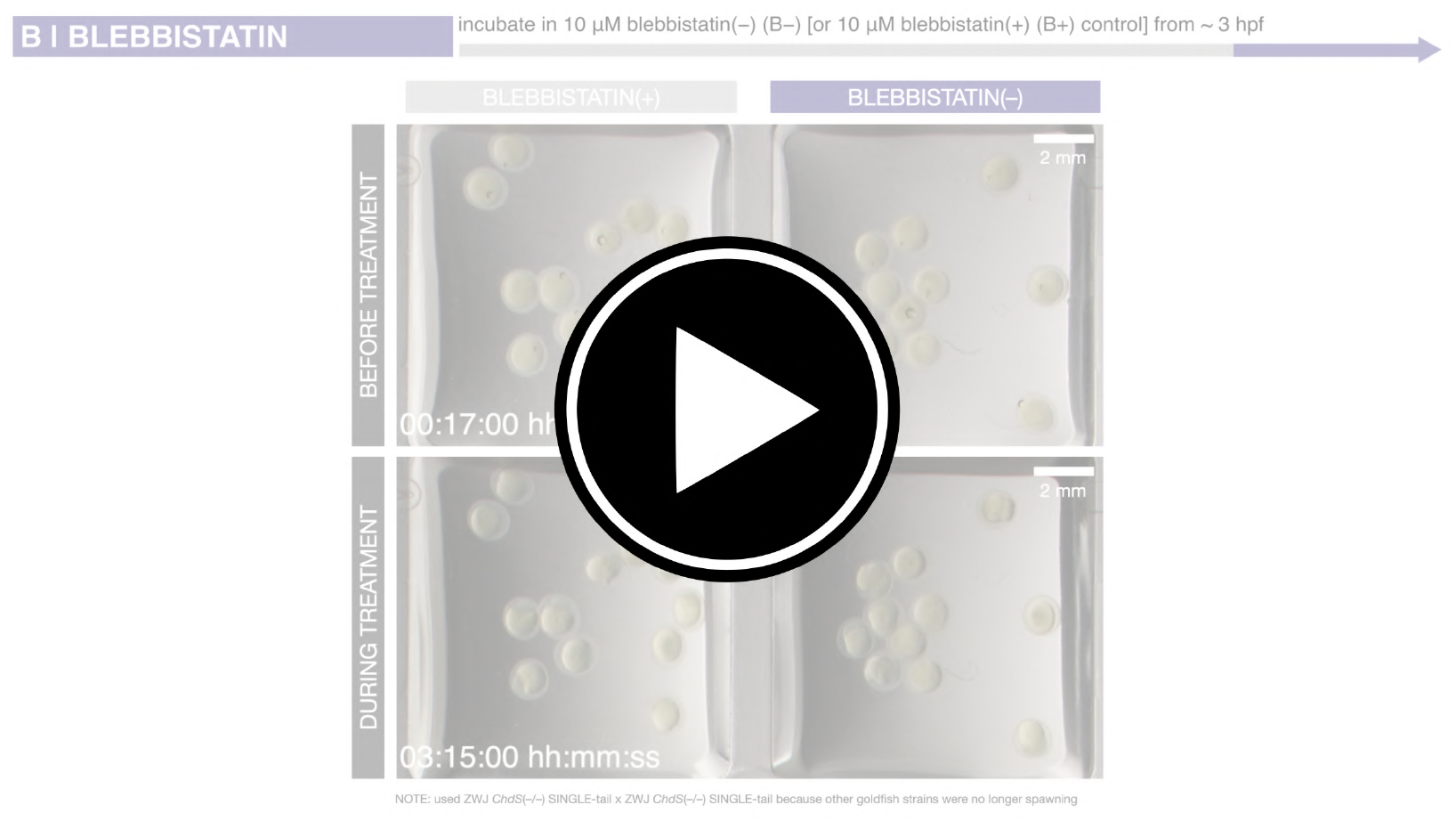
Rhythmic contractions of the yolk in goldfish embryos treated with a myosin inhibitor, blebbistatin(–). Simultaneous timelapse of goldfish embryos incubated in either 10 *µ*M blebbistatin(–) (right) or 10 *µ*M blebbistatin(+) control (left) from *∼* 3 hours post-fertilization. Top row: Simultaneous timelapse until 03:00:00 hh:mm:ss post-fertilization, before treatment. Bottom row: Simultaneous timelapse from 03:15:00 hh:mm:ss post-fertilization, during treatment. Time is in hh:mm:ss post-fertilization. Scale bar = 2 mm. Due to unavailability of spawning wild-type (ZWJ-*ChdS*(+/+) strain) goldfish, ZWJ-*ChdS*(–/–)-singletail goldfish was used. Movie is available at https://youtu.be/ZkxuXLxUj *A*.

**Fig. M12.**
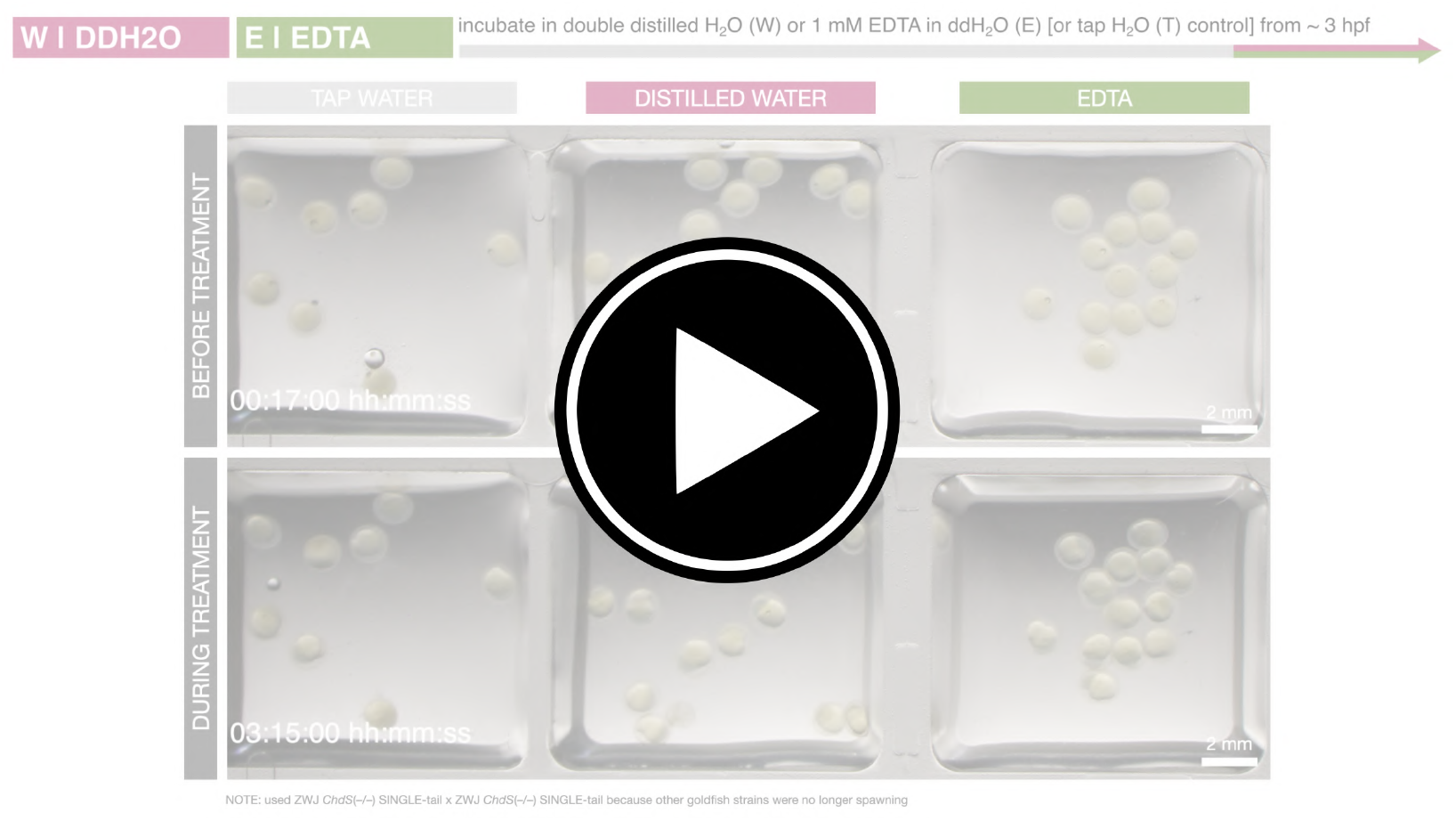
Rhythmic contractions of the yolk in goldfish embryos treated with Ca^2+^ free water or Ca^2+^ chelating agent, EDTA. Simultaneous timelapse of goldfish embryos incubated in either double distilled water (ddH_2_O, middle), 1 mM EDTA in ddH_2_O (right), or tap water control (left) from *∼* 3 hours post-fertilization. Top row: Simultaneous timelapse until 03:00:00 hh:mm:ss post-fertilization, before treatment. Bottom row: Simultaneous timelapse from 03:15:00 hh:mm:ss post-fertilization, during treatment. Time is in hh:mm:ss post-fertilization. Scale bar = 2 mm. Due to unavailability of spawning wild-type (ZWJ-*ChdS*(+/+) strain) goldfish, ZWJ-*ChdS*(–/–)-singletail goldfish was used. Movie is available at https://youtu.be/Ar1B1rbL6L0.

**Fig. M13.**
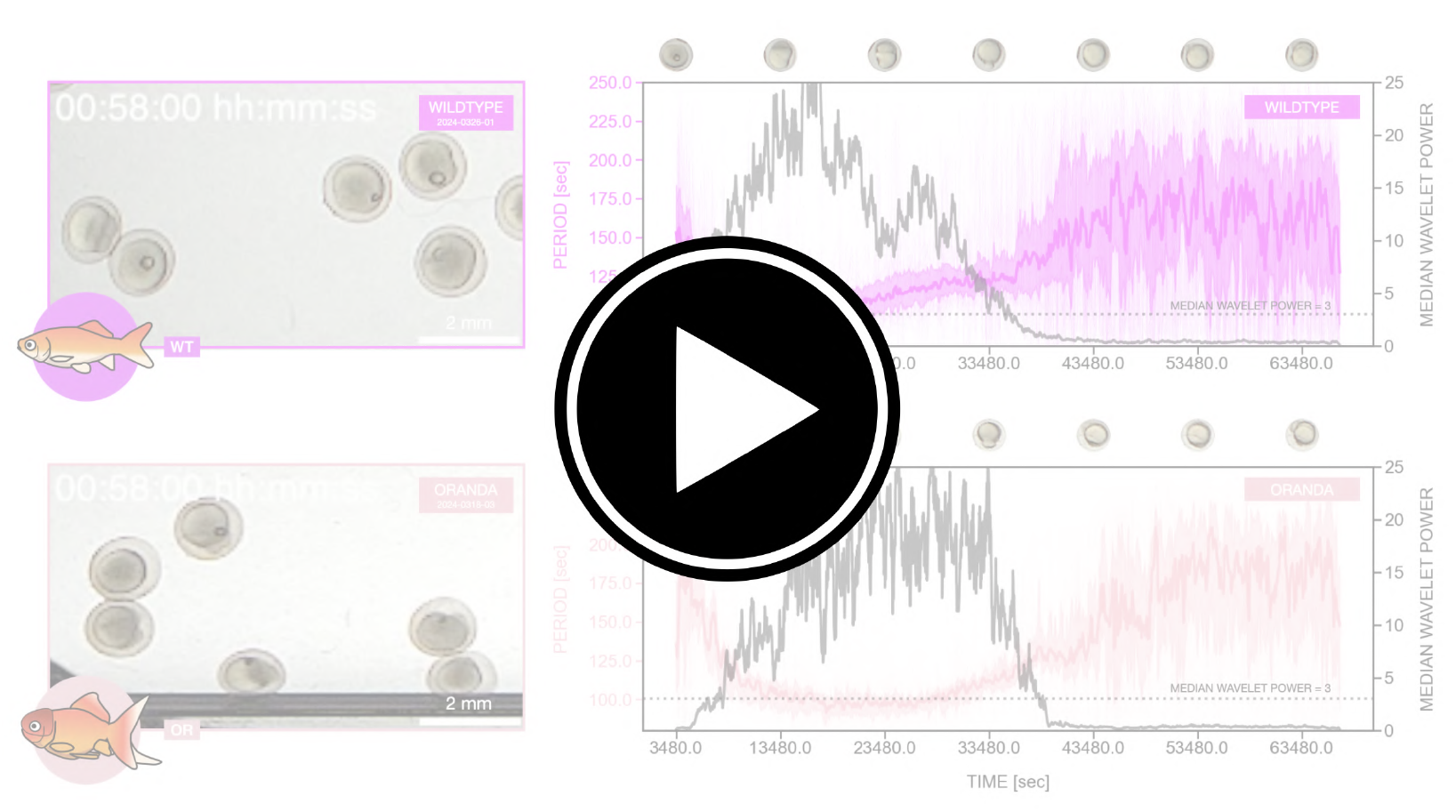
Rhythmic contractions of the yolk in twin-tail goldfish embryos are faster than those in wild-type. Left: Timelapse of wild-type (top) and twin-tail (i.e. *Oranda*, bottom) goldfish embryos. Time is in hh:mm:ss post-fertilization. Scale bar = 2 mm. Simultaneous imaging was unfortunately not possible because the two goldfish strains were not spawning at the same time. Right: Temporal evolution of period of yolk contractions for the entire duration of the experiment, obtained from wavelet analysis of detrended timeseries (via sinc-filter detrending, cut-off period = 250 seconds). The period evolution for each sample and the median of the periods are represented as a color-coded dashed line and a color-coded solid line, respectively. The color-coded shaded area corresponds to the interquartile range. Snapshots of representative fish embryos are shown above each plot. Median wavelet power is also plotted in gray, with a horizontal dotted gray line marking wavelet power threshold = 3, which corresponds to 95% confidence interval in case of white noise. Note shorter (i.e. faster) period of yolk contractions in twin-tail *Oranda* goldfish embryos. These data are the same as those plotted in Figure 6C. Movie is available at https://youtu.be/XZ7K8HKpS0Y.

**Fig. M14.**
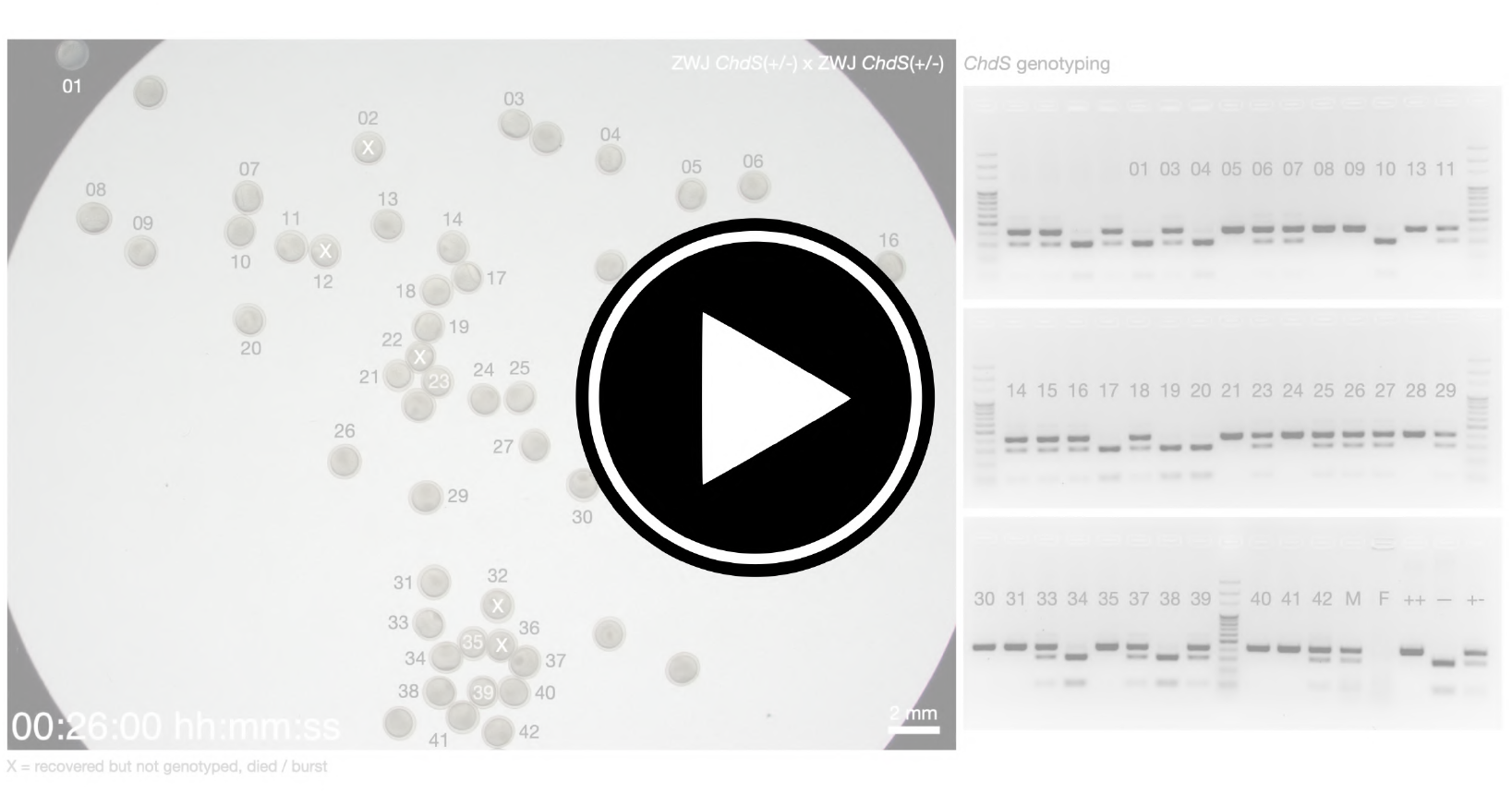
Rhythmic contractions of the yolk in goldfish embryos with different *ChdS* genotypes. Left: Simultaneous timelapse of goldfish embryos from mating of ZWJ-*ChdS*(+/–) goldfish. Numbers denote sample ID for genotyping and X marks samples that were recovered but were not genotyped (due to loss during handling). Time is in hh:mm:ss post-fertilization. Scale bar = 2 mm. Right: *ChdS* genotyping of samples in the left. Embryos were recovered at 1 day post-fertilization, after timelapse imaging, and were subsequently genotyped from 3 days post-fertilization. For genotyping, *∼* 400 bp of the *ChdS* gene was amplified and then treated with the Eco88I restriction enzyme, which recognizes a restriction enzyme site in the *ChdS*(–) allele resulting in a shorter *∼* 300 bp (and *∼* 100 bp) band (Abe et al.,2014). Movie is available at https://youtu.be/nVYwIuxD964.

**Fig. M15.**
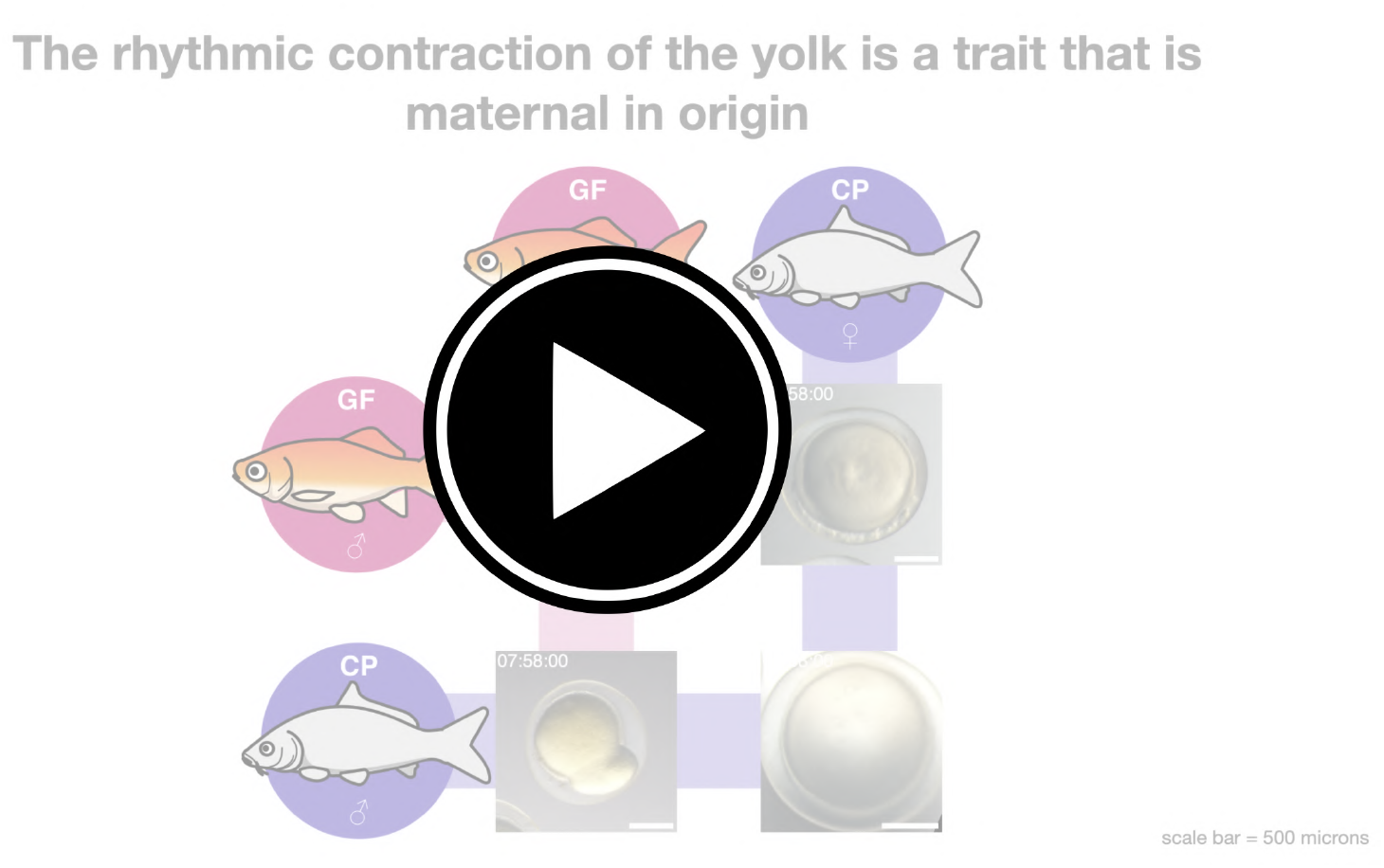
The rhythmic contraction of the yolk is a trait that is maternal in origin. Timelapse of embryos of GF x GF goldfish, CP x CP common carp, and their hybrids: (1) CP x GF from common carp sperm and goldfish egg and (2) GF x CP from goldfish sperm and common carp egg. Time is in hh:mm:ss post-fertilization. Scale bar = 500 *µ*m. Samples are the same as those in Figure 8B. Note rhythmic contractions of the yolk in embryos where the egg had been derived from goldfish. Movie is available at https://youtu.be/8l5PfF2uOjE.

## Bibliography

Abe, G., Lee, S.-H., Chang, M., Liu, S.-C., Tsai, H.-Y., and Ota, K. G. (2014). The origin of the bifurcated axial skeletal system in the twin-tail goldfish. Nature communications, 5(1):3360.

Abe, G., Lee, S.-H., Li, I.-J., Chang, C.-J., Tamura, K., and Ota, K. G. (2016). Open and closed evolutionary paths for drastic morphological changes, involving serial gene duplication, sub-functionalization and selection. Scientific reports, 6(1):26838.

Bénazéraf, B. and Pourquié, O. (2013). Formation and segmentation of the vertebrate body axis. Annual review of cell and developmental biology, 29:1–26.

Brownlee, C. and Dale, B. (1990). Temporal and spatial correlation of fertilization current, calcium waves and cytoplasmic contraction in eggs of ciona intestinalis. Proceedings of the Royal Society of London. B. Biological Sciences, 239(1296):321–328.

Burkel, B. M., Von Dassow, G., and Bement, W. M. (2007). Versatile fluorescent probes for actin filaments based on the actin-binding domain of utrophin. Cell motility and the cytoskeleton, 64(11):822–832.

Byrnes, W. M. and Newman, S. A. (2014). Ernest everett just: Egg and embryo as excitable systems. Journal of Experimental Zoology Part B: Molecular and Developmental Evolution, 322(4):191–201.

Cai, N. (1989). Studies on the formation of single caudal fin of the gold fish, carassius auratus – i. the effect of the cytoplasm on the development of caudal. Oceanologia et Limnologia Sinica, 20(5):453–459.

Cartwright, J. H., Piro, O., and Tuval, I. (2009). Fluid dynamics in developmental biology: moving fluids that shape ontogeny. HFSP journal, 3(2):77–93.

Chen, H.-C., Wang, C., Li, I.-J., Abe, G., and Ota, K. G. (2022). Pleiotropic functions of chordin gene causing drastic morphological changes in ornamental goldfish. Scientific Reports, 12(1):19961.

Chen, J., Xia, L., Bruchas, M. R., and Solnica-Krezel, L. (2017). Imaging early embryonic calcium activity with gcamp6s transgenic zebrafish. Developmental biology, 430(2):385–396.

Chen, T.-W., Wardill, T. J., Sun, Y., Pulver, S. R., Renninger, S. L., Baohan, A., Schreiter, E. R., Kerr, R. A., Orger, M. B., Jayaraman, V., et al. (2013). Ultrasensitive fluorescent proteins for imaging neuronal activity. Nature, 499(7458):295–300.

Cooke, J. (1988). The early embryo and the formation of body pattern. American Scientist, 76(1):35–41.

Crozier, W. J. (1924). On biological oxidations as function of temperature. The Journal of General Physiology, 7(2):189–216.

Crozier, W. J. (1926). On curves of growth, especially in relation to temperature. The Journal of General Physiology, 10(1):53–73.

Deguchi, R., Shirakawa, H., Oda, S., Mohri, T., and Miyazaki, S. (2000). Spatiotemporal analysis of ca2+ waves in relation to the sperm entry site and animal–vegetal axis during ca2+ oscillations in fertilized mouse eggs. Developmental biology, 218(2):299–313.

Deneke, V. E. and Di Talia, S. (2018). Chemical waves in cell and developmental biology. Journal of Cell Biology, 217(4):1193–1204.

Devillers, C. (1961). Structural and dynamic aspects of the development of the teleostean egg. In Advances in morphogenesis, volume 1, pages 379–428. Elsevier.

Di Talia, S. and Vergassola, M. (2022). Waves in embryonic development. Annual review of biophysics, 51:327–353.

Foster, P. J., Fürthauer, S., and Fakhri, N. (2022). Active mechanics of sea star oocytes. bioRxiv, pages 2022–04.

Friedland, G., Jantz, K., Lenz, T., and Rojas, R. (2006). Extending the siox algorithm: alternative clustering methods, sub-pixel accurate object extraction from still images, and generic video segmentation.

Ge, X., Grotjahn, D., Welch, E., Lyman-Gingerich, J., Holguin, C., Dimitrova, E., Abrams, E. W., Gupta, T., Marlow, F. L., Yabe, T., et al. (2014). Hecate/grip2a acts to reorganize the cytoskeleton in the symmetry-breaking event of embryonic axis induction. PLoS genetics, 10(6):e1004422.

Gilland, E., Miller, A. L., Karplus, E., Baker, R., and Webb, S. E. (1999). Imaging of multicellular large-scale rhythmic calcium waves during zebrafish gastrulation. Proceedings of the National Academy of Sciences, 96(1):157–161.

Gingerich, J. L., Westfall, T. A., Slusarski, D. C., and Pelegri, F. (2005). Hecate, a zebrafish maternal effect gene, affects dorsal organizer induction and intracellular calcium transient frequency. Developmental biology, 286(2):427–439.

Gnaiger, E. (2021). Beyond counting papers–a mission and vision for scientific publication. Bioenergetics Communications, 2021:5–5.

Goedhart, J. (2021). Superplotsofdata—a web app for the transparent display and quantitative comparison of continuous data from different conditions. Molecular biology of the cell, 32(6):470–474.

Goodwin, B. C. and Cohen, M. H. (1969). A phase-shift model for the spatial and temporal organization of developing systems. Journal of Theoretical Biology, 25(1):49–107.

Gore, A. V., Maegawa, S., Cheong, A., Gilligan, P. C., Weinberg, E. S., and Sampath, K. (2005). The zebrafish dorsal axis is apparent at the four-cell stage. Nature, 438(7070):1030–1035.

Gore, A. V. and Sampath, K. (2002). Localization of transcripts of the zebrafish morphogen squint is dependent on egg activation and the microtubule cytoskeleton. Mechanisms of development, 112(1-2):153–156.

Goutel, C., Kishimoto, Y., Schulte-Merker, S., and Rosa, F. (2000). The ventralizing activity of radar, a maternally expressed bone morphogenetic protein, reveals complex bone morphogenetic protein interactions controlling dorso-ventral patterning in zebrafish. Mechanisms of development, 99(1-2):15–27.

Grimes, D. T. and Burdine, R. D. (2017). Left–right patterning: breaking symmetry to asymmetric morphogenesis. Trends in Genetics, 33(9):616–628.

Guevorkian, K. and Maître, J.-L. (2017). Micropipette aspiration: A unique tool for exploring cell and tissue mechanics in vivo. In Methods in cell biology, volume 139, pages 187–201. Elsevier.

Halet, G., Tunwell, R., Parkinson, S. J., and Carroll, J. (2004). Conventional pkcs regulate the temporal pattern of ca2+ oscillations at fertilization in mouse eggs. The Journal of cell biology, 164(7):1033–1044.

Harris, C. R., Millman, K. J., Van Der Walt, S. J., Gommers, R., Virtanen, P., Cournapeau, D., Wieser, E., Taylor, J., Berg, S., Smith, N. J., et al. (2020). Array programming with numpy. Nature, 585(7825):357–362.

Hibi, M., Hirano, T., and Dawid, I. B. (2002). Organizer formation and function. Pattern Formation in Zebrafish, pages 48–71.

Hibi, M., Takeuchi, M., Hashimoto, H., and Shimizu, T. (2018). Axis formation and its evolution in ray-finned fish. Reproductive and Developmental Strategies: The Continuity of Life, pages 709–742.

Hoebeke, J., Van Nijen, G., and De Brabander, M. (1976). Interaction of oncodazole (r 17934), a new anti-tumoral drug, with rat brain tubulin. Biochemical and biophysical research communications, 69(2):319–324.

Hubbard, M. and Rothschild, L. W. (1939). Spontaneous rhythmical impedance changes in the trout’s egg. Proceedings of the Royal Society of London. Series B-Biological Sciences, 127(849):510–526.

Hunter, J. D. (2007). Matplotlib: A 2d graphics environment. Computing in science & engineering, 9(03):90–95.

Ishii, H. and Tani, T. (2021). Dynamic organization of cortical actin filaments during the ooplasmic segregation of ascidian ciona eggs. Molecular Biology of the Cell, 32(3):274–288.

Jesuthasan, S. and Strähle, U. (1997). Dynamic microtubules and specification of the zebrafish embryonic axis. Current Biology, 7(1):31–42.

Just, E. (1919). The fertilization reaction in echinarachnius parma: I. cortical response of the egg to insemination. The Biological Bulletin, 36(1):1–10.

Just, E. E. (1939). The biology of the cell surface. P. Blakiston’s Son and Co., Inc., Philadelphia, USA.

Kasansky, W. J. (1936). Verschiedene bewegungsarten des hechtei-inhaltes (esox lucius l.) in den friihen stadien der embryonalen entwicklung. Zoologischer Anzeiger, 115(1/2):89–96.

Kosuta, C., Daniel, K., Johnstone, D. L., Mongeon, K., Ban, K., LeBlanc, S., MacLeod, S., Et-Tahiry, K., Ekker, M., MacKenzie, A., et al. (2018). High-throughput dna extraction and genotyping of 3dpf zebrafish larvae by fin clipping. Journal of Visualized Experiments: JoVE, (136).

Kyozuka, K., Chun, J. T., Puppo, A., Gragnaniello, G., Garante, E., and Santella, L. (2008). Actin cytoskeleton modulates calcium signaling during maturation of starfish oocytes. Developmental biology, 320(2):426–435.

Langdon, Y. G. and Mullins, M. C. (2011). Maternal and zygotic control of zebrafish dorsoventral axial patterning. Annual review of genetics, 45:357–377.

Lee, K. W., Webb, S. E., and Miller, A. L. (1999). A wave of free cytosolic calcium traverses zebrafish eggs on activation. Developmental biology, 214(1):168–180.

Lee, S.-H., Wang, C. Y., Li, J., Abe, G., and Ota, K. (2023). Competition or contingency? using crispr/cas9-induced mutants to examine the potential origin of a unique mutant allele in twintail goldfish. Research Square.

Lereboullet, A. (1862). Recherches d’embryologie comparée sur le développement du brochet, de la perche et de l’écrevisse. Impr. impériale.

Lim, S., Kumari, P., Gilligan, P., Quach, H. N. B., Mathavan, S., and Sampath, K. (2012). Dorsal activity of maternal squint is mediated by a non-coding function of the rna. Development, 139(16):2903–2915.

Limatola, N., Chun, J. T., and Santella, L. (2022a). Regulation of the actin cytoskeleton-linked ca2+ signaling by intracellular ph in fertilized eggs of sea urchin. Cells, 11(9):1496.

Limatola, N., Chun, J. T., and Santella, L. (2022b). Species-specific gamete interaction during sea urchin fertilization: Roles of the egg jelly and vitelline layer. Cells, 11(19):2984.

Litschel, T., Kelley, C. F., Holz, D., Adeli Koudehi, M., Vogel, S. K., Burbaum, L., Mizuno, N., Vavylonis, D., and Schwille, P. (2021). Reconstitution of contractile actomyosin rings in vesicles. Nature communications, 12(1):2254.

Ma, J. and Carney, T. J. (2024). Protease-activated receptor 2 links protease activity with calcium waves during egg activation and blastomere cleavage. bioRxiv, pages 2024–05.

Maître, J.-L., Niwayama, R., Turlier, H., Nédélec, F., and Hiiragi, T. (2015). Pulsatile cellautonomous contractility drives compaction in the mouse embryo. Nature cell biology, 17(7):849–855.

Matsui, Y. (1935). Kagaku to shumi kara mita kingyo no kenkyu .

McKinney, W. (2010). Data structures for statistical computing in python. 56–61. In Proc 9th Python Sci Conf (SCIPY 2010).

Meinhardt, H. (2006). Primary body axes of vertebrates: Generation of a near-cartesian coordinate system and the role of spemann-type organizer. Developmental Dynamics, 235(11):2907–2919.

Minin, A. and Ozerova, S. (2008). Spontaneous activation of fish eggs is abolished by protease inhibitors. Russian Journal of Developmental Biology, 39:293–296.

Miyazaki, M., Chiba, M., Eguchi, H., Ohki, T., and Ishiwata, S. (2015). Cell-sized spherical confinement induces the spontaneous formation of contractile actomyosin rings in vitro. Nature cell biology, 17(4):480–489.

Mizuno, T., Yamaha, E., Kuroiwa, A., and Takeda, H. (1999). Removal of vegetal yolk causes dorsal deficencies and impairs dorsal-inducing ability of the yolk cell in zebrafish. Mechanisms of development, 81(1-2):51–63.

Mizuno, T., Yamaha, E., and Yamazaki, F. (1997). Localized axis determinant in the early cleavage embryo of the goldfish, carassius auratus. Development Genes and Evolution, 206:389–396.

Mohri, T. and Kyozuka, K. (2022). Starfish oocytes of a. pectinifera reveal marked differences in sperm-induced electrical and intracellular calcium changes during oocyte maturation and at fertilization. Molecular Reproduction and Development, 89(1):3–22.

Mönke, G., Sorgenfrei, F. A., Schmal, C., and Granada, A. E. (2020). Optimal time frequency analysis for biological data-pyboat. BioRxiv, pages 2020–04.

Newman, S. A. (2009). Ee just’s “independent irritability” revisited: the activated egg as excitable soft matter. Molecular Reproduction and Development: Incorporating Gamete Research, 76(10):966–974.

Nojima, H., Rothhämel, S., Shimizu, T., Kim, C.-H., Yonemura, S., Marlow, F. L., and Hibi, M. (2010). Syntabulin, a motor protein linker, controls dorsal determination. Development, 137(6):923–933.

Ober, E. A. and Schulte-Merker, S. (1999). Signals from the yolk cell induce mesoderm, neuroectoderm, the trunk organizer, and the notochord in zebrafish. Developmental biology, 215(2):167–181.

Oppenheimer, J. M. (1936). The development of isolated blastoderms of fundulus heteroclitus. Journal of Experimental Zoology, 72(2):247–269.

Ota, K. G. (2021). Goldfish development and evolution. Springer.

Ota, K. G. and Abe, G. (2016). Goldfish morphology as a model for evolutionary developmental biology. Wiley Interdisciplinary Reviews: Developmental Biology, 5(3):272–295.

Özgüç, Ö. (2021). Mechanical and molecular regulation of periodic cortical waves of contraction. PhD thesis, Sorbonne Université.

Özgüç, Ö., de Plater, L., Kapoor, V., Tortorelli, A. F., Clark, A. G., and Maître, J.-L. (2022). Cortical softening elicits zygotic contractility during mouse preimplantation development. PLoS Biology, 20(3):e3001593.

Pereyra, M., Drusko, A., Krämer, F., Strobl, F., Stelzer, E. H., and Matthäus, F. (2021). Quickpiv: Efficient 3d particle image velocimetry software applied to quantifying cellular migration during embryogenesis. BMC bioinformatics, 22:1–20.

Pereyra, M., Golden, M., Lange, Z., Golden, A., Strobl, F., Stelzer, E. H., and Matthaeus, F. (2024). Spatio-temporal segmentation of contraction waves in the extra-embryonic membranes of the red flour beetle. bioRxiv, pages 2024–08.

Piccolo, S., Sasai, Y., Lu, B., and De Robertis, E. M. (1996). Dorsoventral patterning in xenopus: inhibition of ventral signals by direct binding of chordin to bmp-4. Cell, 86(4):589–598.

Postma, M. and Goedhart, J. (2019). Plotsofdata—a web app for visualizing data together with their summaries. PLoS biology, 17(3):e3000202.

Pourquié, O. (2011). Vertebrate segmentation: from cyclic gene networks to scoliosis. Cell, 145(5):650–663.

Ransom, W. H. (1867). Xiv. observations on the ovum of osseous fishes. Philosophical Transactions of the Royal Society of London, (157):431–501.

Rissi, M., Wittbrodt, J., Délot, E., Naegeli, M., and Rosa, F. M. (1995). Zebrafish radar: A new member of the tgf-β superfamily defines dorsal regions of the neural plate and the embryonic retina. Mechanisms of development, 49(3):223–234.

Roegiers, F., McDougall, A., and Sardet, C. (1995). The sperm entry point defines the orientation of the calcium-induced contraction wave that directs the first phase of cytoplasmic reorganization in the ascidian egg. Development, 121(10):3457–3466.

Rothschild, L. (1940). Rhythmical impedance changes in the trout’s egg. Nature, 145(3680):744– 744.

Rothschild, L. (1947). Spontaneous rhythmical impedance changes in the egg of the trout. ii. Journal of Experimental Biology, 23(3-4):267–276.

Rusconi, M. (1840). Ueber künstliche befruchtung von fischen ng über einige neue versuche in betrelf künstlicher beftuchtung an fröschen. Archiv für Anatomie, Physiologie, und Wissenschaftliche Medicin, pages 185–193.

Sakamoto, R., Izri, Z., Shimamoto, Y., Miyazaki, M., and Maeda, Y. T. (2022). Geometric trade-off between contractile force and viscous drag determines the actomyosin-based motility of a cell-sized droplet. Proceedings of the National Academy of Sciences, 119(30):e2121147119.

Sakamoto, R., Tanabe, M., Hiraiwa, T., Suzuki, K., Ishiwata, S., Maeda, Y. T., and Miyazaki, M. (2020). Tug-of-war between actomyosin-driven antagonistic forces determines the positioning symmetry in cell-sized confinement. Nature communications, 11(1):3063.

Salbreux, G., Joanny, J.-F., Prost, J., and Pullarkat, P. (2007). Shape oscillations of non-adhering fibroblast cells. Physical biology, 4(4):268.

Santella, L. and Chun, J. T. (2022). Structural actin dynamics during oocyte maturation and fertilization. Biochemical and Biophysical Research Communications, 633:13–16.

Sardet, C., Paix, A., Prodon, F., Dru, P., and Chenevert, J. (2007). From oocyte to 16-cell stage: cytoplasmic and cortical reorganizations that pattern the ascidian embryo. Developmental Dynamics: An Official Publication of the American Association of Anatomists, 236(7):1716– 1731.

Sardet, C., Roegiers, F., Dumollard, R., Rouviere, C., and McDougall, A. (1998). Calcium waves and oscillations in eggs. Biophysical chemistry, 72(1-2):131–140.

Sasai, Y., Lu, B., Steinbeisser, H., Geissert, D., Gont, L. K., and De Robertis, E. M. (1994). Xenopus chordin: a novel dorsalizing factor activated by organizer-specific homeobox genes. Cell, 79(5):779–790.

Saunders, D. (2024). Prereview of “on the independent irritability of goldfish eggs and embryos – a living communication on the rhythmic yolk contractions in goldfish”. Zenodo.

Schindelin, J., Arganda-Carreras, I., Frise, E., Kaynig, V., Longair, M., Pietzsch, T., Preibisch, S., Rueden, C., Saalfeld, S., Schmid, B., et al. (2012). Fiji: an open-source platform for biological-image analysis. Nature methods, 9(7):676–682.

Sidi, S., Goutel, C., Peyriéras, N., and Rosa, F. M. (2003). Maternal induction of ventral fate by zebrafish radar. Proceedings of the National Academy of Sciences, 100(6):3315–3320.

Simon, J. Z. and Cooper, M. S. (1995). Calcium oscillations and calcium waves coordinate rhythmic contractile activity within the stellate cell layer of medaka fish embryos. Journal of Experimental Zoology, 273(2):118–129.

Spector, I., Shochet, N. R., Kashman, Y., and Groweiss, A. (1983). Latrunculins: novel marine toxins that disrupt microfilament organization in cultured cells. Science, 219(4584):493–495.

Straight, A. F., Cheung, A., Limouze, J., Chen, I., Westwood, N. J., Sellers, J. R., and Mitchison, T. J. (2003). Dissecting temporal and spatial control of cytokinesis with a myosin ii inhibitor. Science, 299(5613):1743–1747.

Stricker, S. A. (1999). Comparative biology of calcium signaling during fertilization and egg activation in animals. Developmental biology, 211(2):157–176.

Sun, Y.-H., Chen, S.-P., Wang, Y.-P., Hu, W., and Zhu, Z.-Y. (2005). Cytoplasmic impact on crossgenus cloned fish derived from transgenic common carp (cyprinus carpio) nuclei and goldfish (carassius auratus) enucleated eggs. Biology of reproduction, 72(3):510–515.

Tran, L. D., Hino, H., Quach, H., Lim, S., Shindo, A., Mimori-Kiyosue, Y., Mione, M., Ueno, N., Winkler, C., Hibi, M., et al. (2012). Dynamic microtubules at the vegetal cortex predict the embryonic axis in zebrafish. Development, 139(19):3644–3652.

Tsai, F.-C., Stuhrmann, B., and Koenderink, G. H. (2011). Encapsulation of active cytoskeletal protein networks in cell-sized liposomes. Langmuir, 27(16):10061–10071.

Tsai, H.-Y., Chang, M., Liu, S.-C., Abe, G., and Ota, K. G. (2013). Embryonic development of goldfish (carassius auratus): a model for the study of evolutionary change in developmental mechanisms by artificial selection. Developmental dynamics, 242(11):1262–1283.

Tsuruwaka, Y., Konishi, T., Miyawaki, A., and Takagi, M. (2007). Real-time monitoring of dynamic intracellular ca2+ movement during early embryogenesis through expression of yellow cameleon. Zebrafish, 4(4):253–260.

Tung, T.-C., Chang, C.-Y., and Tung, Y.-F.-Y. (1945). Experiments on the developmental potencies of blastoderms and fragments of teleostean eggs separated latitudinally. In Proceedings of the Zoological Society of London, volume 115, pages 175–188. Wiley Online Library.

Tung, T.-C., Lee, C.-Y., and Tung, Y.-F.-Y. (1955a). Further research on the developmental ability of fish eggs. Acta Biologiae Experimentalis Sinica, 2.

Tung, T.-C. and Tung, Y.-F.-Y. (1943). Experimental studies on the development of goldfish. In Proceedings of the Chinese Physiological Society.

Tung, T.-C. and Tung, Y.-F.-Y. (1944). The development of egg-fragments, isolated blastomeres and fused eggs in the goldfish. In Proceedings of the Zoological Society of London, volume 114, pages 46–64. Wiley Online Library.

Tung, T.-C., Wu, S.-Q., and Tung, Y.-F.-Y. (1955b). The development of the isolated fragments of carassius eggs centrifuged after fertilization. Acta Biologiae Experimentalis Sinica, 1.

Turing, A. M. (1952). The chemical basis of morphogenesis. Bulletin of mathematical biology, 52:153–197.

Uriu, K. (2016). Genetic oscillators in development. Development, Growth & Differentiation, 58(1):16–30.

Van Der Walt, S., Colbert, S. C., and Varoquaux, G. (2011). The numpy array: a structure for efficient numerical computation. Computing in science & engineering, 13(2):22–30.

Van der Walt, S., Schönberger, J. L., Nunez-Iglesias, J., Boulogne, F., Warner, J. D., Yager, N., Gouillart, E., and Yu, T. (2014). scikit-image: image processing in python. PeerJ, 2:e453.

Virtanen, P., Gommers, R., Oliphant, T. E., Haberland, M., Reddy, T., Cournapeau, D., Burovski, E., Peterson, P., Weckesser, W., Bright, J., et al. (2020). Scipy 1.0: fundamental algorithms for scientific computing in python. Nature methods, 17(3):261–272.

Waskom, M. L. (2021). Seaborn: statistical data visualization. Journal of Open Source Software, 6(60):3021.

Westfall, T. A., Hjertos, B., and Slusarski, D. C. (2003). Requirement for intracellular calcium modulation in zebrafish dorsal–ventral patterning. Developmental biology, 259(2):380–391.

Wintrebert, P. and Yung, K.-C. (1926). La contraction protoplasmique des ébauches embryonnaires chez l’Épinoche et l’Épinochette. Comptes rendus hebdomadaires des séances de l’Académie des sciences, 183:455–456.

Wülker, W. (1953). Bewegungsrhythmen im Teleostier - Ei (Zeitrafferfilm-Untersuchung). 1. Esox lucius, Salmo trutta, S.fontinalis, S.irideus, volume 73.

Yamamoto, T.-o. (1934). On the rhythmic movements of the egg of goldfish. J. Fac. Sci., Imp. Univ. Tokyo, 3(3):275–285.

Yamamoto, T.-o. (1938). Contractile movement of the egg of a bony fish, salanx microdon. Proceedings of the Imperial Academy, 14(4):149–151.

Yamamoto, T.-o. (1940). Rhythmical contractile movement of eggs of trouts. Annotationes Zoologicae Japonenses, 19(1):69–79.

Yamamoto, T.-o. (1954). Cortical changes in eggs of the goldfish (carassius auratus) and the pond smelt (hypomesus olidus) at the time of fertilization and activation. Japanese Journal of Ichthyology, 3(3-5):162–170.

Yu, P., Wang, Y., Li, Z., Jin, H., Li, L.-L., Han, X., Wang, Z.-W., Yang, X.-L., Li, X.-Y., Zhang, X.-J., et al. (2022). Causal gene identification and desirable trait recreation in goldfish. Science China Life Sciences, 65(12):2341–2353.

Zhang, Y., Xia, H., Li, S., Liu, J., Liu, L., and Yang, P. (2023). Early embryonic development of green crucian carp carassius auratus indigentiaus subsp. nov.

